# Brain kernel: a new spatial covariance function for fMRI data

**DOI:** 10.1101/2021.03.22.436524

**Authors:** Anqi Wu, Samuel A. Nastase, Christopher A. Baldassano, Nicholas B. Turk-Browne, Kenneth A. Norman, Barbara E. Engelhardt, Jonathan W. Pillow

**Affiliations:** Center for Theoretical Neuroscience, Columbia University, New York City, NY, USA; Princeton Neuroscience Institute, Princeton University, Princeton, NJ, USA; Department of Psychology, Princeton University, Princeton, NJ, USA; Department of Psychology, Columbia University, New York City, NY, USA; Department of Psychology, Yale University, New Haven, CT, USA; Department of Computer Science, Princeton University, Princeton, NJ, USA

**Keywords:** Brain kernel, Gaussian process, latent variable model, brain decoding, factor modeling, resting-state fMRI, task fMRI

## Abstract

A key problem in functional magnetic resonance imaging (fMRI) is to estimate spatial activity patterns from noisy high-dimensional signals. Spatial smoothing provides one approach to regularizing such estimates. However, standard smoothing methods ignore the fact that correlations in neural activity may fall off at different rates in different brain areas, or exhibit discontinuities across anatomical or functional boundaries. Moreover, such methods do not exploit the fact that widely separated brain regions may exhibit strong correlations due to bilateral symmetry or the network organization of brain regions. To capture this non-stationary spatial correlation structure, we introduce the *brain kernel*, a continuous covariance function for whole-brain activity patterns. We define the brain kernel in terms of a continuous nonlinear mapping from 3D brain coordinates to a latent embedding space, parametrized with a Gaussian process (GP). The brain kernel specifies the prior covariance between voxels as a function of the distance between their locations in embedding space. The GP mapping warps the brain nonlinearly so that highly correlated voxels are close together in latent space, and uncorrelated voxels are far apart. We estimate the brain kernel using resting-state fMRI data, and we develop an exact, scalable inference method based on block coordinate descent to overcome the challenges of high dimensionality (10-100K voxels). Finally, we illustrate the brain kernel’s usefulness with applications to brain decoding and factor analysis with multiple task-based fMRI datasets.

## 1 Introduction

An important problem in neuroscience is to characterize the covariance of high-dimensional neural activity. Understanding this covariance structure could provide insight into the brain’s functional organization and help regularize estimates of encoding or decoding models. Although advances have been made in both theory and methodology for estimating large covariance [1, 2] and precision matrices [3, 4], few methods have been designed with the particular challenges of functional magnetic resonance imaging (fMRI) data in mind (see [5] for an exception).

One of the challenges of modeling the covariance of fMRI data is that the spatial discretization of the brain may differ across experiments. FMRI measures blood oxygenation level dependent (BOLD) signals in discrete spatial regions called “voxels”. Each voxel represents a tiny cube of brain tissue. Although brains are typically registered to an anatomical template in a standard space, such as the volumetric Montreal Neurological Institute (MNI) template or the surface-based template used by HCP [6], these spaces differ in resolution and geometry [7]. In many cases, brains are aligned onto the “same” 3D space but with different voxel coordinates. A covariance matrix for one set of voxels cannot be applied to data registered to a different set of voxels. Thus, modeling the covariance of fMRI data presently requires a new covariance matrix to be constructed whenever a different set of voxels is used.

A second challenge for fMRI covariance estimation is spatial nonstationarity. Standard spatial smoothing models assume that correlation falls off as a function of the Euclidean distance between voxels. In real brains, however, correlation patterns depend on relationships to anatomical and functional boundaries, and may exhibit strong dependencies over long distances due to bilateral symmetry and the network organization of brain regions.

To address these challenges, we propose the *brain kernel*, a continuous covariance function for whole-brain fMRI data. This function arises from a generative model of fMRI data, and seeks to describe covariance of neural signals across the entire brain [8]. Specifically, the brain kernel defines, for any finite set of *n* voxel locations in the brain, a positive definite *n* × *n* covariance matrix over *n*-dimensional vectors of neural activity at those locations. The brain kernel improves upon prior work by i) capturing fMRI voxel covariance matrices for any registration reference, and ii) capturing spatial nonstationarity through a nonlinear latent manifold.

Our approach uses a Gaussian process to parametrize a continuous nonlinear mapping from 3D brain coordinates to a latent embedding space, such that correlations in neural activity fall off as a fixed function of distance in the latent space. Thus, the nonlinear function seeks to warp the 3D brain in order to place locations with correlated neural activity at nearby locations in the latent space. Locations with uncorrelated activity, conversely, are mapped to more distant points in the latent space, even if they are physically close together in the brain.

The paper is organized as follows. In Sec. 2 we provide a brief overview of Gaussian process models. In Sec. 3, we formally introduce the brain kernel model for fMRI data. In Sec. 4, we describe an efficient inference method for fitting the brain kernel, and illustrate the challenges and benefits of an exact inference method using simulated and real fMRI datasets. In Sec. 5, we describe the brain kernel fit to whole-brain resting-state fMRI data. Finally, in Sec. 6, we demonstrate the usefulness of the inferred brain kernel with applications to decoding and factor modeling.

## 2 Mathematical background

Before introducing the brain kernel model, we briefly review the mathematical building blocks for Gaussian process models.

### 2.1 Gaussian processes (GPs)

Gaussian processes provide a flexible and tractable prior distribution over nonlinear functions [9]. A GP is parametrized by a mean function *m*(x), which specifies the mean value of the function *f*(x) at input point x, and a covariance function *k*(x_1_, x_2_), which specifies cov(*f*(x_1_),*f*(x_2_)), the covariance between the function values *f*(x_1_) and *f*(x_2_), for any pair of inputs x_1_ and x_2_.

Technically a GP is a random process for which the values taken at any finite set of input points has a well-defined multivariate Gaussian distribution. The mean and covariance of that Gaussian are given by evaluating the mean and covariance functions at the corresponding set of input points. Let (x_1_,…, x_*n*_) denote a collection of *n* points in the input domain. If a function *f* has a Gaussian process distribution, 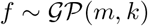, then the vector of function values **f** = (*f*(x_1_),…, *f*(x_*n*_))^⊤^ has a multivariate Gaussian distribution:

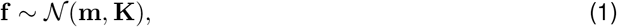

where **m** = (*m*(x_1_),…, *m*(x_*n*_))^⊤^ is the mean vector, and **K** is the (*n* × *n*) covariance matrix whose *i, j*’th element is *k*(x_*i*_, x_*j*_).

### 2.2 GP regression

A common application of GPs is to predict function values at test points given a set of training data consisting of observed inputs and function values. In GP regression, these predictions come from the conditional distribution over unknown function values given the observed values.

Let **X** = (**x**_1_,…, **x**_*n*_) denote a set of *n* input points and let **f** = (*f*(x_1_),…, *f*(x_n_))^⊤^ denote the vector of function values observed at these points. Here we assume the observed data is noiseless, and we will consider noisy observations in Sec. 3. We consider a set of n* novel input points **X**_∗_ = (x_*n*+1_,…, x_*n*+*n*_∗), for which we would like to predict the corresponding (unobserved) function values, denoted **f**_∗_ = (*f*(x_*n*+1_),…, *f*(x_*n*+*n*_∗))^⊤^.

The GP gives us the following joint prior distribution over the observed and unobserved function values:

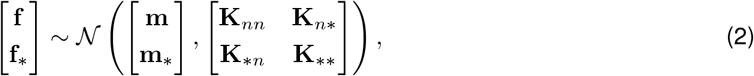

where **m**_∗_ = (*m*(**x**_*n*+1_),…, *m*(**x**_*n*+*n*_∗))^⊤^ is the mean for novel test points in **X**_∗_, and matrices **K**_*n*∗_, **K**_∗*n*_, and **K**_∗∗_ are of size (*n* × *n*^∗^), (*n*^∗^ × *n*), and (*n*^∗^ × *n*^∗^), respectively, formed by evaluating the covariance function k at the relevant points in **X** and **X**_∗_.

By applying the standard formula for Gaussian conditional distributions, we obtain the following conditional distribution of **f**^∗^ given **f**:

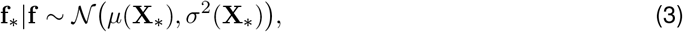

where mean and covariance are given by

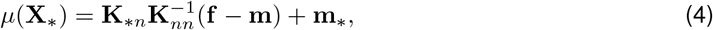

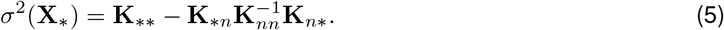

GP regression uses eq. 4, the posterior mean of the function given the training data, to predict function values at test points **X**_∗_ given the training data {**X, f**}.

Although we have assumed so far that the function *f* is scalar-valued, we can extend the GP regression framework to vector-valued functions by using a separate GP for each output dimension of *f*.

## 3 The brain kernel model

Here we introduce the brain kernel (BK) model, which is a probabilistic model of fMRI measurements at an arbitrary set of 3D spatial voxel locations. The brain kernel itself is a covariance function for neural activity that arises under this BK model, which we will infer from registered fMRI data.

### 3.1 Nonlinear embedding function

The first component of the brain kernel model is a nonlinear function, *f*: ℝ^3^ → ℝ^*d*^, which provides a continuous nonlinear mapping from 3D brain coordinates to a d-dimensional latent embedding space. The goal of this mapping is to embed brain regions with similar activity at nearby locations in the embedded space. Typically, we consider *d* > 3, so that the embedding is higher-dimensional than the three physical dimensions of the brain. This gives the embedding flexibility to capture complex non-smooth dependencies between brain regions. One example of such a dependency is the functional symmetry of the two hemispheres [10–12], which suggests that one might wish to map symmetric points on the two hemispheres to nearby points in the embedded latent space. This would not be possible with a continuous mapping in three dimensions. But, in four dimensions, one can fold the three-dimensional brain along the fourth dimension, analogous to the way that folding a 2D brain slice along the mid-line would allow for close alignment of symmetric points from the two hemispheres.

Let **x** ∈ ℝ^3^ denote an input vector, specifying the three-dimensional location of a voxel in the brain, and let **z** ∈ ℝ^*d*^ denote the output of *f*, so that **z** = *f*(**x**) is the *d*-dimensional embedding location of a voxel at **x**. Thus, for a set of voxel locations **X** = (**x**_1_,…, **x**_n_), the embedded locations in the latent space are **Z** = (**z**_1_,…, **z**_*n*_) = (*f*(**x**_1_),…, *f*(**x**_*n*_)).

To impose smoothness on the embedding function, we place a GP prior on *f*. We use a linear mean function, *m*(**x**) = **Bx**, where **B** is a *d* × 3 matrix. This choice ensures that the embedding defaults to a linearly stretched version of the brain in the absence of likelihood terms. Because *f* is a vector function with outputs of dimension *d*, the mean function output is also *d*-dimensional. For the covariance function, we use a Gaussian or “radial basis function” (RBF) covariance for each output dimension of *f*:

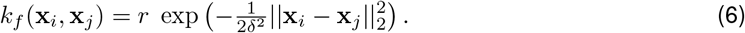

The hyperparameters governing this covariance function consist of a marginal variance *r* and a length-scale *δ*, which control the range and smoothness of *f*, respectively. The GP prior over each output dimension of *f* can therefore be written as

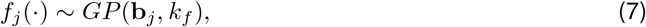

where *f_j_*(·) denotes the *j*th output dimension of the function *f*(·), and **b**_*j*_ is the *j*th row of the matrix **B**. The prior is therefore governed by a set of hyperparameters denoted *θ_f_* = {**B**, *r, δ*}. Thus, all output dimensions of the function f are assumed *a priori* independent with the same covariance function and differing mean functions.

This GP prior over the function *f* implies a multivariate normal prior over any set of embedded voxel locations **Z**. Let **z**_*j*_ denote the *j*th latent embedding of the entire set of brain voxels in the training data. Then the prior over **z**_*j*_ given the true voxel locations **X** is

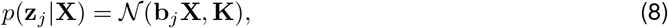

where **K** ∈ ℝ^*n*×*n*^ is the covariance matrix with the *i, j*th element given by *k*(**x**_*i*_, **x**_*j*_) (eq. 6).

### 3.2 From embedding space to neural activity

The second component of the brain kernel model is a probability distribution over neural activity as a function of locations in embedding space. Our modeling assumption is that neural activity changes smoothly as a function of locations in embedding space, or equivalently, that correlations in neural activity decrease smoothly with distance in latent space. We formalize this assumption using the brain kernel, which provides a mapping from latent embedding locations to a covariance matrix for neural activity.

Let **v** ∈ ℝ^*n*^ denote a vector of neural activity from *n* voxels with positions **X** = (**x**_1_,…, **x**_*n*_) and latent embedding locations **Z** = (*f*(**x**_1_),…, (*f*(**x**_*n*_)). The BK model assumes this neural activity vector has a multivariate Gaussian distribution with zero mean and covariance determined by the brain kernel:

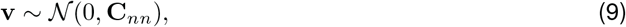

where

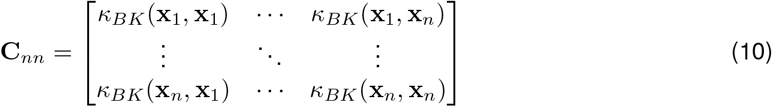

is the covariance matrix, which results from applying the brain kernel *k_BK_*(·, ·) to every pair of voxel locations (**x**_*i*_, **x**_*j*_) in the set **X**.

The brain kernel itself is the bivariate function *k_BK_*: ℝ^3^ × ℝ^3^ → ℝ from pairs of 3D voxel locations to a covariance of neural activity at those pairs of locations:

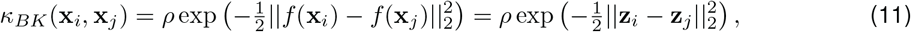

where *ρ* is the marginal variance. The length-scale is omitted here because **z** is an unknown latent variable that we need to optimize which absorbs the unknown length-scale for simplicity. The brain kernel therefore specifies a positive semidefinite covariance matrix for neural activity at any set of 3D voxel locations, which is a function of the embedded latent locations of those voxels via the nonlinear function *f*. The brain kernel model transforms the data representation from voxel space to the latent embedding space, and this transformation explicitly estimates the covariance over neural activity (Figure 1).

**Fig 1.**
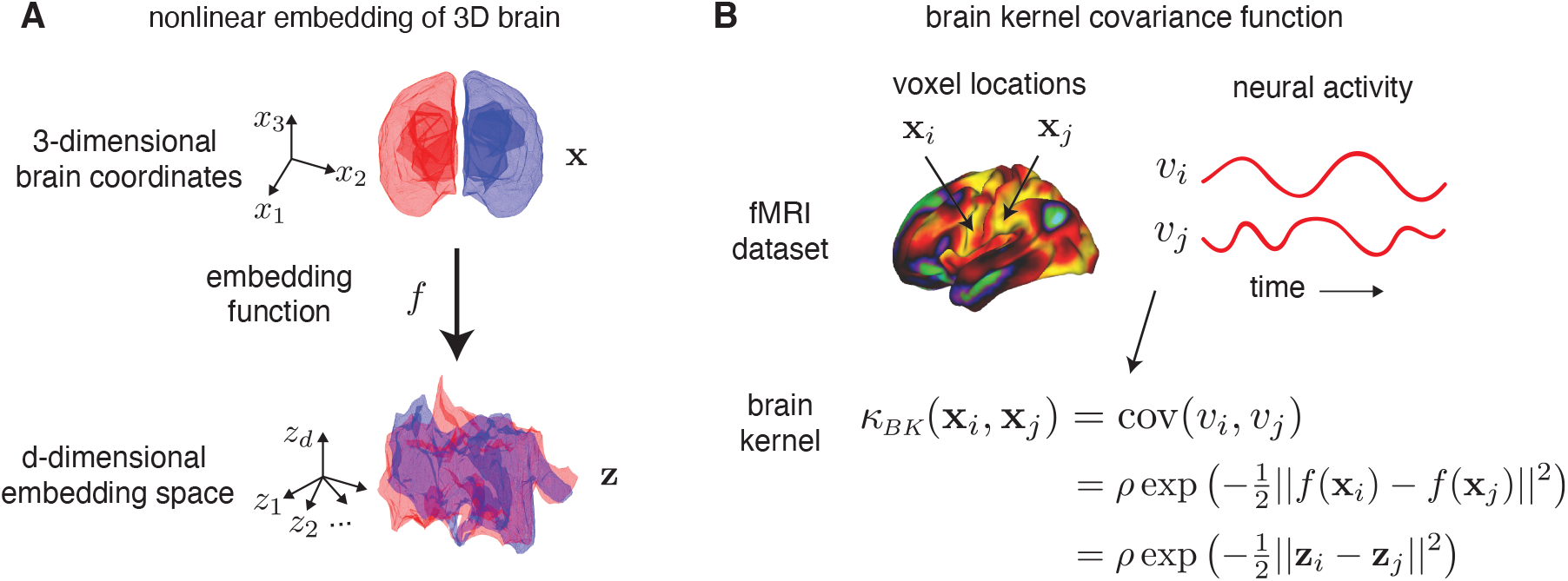
The diagram of the brain kernel model. **A**. The nonlinear latent embedding of the 3D coordinates into a *d*-dimensional latent space using a function *f*. This *f* function is sampled from a GP prior. **B**. Given the voxel locations and the BOLD activity of these two locations in an fMRI dataset (top right), our goal is to construct the brain kernel with the locations as inputs that matches their covariance of the BOLD activity (bottom right). The covariance is equal to the Euclidean-based kernel of the voxels’ embedding in the latent space.

### 3.3 From neural activity to BOLD signal

Next, we assume that the experimenter does not directly measure the neural activity vector **v**, but instead receives measurements corrupted by independent Gaussian noise. If **y** denotes the vector of fMRI measurements, we assume *y_i_* = *v_i_* + *ξ_i_*, where *i* is the index for voxels and 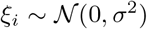 represents measurement noise. The vector y denotes a single measurement of neural activity for all voxels. This induces the following marginal distribution over fMRI measurements given the embedding:

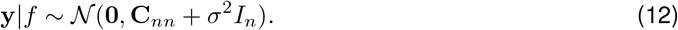

Note however that the brain kernel generates **C**_*nn*_, the covariance of the underlying neural activity **v**, as opposed to the covariance of the noisy fMRI measurements **y**; the covariance of these measurements is (**C**_*nn*_ + σ^2^**I**_*n*_). For simplicity, we will omit the subscript in **C**_*nn*_ in the following text and write simply **C**.

The full set of hyperparameters governing the brain kernel model are therefore *θ* = {**B**, *r, δ*, *ρ*, σ^2^}, where {**B**, *r, δ*} describe the nonlinear embedding function, *ρ* is the marginal variance of neural activity, and σ^2^ is the variance of additive Gaussian noise.

## 4 Inference methods

To fit the brain kernel model, we estimate the latent embeddings of all voxels **Z** as well as the model hyperparameters *θ* = {**B**, *r, δ*, *ρ*, σ^2^} given a time series of whole-brain fMRI measurements Y as well as voxels’ 3D locations **X**. More specifically, **Y** = (**y**_1_,… **y**_*T*_) ∈ ℝ^*n*×*T*^, where *y_t_* ∈ ℝ^*n*^ refers to the vector of fMRI measurements at time index *t* ∈ {1,…, *T*}. Let **X** = (**x**_1_,…, **x**_*n*_) ∈ ℝ^3×*n*^, where **x**_*i*_ ∈ ℝ^3^ and *i* ∈ {1,…, *n*}, denote the set of *n* 3D voxel locations for this dataset. Let **Z** = (**z**_1_,…, **z**_*n*_) = (*f*(**x**_1_),…, *f*(**x**_n_)) ∈ ℝ^*d*×*n*^, where **z**_*i*_ ∈ ℝ^*d*^ denotes the set of *n* voxels’ *d*-dimensional latent representations.

We propose two empirical estimators: maximum a posteriori (MAP) and penalized least squares (PLS). Briefly, MAP solves the problem using conventional Bayesian probabilistic inference. Since we have already defined the distribution to generate fMRI measurements from the latent embeddings (eq. 12) and the prior for the latent embeddings (eq. 8), we can estimate the latent embeddings **Z** using MAP methods. PLS formulates an objective function by minimizing the squared error between the sample covariance of the data cov(**Y**), and the model-defined covariance **C** (eq. 10). The goal of both inference methods is to find the latent embedding locations **Z** and the model hyperparameters such that the neural activity covariance **C** resembles the sample covariance of **Y** as closely as possible.

### 4.1 Maximum a posteriori (MAP) estimation

Estimating large covariance matrices is a fundamental problem in modern multivariate analysis. One common approach uses maximum likelihood methods. We have already defined the data distribution **Y** given the latent embedding **Z** in eq. 12 and described the prior over **Z** in eq. 8.

The joint distribution is then

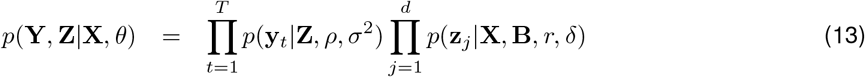

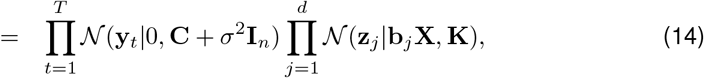

where **C** is a function of **Z** and *ρ*. Thus, the loss function for the maximum a posteriori (MAP) estimator is

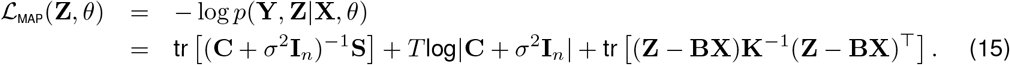

**S** is the sample covariance of measured neural activity, defined as

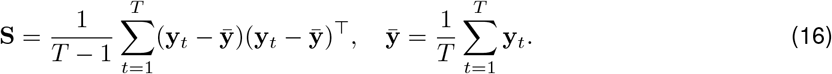

Minimizing 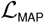 w.r.t. **Z** and *θ*, we derive the MAP estimators 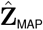 and *θ*_MAP_.

### 4.2 Penalized least squares (PLS) estimation

Another common approach aims at finding an estimator that resembles the sample covariance while also satisfying structural assumptions about the data [13, 14]. In prior work, Fan et al. [14] showed that a generalized thresholding covariance estimator can be cast as a penalized least squares (PLS) problem:

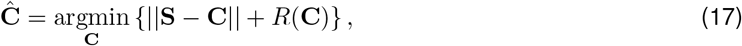

where *R*(·) is a penalty function that imposes structure on the covariance matrix **C**. Sparsity in **C** is often encoded using a shrinkage penalty. However, we abandon sparsity in the brain kernel because the covariance of brain activity may have dense structures. Instead, we regularize the latent subspace of the covariance matrix using the normal prior on the latent embedding **Z** (eq. 8), which is a Bayesian regularization of the log likelihood with the form tr [(**Z** − **BX**)K^−1^(**Z** − **BX**)^⊤^]. This is equivalent to an *l*_2_-norm penalty on 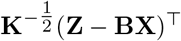. Including the noise variance term σ^2^, the loss function for the empirical estimator is

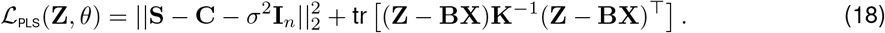

Minimizing 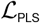 w.r.t. **Z** and *θ*, we derive the empirical PLS estimators 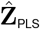 and *θ*_PLS_. Here, the least square term estimates the embedding **Z** using the sample covariance, while the second term regularizes **Z** with the kernel matrix **K** and mean values **BX** constructed from **X**. Eq. 18 can also be considered as inheriting the log of the GP prior from eq. 15 and replacing the data likelihood term with a squared loss.

### 4.3 Exact inference by block coordinate descent

Calculating the log posterior (eq. 15) for MAP inference has a computational complexity of *O*(*n*^3^), due to the need to compute the inverse and the determinant of the covariance matrix. This cost is often impractical in fMRI settings, where the number of voxels n may be on the order of thousands to hundreds of thousands.

To optimize **Z** and the model hyperparameters, we could use gradient descent or Newton’s method as the optimizer; however, this approach is computationally impractical. Thus, we need to consider a scalable inference method. Existing scalable inference methods for large datasets [15–17] exploit low-rank approximations to the full Gaussian process. However, these approximations suffer from a loss of accuracy in covariance estimation. Thus, we develop a block coordinate descent (BCD) algorithm as an exact inference method for the brain kernel model (see Methods, Algorithm 1). Coordinate descent has been successfully applied to solve penalized regression models [18], to estimate covariance graphical lasso models [19], and to compute large-scale sparse inverse covariance matrices [4]. Our PLS and MAP estimators are non-convex smooth functions. We apply an iterative block coordinate descent method solved by the proximal Newton approach [20]. Given such a scalable optimizer, we alternate between optimizing **Z** and the hyperparameters using either eq. 15 (MAP) or eq. 18 (PLS) as the objective loss function. More details about the optimization can be found in the Methods section.

In practice, we used the PLS estimate to initialize the MAP estimate, as PLS optimization is faster and requires less memory, but the MAP estimate is more principled and achieves a higher accuracy. We used the BCD algorithm to optimize both the PLS and MAP objectives.

### 4.4 Predicting activity for new voxels

Although we fit the brain kernel model to data collected with a particular grid of voxels, our framework allows us to apply the model to fMRI measurements collected using different voxel grids. To do so, we use the fact that the brain kernel is defined using a Gaussian process; the mean of this GP provides a smooth mapping from 3D voxel space to the *d*-dimensional latent embedding space, which can be evaluated at any 3D brain locations. We obtain the embedding location of a novel 3D voxel location **x**^∗^ using the posterior mean of this GP:

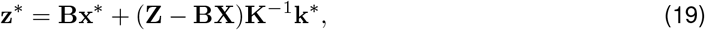

where **k**^∗^ = [*k_f_* (**x**^∗^, **x**_1_), …, *k_f_* (**x**^∗^, **x**_*n*_)]^⊤^ represents the vector formed by evaluating the RBF covariance function for the voxel at location **x**^∗^ and all the observed voxels in **X**. The brain kernel for any arbitrary pair of voxel locations in the test set 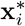 and 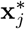 is therefore given by:

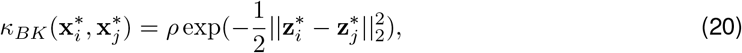

with 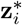 and 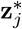 the corresponding latent embeddings (eq. 19).

Now given the newly estimated brain kernel and the new voxel location **x**^∗^, we could predict its activity via

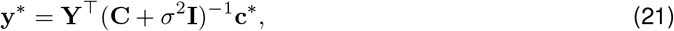

where **c**^∗^ = [*k_BK_*(**x**^∗^, **x**_1_),…, *k_BK_*(**x**^∗^, **x**_*n*_)]^⊤^ ∈ ℝ^*n*×1^ and **y**^∗^ ∈ ℝ^*T*×1^ is the predicted activity of the new voxel across all measurements.

### 4.5 Synthetic experiments

To illustrate the performance of our proposed inference method to optimize the brain kernel objective function, we began with an application to simulated data, where the ground-truth embedding is known. We first compared our block coordinate descent (BCD) method (Algorithm 1), which maximizes the exact log evidence (eq. 15), with two variational inference methods that optimize a lower bound on log evidence: an inducing-point method [15], and a dynamical variational inference method (dynamic VIP^1^) [16].

We created a simulated dataset using a 1-dimensional brain and 1-dimensional latent embedding function, sampled from the brain kernel model (Fig. 2). Here, the latent embedding is a nonlinearly warped version of the 1D brain. We considered a set of 100 voxel locations on an evenly spaced 1D grid: **X** = [1, 2,…, 100]^⊤^. We then sampled the voxels’ latent locations **Z** from a GP with mean *m*(x) = 0.6x and RBF covariance with length-scale *δ* = 10 and marginal variance *r* = 9: 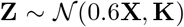, where **K**_*ij*_ = 9 exp(−(*i* − *j*)^2^/200). Given this embedding function, the brain kernel defines a covariance matrix for neural activity at these 100 voxels, denoted **C**, with the *i, j*th entry given by **C**_*ij*_ = exp(−(**z**_*i*_ − **z**_*j*_)^2^/2) (Fig. 2A top). To obtain simulated fMRI measurements, we sampled 750 observations from a Gaussian distribution with zero mean and covariance **C** + 5*I*, where σ^2^ = 5 represents the variance of additive measurement noise.

**Fig 2.**
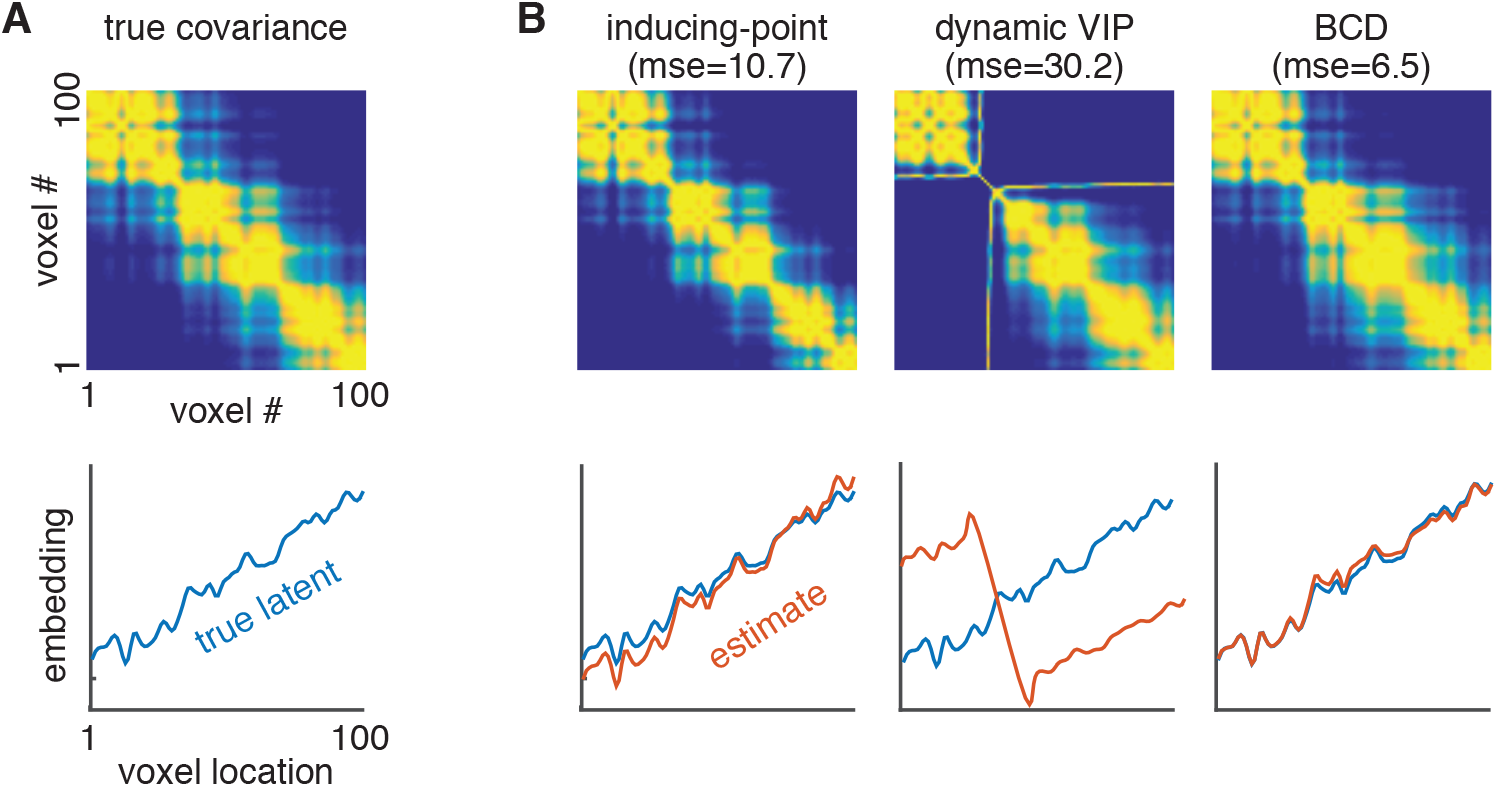
Recovery of 1D brain from synthetic data. **A**. The true covariance matrix for neural activity at 100 evenly-spaced voxels in a 1D brain (top), generated by a 1D latent embedding function sampled from the 1D brain kernel model (bottom, blue curve). **B**. Model estimates. The first column is the inducing-point method; the second column is dynamic VIP; and the last column is our BCD method. In each column, we show the estimated covariance matrix (top) and the estimated 1D latent embeddings (bottom, red curve). We also show the mean squared error (MSE) between the estimated covariance matrix and the true covariance matrix.

We compared the different estimators on the task of recovering the true covariance and latent embedding function from this dataset (Fig. 2B). The inducing-point method with six inducing points (column 1) performed well at recovering the latent embedding function, although it was outperformed by our BCD estimator in terms of mean squared error (BCD; column 3). The dynamic VIP estimate with six inducing points (column 2) converged to a local optimum far from the true latent embedding, yielding a substantially higher error. In contrast, exact inference using BCD outperformed both inducing point-based approximate methods in terms of accuracy at recovering the true latent from simulated data.

Next, we conducted a set of synthetic experiments to examine how well different models captured the covariance of simulated fMRI data, using voxels on a 1-dimensional, 2-dimensional, or 3-dimensional grid. We fit these simulated datasets using the brain kernel model optimized with (1) BCD, (2) variational inducing-point, and (3) the dynamic VIP method, and two additional models: (4) a linear brain kernel (LBK) model, and (5) a GP with RBF covariance function (RBF). Note that the first three methods are based on the original brain kernel model but have different inference methods, while the last two methods represent simplifications of the proposed brain kernel model. The linear brain kernel model assumes a purely linear embedding function; it thus allows rotating and linearly dilating the voxel grid, but does not allow for nonlinear warping of voxel locations. The covariance function for neural activity v under the LBK model is given by:

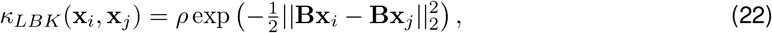

where **B** is a *d* × *v* linear embedding matrix and *v* is the number of dimensions of *x*, e.g., *v* = 2 for a 2-dimensional grid of voxels, and *d* is the dimension of the latent space. To assess the importance of the structured latent embedding in covariance estimation, we fit the data with a standard GP with RBF covariance function:

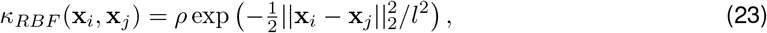

where *l* is the length-scale for the RBF kernel. This model imposes smoothness using Euclidean distance, without estimating any (linear or nonlinear) transformation of the true voxel locations. To fit the LBK and RBF models, we minimized the following negative log likelihood for hyperparameters *θ*:

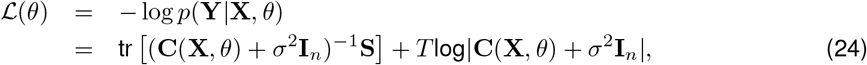

where **C**(**X**, *θ*) is the model-generated covariance matrix (given by eq. 22 for the linear brain kernel model with *θ* = {**B**, *ρ*}, and by eq. 23 for the RBF model with *θ* = {*l*, *ρ*}), **S** is the sample covariance (eq. 16), σ^2^ is the noise variance, and *T* is the number of samples in the simulated dataset.

For each grid dimension, we simulated ten independent datasets, each with different nonlinear embedding functions sampled from the brain kernel model. For the 1D experiments, we generated each dataset with 500 voxels and 750 samples, and embedded the 1D voxel space to a 1D latent embedding space. For the 2D experiments, we used a 25 × 25 voxel grid, embedded nonlinearly in a 3D latent embedding space, and generated datasets of 1000 samples, where each sample is a vector of 625 noisy measurements of brain activity at voxel locations. For the 3D datasets, we used a 10 × 10 × 10 grid of voxels, embedded nonlinearly into a 6D latent space, and we generated 1500 samples per dataset.

To compare models, we performed two different cross-validation tests: (i) prediction on held-out voxels (Fig. 3A), and (ii) predictions on held-out samples (Fig. 3B).

**Fig 3.**
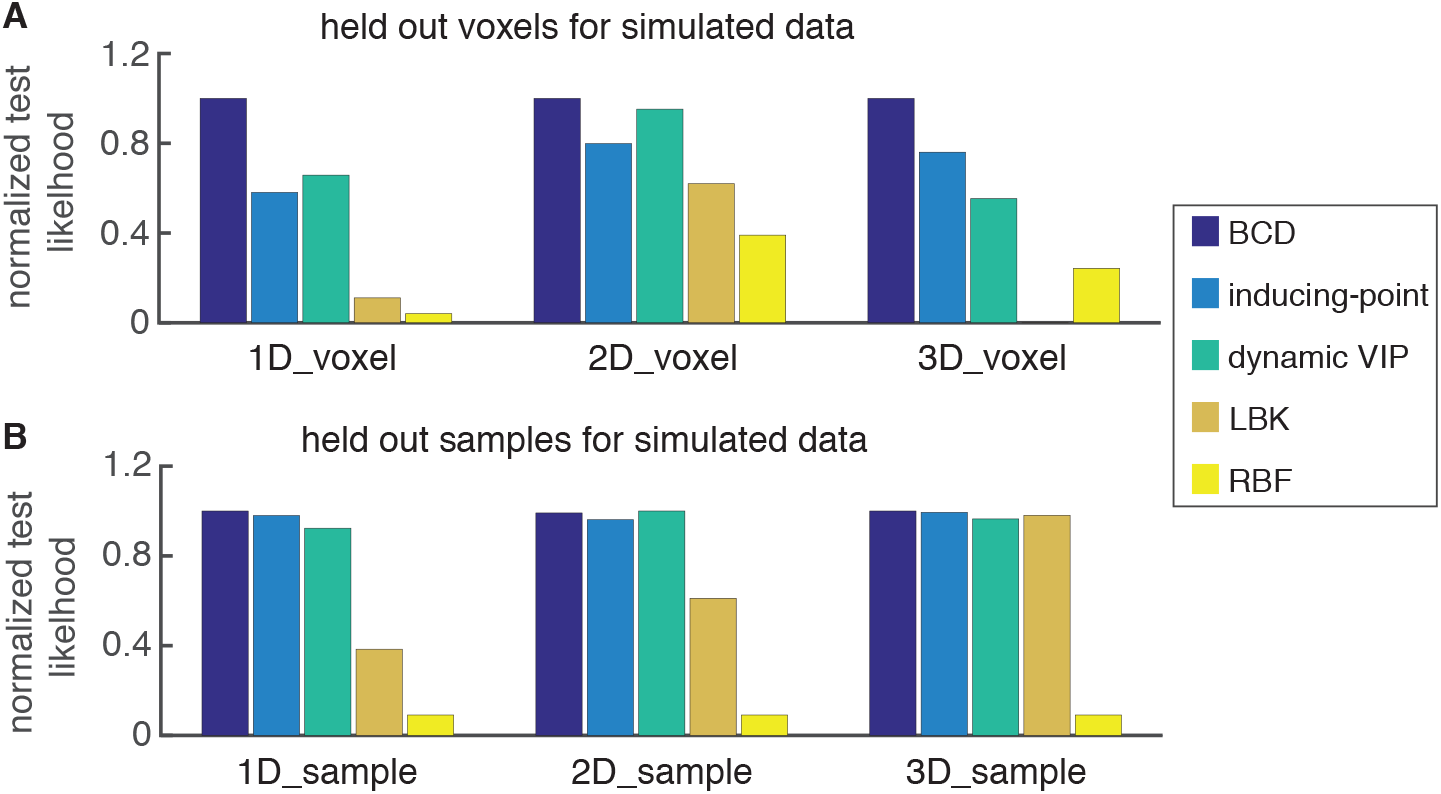
Quantitative comparisons for BCD, inducing-point, dynamic VIP, LBK, and RBF on three simulated datasets with 1D, 2D, and 3D input voxel locations. **A** is the held-out voxel experiment, and **B** is the held-out sample experiment. Within each experiment, we show the normalized test likelihood values for each method with three different simulated datasets.

For the first of the two cross-validation tests, we removed ten randomly-selected voxels **X**^∗^ from the simulated 1D datasets and set them aside as test data, and we optimized the model parameters on a training set of measurements from the remaining 490 voxels. (For experiments with 2D and 3D grids, we set aside 62 and 100 voxels as held-out data, and we trained models using the remaining 563 and 900 voxels, respectively.) For the BK and LBK models, we computed the embedding locations **Z**^∗^ for the test voxels using the GP posterior mean given the inferred embedding locations **Z** (eq. 19), and we used these locations to evaluate the covariance of the test voxel activity (eq. 21). For the RBF model, there is no embedding, so the covariance of the test data depends only on the test voxel locations **X**^∗^. We used the resulting predictive covariance to compute the log likelihood of the test data. We found that the BCD-optimized brain kernel model estimate outperformed other methods, while the LBK and RBF models performed worse, presumably due to their inability to capture the nonlinear embedding of the simulated data (Fig. 3A).

Next, we evaluated cross-validation performance on held-out samples, which were measured at the same set of voxels as the training set. For these simulations, we randomly selected 75 samples as test data for the 1D datasets, and used the remaining 675 samples to fit the five models. (For 2D and 3D grids, we used a train-test split of 900:100 and 1350:150 samples, respectively.) We estimated a covariance function using measurements in the training set, and computed a test likelihood using this covariance function on the test set. We generated ten random splits for each dataset and computed the normalized test log likelihood. In this predictive task, the BCD estimate again outperformed other methods (Fig. 3B).

Overall, these simulations demonstrated that the inducing point-based GP methods, which are often used due to their scalability, had higher errors than our exact BCD inference method. In addition, by comparing to the linear brain kernel model and the standard GP model with an RBF kernel, we showed that the ability to capture a nonlinear transformation of the voxel locations is critical for accurately modeling the covariance of simulated brain activity.

## 5 Inferring the brain kernel from resting-state fMRI data

Now that we have described the brain kernel model and validated our inference method using simulated data, we turn to the problem of inferring the brain kernel from real data. We fit the brain kernel model to large-scale publicly-available resting-state fMRI data from the Human Connectome Project (HCP) [6]. An advantage of this approach is the large size of the HCP sample, which mitigates overfitting and allows us to learn an embedding function that captures correlation patterns common to a vast collection of different brains. However, applying the resulting brain kernel to task-based fMRI datasets assumes that the correlations presented in resting-state fMRI data are applicable to activation patterns in other brain states. Previous studies have shown interesting relations between brain activity during a task and at rest. One study showed that coherent spontaneous activity accounted for variability in event-related BOLD responses [21]. A second study concluded that functional networks used by the brain in action were continuously and dynamically “active” even when at “rest” [22]. A third study showed that resting-state activation patterns had strong statistical similarities to cognitive task activation patterns [23]. These provide reasons for optimism, though the possibility of changes in correlation across different tasks could potentially affect the application of the brain kernel which we will show empirically later.

We examined resting-state fMRI data from 812 subjects collected via the Human Connectome Project (HCP) [6]. These data were acquired in four fMRI runs of approximately 15 minutes each, two runs in one session and two in another session, with eyes open and relaxed fixation on a projected bright cross-hair on a dark background. A sophisticated preprocessing pipeline was used to align voxels across subjects. Detailed information about imaging protocols, image acquisition, and preprocessing can be found in [24]. The resulting dataset consisted of 59,412 voxels in a cortical surface coordinate system. Although the full covariance matrix of these data is of size ≈ 59*K* × 59*K*, we fit the model using the first 4500 eigenvectors of the full matrix provided by HCP.

We fit the brain kernel with different numbers of latent dimensions and computed the test log likelihood with held-out voxels and samples (Fig. 4D). We found that test performance plateaued with increasing dimensionality, and selected *d* = 20 dimensions for subsequent analyses. We then estimated the brain kernel by fitting the 20-dimensional latent embedding for each voxel using BCD optimization of eq. 18 and eq. 15. The resulting function is a matrix of embedding locations of size ≈ 59*K* × 20, where each row contains the embedded location of a single voxel in the resting-state fMRI dataset. This embedding, in addition to the hyperparameters (the linear projection matrix B and the hyperparameters {*γ, δ*} for **K**), provides a full parametrization of the brain kernel. The entire fitting process took approximately 1 week on a single CPU, since we needed to optimize the latent embedding location of each voxel in the HCP dataset so that the covariance in BOLD activity for any pair of voxels was accurately described by the brain kernel.

**Fig 4.**
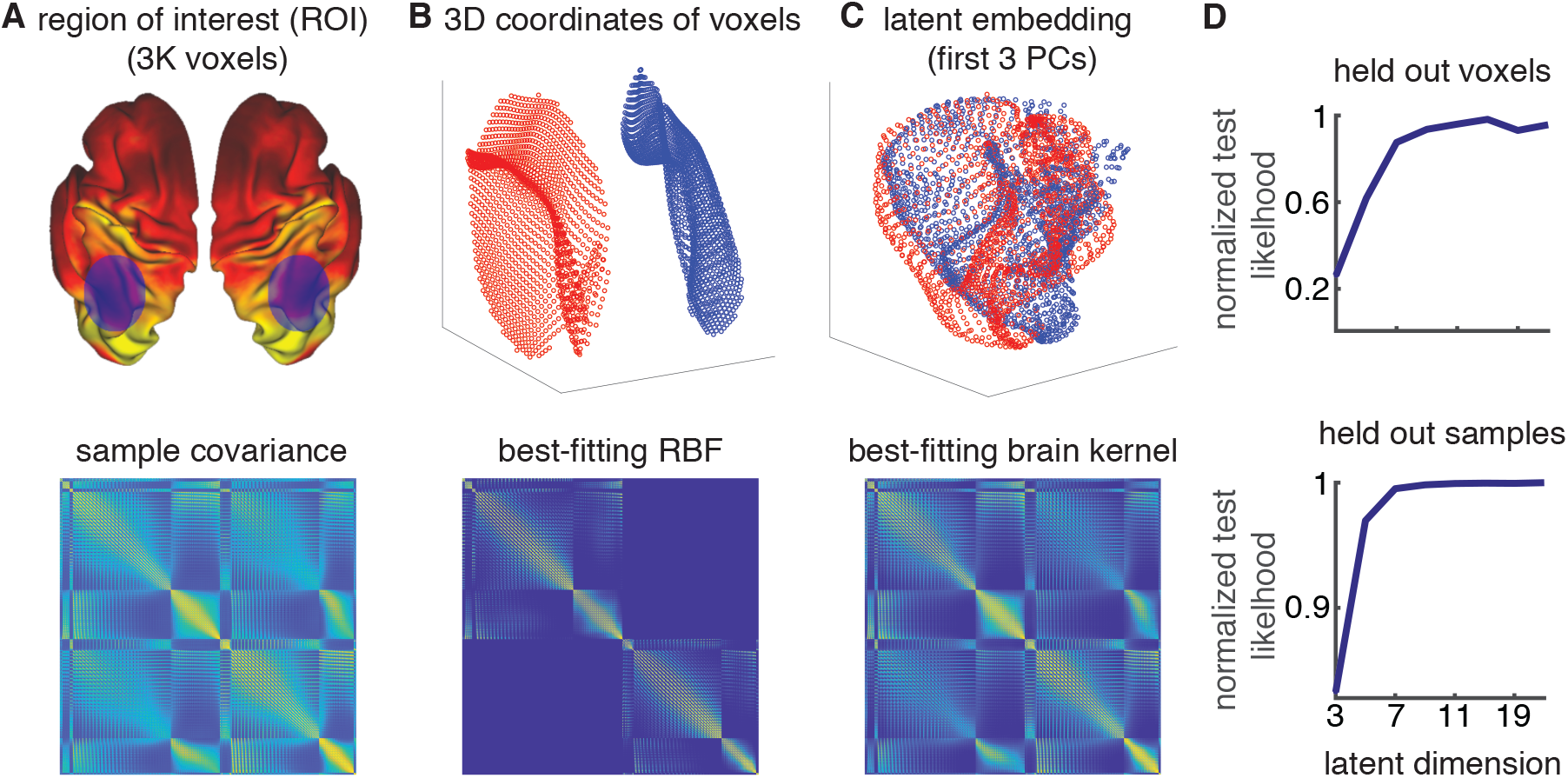
3D embeddings and the corresponding covariance matrices of the resting-state fMRI data. **A**. The two example regions of interest (ROIs) we selected in the left and right parietal lobes (top) and the full covariance for the 3K voxels in these two ROIs (bottom). **B**. The original 3D voxel coordinates (top) and the best-fitting RBF kernel (bottom). **C**. A 3-dimensional projection of the estimated 20-dimensional brain kernel embedding (top) and the corresponding brain kernel covariance matrix (bottom). **D** We show the influence of the latent dimensionality on the predictive performance for held-out voxels and held-out samples with the resting-state fMRI data.

Although it is impossible to visualize the 20-dimensional nonlinear embedding that defines the brain kernel, we can gain insight into its shape by plotting low-dimensional projections on subsets of voxels. To visualize the brain kernel, we selected two symmetric ROIs in the left and right parietal lobes (Fig. 4A, top). Taken together, these two ROIs contain 3K voxels. We visualize the sample covariance of these voxels (Figure 4A, bottom). This covariance contains four identifiable blocks: two diagonal blocks that correspond to the covariance of voxels within each ROI, and two off-diagonal blocks that correspond to cross-ROI covariance. The off-diagonal blocks reveal that the two ROIs have reasonably strong correlations despite being spatially distant in the brain.

The original 3D voxel coordinates (Fig. 4B, top) and the covariance given by the best-fitting RBF kernel (eq. 23; Fig. 4B, bottom) show that the RBF kernel model fails to capture the the off-diagonal blocks of the sample covariance, corresponding to covariance between voxels in opposite hemispheres, due to the fact that the RBF kernel depends only on the Euclidean distance between voxels. In contrast, a 3-dimensional projection of the estimated 20-dimensional brain kernel embedding (Fig. 4C, top) and the corresponding brain kernel covariance (Fig. 4C, bottom) appear to capture this cross-ROI structure in the covariance matrix. The 3D projection corresponds to the first three principal components of the 20-dimensional latent embeddings, and shows that paired voxels from opposite hemispheres are embedded close to each other under the brain kernel.

Conversely, some voxels that are physically close together in the brain are embedded far apart in the embedding space. As an example, we presented a visualization of some selected 3D coordinates (Fig. 5A) and their corresponding 3D latents (Fig. 5B) of three regions on the left hemisphere (color coded). We can see the green region is closer to the orange region in the 3D voxel space, while it’s closer to the blue region in the latent space. The blue region covers mostly the visual area. The green region contains motor functions including eye movement and orientation. The red region corresponds to higher mental functions such as planning and emotion. This could explain the stronger functional connectivity between green and blue in the latent space.

**Fig 5.**
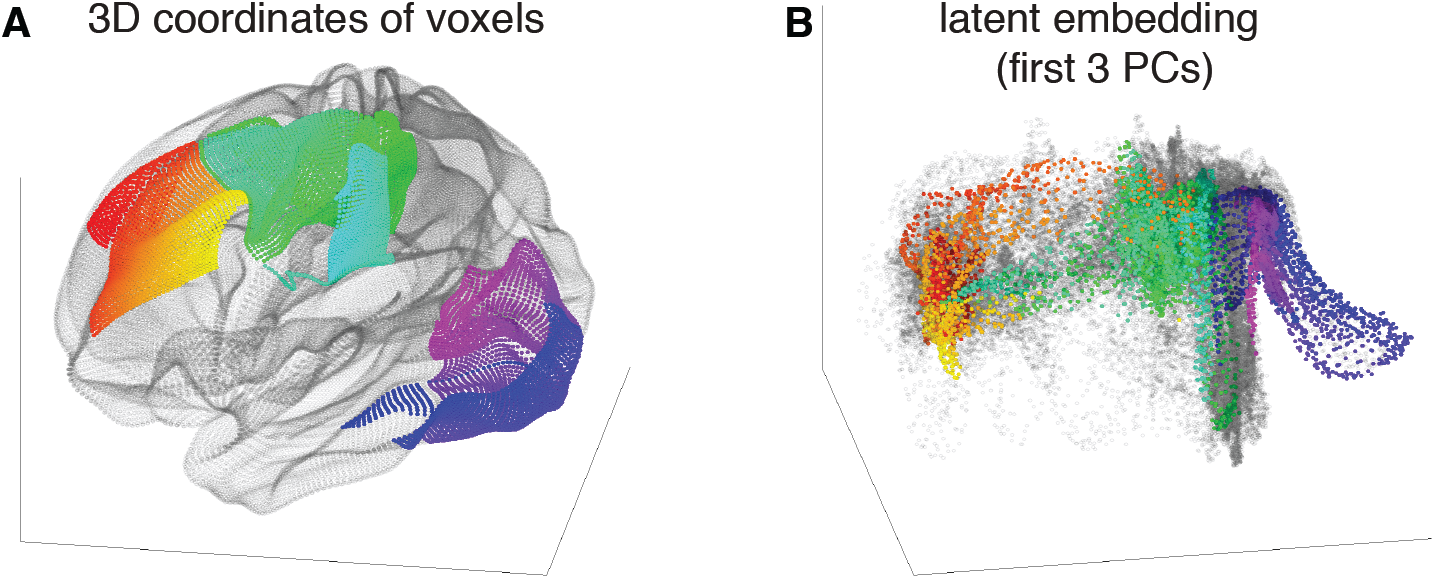
3D coordinates (A) and their latent embeddings (B) of three regions on the left hemisphere. Opposite to Fig. 4B and C where the ROIs are separate in the voxel space but overlap in the latent space, the orange region and the green region are closer in the voxel space but distant to each other in the latent space, compared to the relation between the green region and the blue region.

Overall, this nonlinear embedding allows the brain kernel to more accurately capture the covariance of the real data.

## 6 Applications

To illustrate the usefulness of the brain kernel, we applied it to two different fMRI data analysis problems: decoding and factor modeling (Fig. 6). For both analyses, we used the latent embedding function fit to resting state data (as described in Sec. 5), and tuned two hyperparameters governing the amplitude and length-scale of the brain kernel, which allowed it to adapt to the statistics of each task dataset of interest. We refer to this as *tuning* of the brain kernel covariance for use in applications. We describe these applications in detail below.

**Fig 6.**
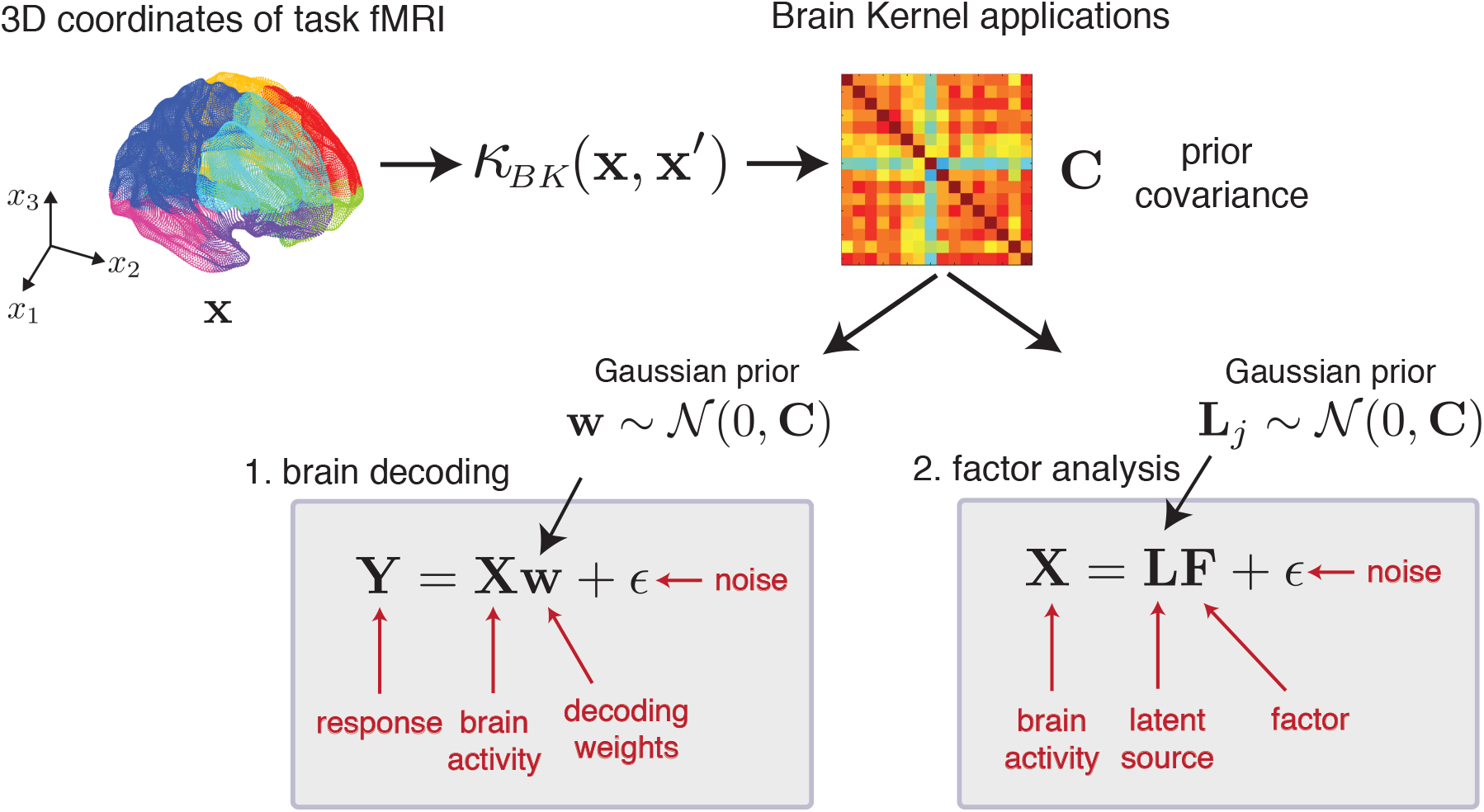
Schematic figure illustrating two applications of the brain kernel to task fMRI data. After fitting the brain kernel to resting-state data, we applied it to task fMRI data with *n* voxels by evaluating the brain kernel at the 3D voxel locations. This results in a *n* × *n* prior covariance matrix for the task data, denoted **C**. We then used the covariance **C** as the prior covariance for two modeling tasks: (1) brain decoding and (2) factor analysis. Two unknown parameters **w** and **L** are both random variables with a Gaussian prior whose covariance is **C**. Thus, we effectively imposed assumptions on the structure of w and **L** via **C**.

### 6.1 Brain decoding

In this section, we illustrate how the brain kernel can be applied to fMRI classification (or “decoding”) tasks.

#### 6.1.1 HCP tasks

We first examined the task fMRI datasets in the HCP database. We explored the working memory task, the gambling task [25], the language processing task [26], the motor task [27], the emotion processing task [28], the relational processing task [29], and the social cognition task (more details found in [24, 30]). We will elaborate on the working memory and gambling tasks and finally summarize the result with all datasets.

##### Working memory task

We obtained the working memory task fMRI measurements from the HCP [30]. The stimuli consisted of four types of pictures: places, tools, faces, and body parts. Stimuli were presented for 2 s on each trial followed by a 500-ms inter-trial interval (ITI). Each task block consisted of ten trials of 2.5 s each. Each run contained 8 task blocks, half for a 2-back working memory task and half for a 0-back working memory task, along with 4 fixation blocks. Two runs were collected for each subject, with 405 volumes per run or approximately 5 minutes. Because the task fMRI was from the same HCP project as the resting-state fMRI, they shared the same coordinate system and preprocessing pipeline. The task fMRI was aligned with the MNI template with the same 59,412 voxels as the resting-state fMRI data. Instead of analyzing the whole-brain data for brain decoding, we worked with functional ROIs. For each presented type of pictures, we took the group-average task contrast for the 2-back vs 0-back working memory task and kept all voxel coordinates whose z-statistics were at least −1.96 or +1.96 standard deviations away from the mean, indicating that the voxels were statistically significant in the contrast map. We repeated the same thresholding procedure for all objects and took a union set of all coordinates to formulate the functional ROIs for the working memory task. We didn’t select the ROIs using the contrast among objects, therefore the resulting ROIs contained no discriminative information with respect to the decoding task.

The experiment required observers to perform a working memory task using four different types of objects. We tested the ability to decode these objects from fMRI data by fitting a binary linear classifier for each pair of objects. We used Bayesian linear regression classifiers to solve six binary classification problems. A Bayesian linear regression classifier has the form,

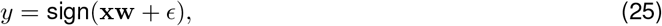

where x is a vector of all voxels for one fMRI measurement or sample, *y* is a ±1 label indicating the binary object category for that sample, **w** is a vector of regression coefficients, and *ϵ* is independent zero-mean Gaussian noise with variance σ^2^. To regularize the estimate of **w**, we assumed a zero-mean Gaussian prior with covariance **C**, i.e., 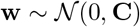. We considered three different choices of prior covariance: (1) a ridge prior, which corresponds to a diagonal covariance with a positive constant along the diagonal; (2) a radial basis function (RBF) covariance (eq. 23), which imposes smoothness based on the voxels’ 3D locations in the brain, and (3) the brain kernel defined as

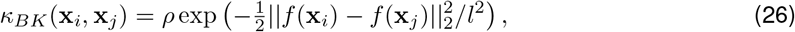

where *f* is the nonlinear embedding function fitted in Sec. 5 and is fixed for decoding. {*ρ, l*} are tuneable hyperparameters that we optimized when tuning the brain kernel to the task data of interest. The computational cost of this tuning is the same as the cost for optimizing the standard kernels such as the RBF covariance, which has an equivalent pair of hyperparameters. This optimization is fast (e.g., 80 s for 3000 voxels on a CPU), and there is no difference in computational cost between the brain kernel and the RBF smoothing prior.

We trained the classifier with these three priors on one run and calculated accuracy performance on the second run, then repeated the same procedure with test and training sets reversed. When we trained the model, we randomly split the training run into five folds and selected the optimal hyperparameters in the covariance functions via 5-fold cross validation. After cross validation, we applied the optimal hyperparameters to the test run for each prior model. We repeated this 5-fold cross validation experiment ten times to reduce variability. We used linear regression to train the classifier, so the ±1 labels were treated as continuous target values, and test accuracy was evaluated by taking the sign of the prediction. We plotted averaged accuracy across both runs and all repeats for the three priors for ten randomly-selected subjects (Fig. 7A). The RBF prior outperformed the ridge prior, but the brain kernel prior outperformed both of the other priors, indicating that smoothing in a nonlinear embedding space defined by correlations of fMRI signals provided additional benefits in regularizing weights for a classification task. Moreover, the framework of the brain kernel for regularization allows us to visualize the inferred decoding weights overlaid on a 3D brain (Figure 7B).

**Fig 7.**
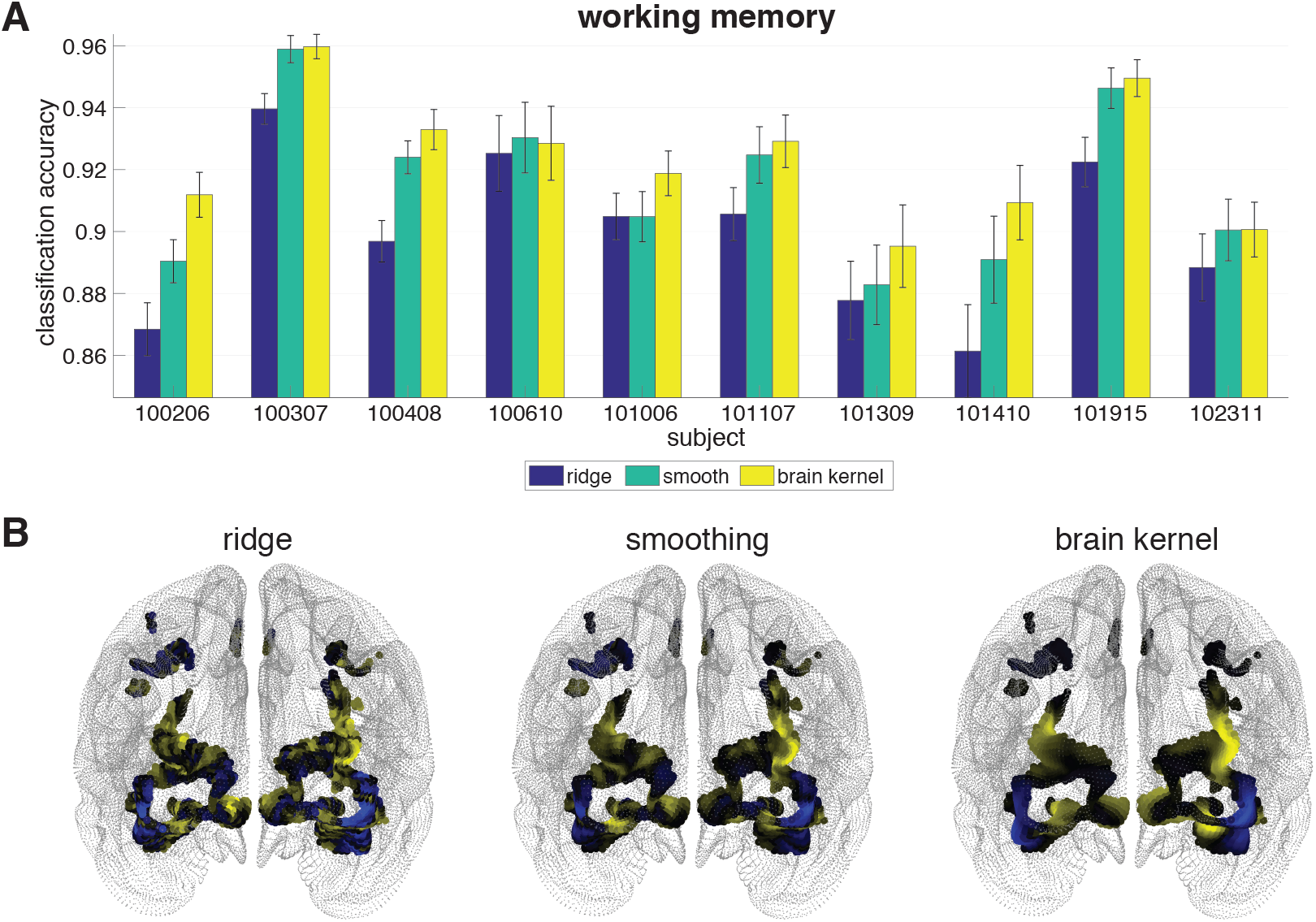
**A**. Accuracy performance on the working memory task. The x-axis indicates subject identifiers. The y-axis is accuracy performance. We compared our brain kernel with a ridge prior and a smooth RBF kernel, color coded. The error bars indicate standard errors. **B**. Visualization of an example set of decoding weights. Blue indicates negative values and yellow indicates positive values.

##### Gambling task

We next examined fMRI data in a gambling task from the HCP [30], adapted from a prior study [25]. Participants were asked to play a card-guessing game. They were shown a mystery card with a number that could range from 1 to 9. They needed to guess whether it was more or less than 5 by pressing on of two buttons. If they made the correct guess, the card showed a green up arrow with “$1” for rewards; if they guessed wrongly, the card showed a red down arrow with “-$0.5” for losses; if the true value was 5, they got a neutral response without win or loss. Participants had 1500 ms to guess, and the feedback was presented for 1000 ms, followed by a 1000-ms inter-trial interval (ITI). Each task block consisted of eight trials that were either mostly reward or mostly loss. Two runs were collected for each subject. Each run contained two mostly reward and two mostly loss blocks, interleaved with four fixation blocks. There were 253 volumes per run, which lasted approximately 3 minutes. The task fMRI was aligned to the MNI template and had the same 59,412 voxels as the resting-state fMRI data. Instead of analyzing the whole-brain data for brain decoding, we worked with functional ROIs which were selected using the same approach as described in the working memory task.

We formulated the task as a binary classification problem separating reward trials and punishment trials. We used the same Bayesian linear regression classifiers as described in the working memory section. We trained the classifier with three priors on one run and calculated the accuracy performance on the second run, then switched the training and test runs. This procedure was repeated ten times. The ±1 labels were treated as continuous target values during training, and the test accuracy was evaluated by taking the sign of the prediction. We computed the averaged accuracies across two runs and 10 repetitions for the three priors for 15 subjects (Fig. 8A). For 11 out of 15 subjects, the brain kernel improved accuracy over both the ridge prior and the smooth RBF prior. For two of the remaining four subjects, the brain kernel outperformed the ridge prior. The discrimination problem with the gambling data was more difficult than the working memory task. Many activations occurred in the visual cortex and the prefrontal cortex for working memory, whereas critical activations for the reward task might be localized to the striatum, which was not included in the cortical data used here. However, with the cortical brain kernel, we were still able to improve the predictive ability of the Bayesian decoding model.

**Fig 8.**
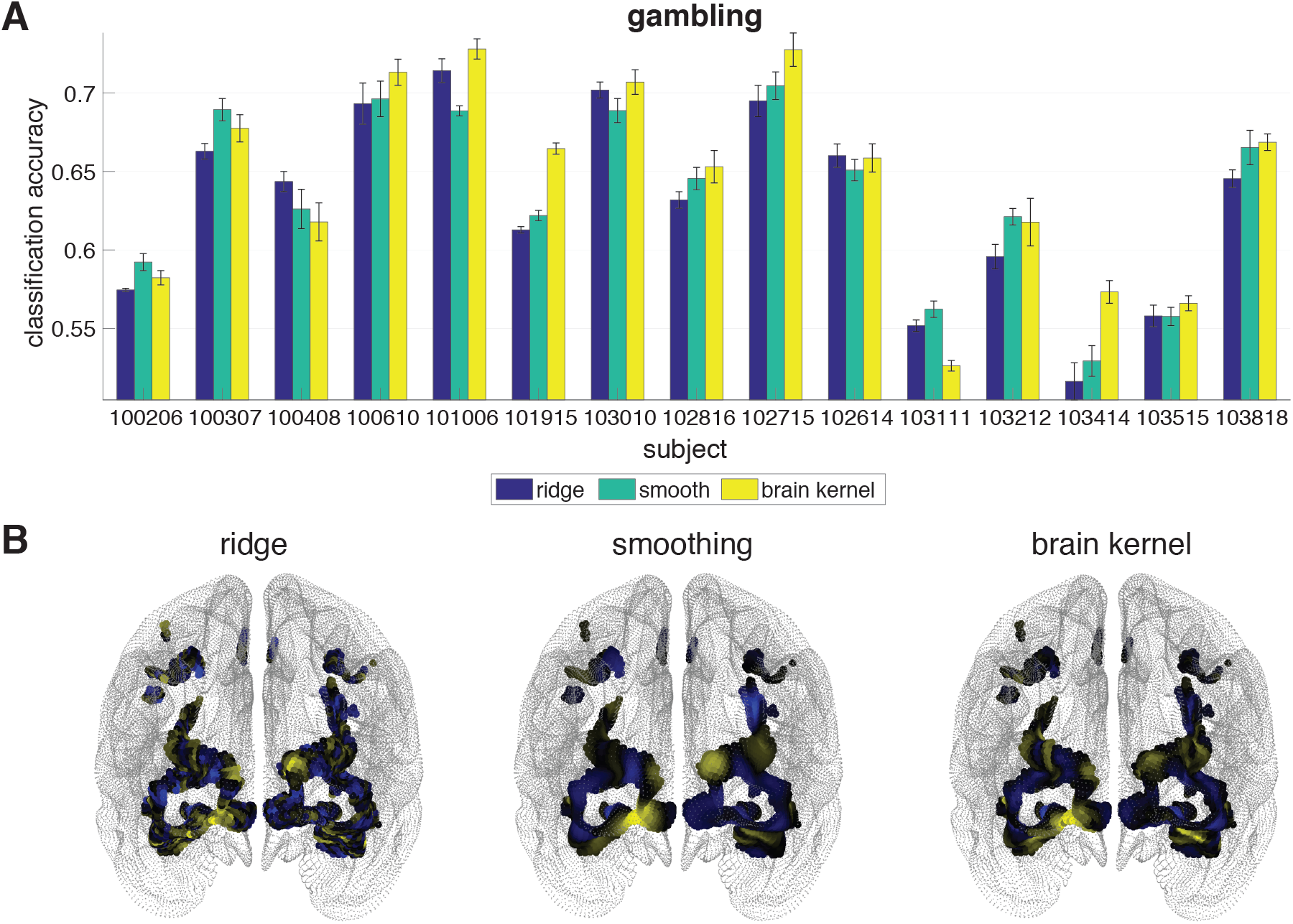
**A**. Accuracy performance on the gambling task. The x-axis indicates subject identifiers. The y-axis is accuracy performance. We compared our brain kernel with a ridge prior and a smooth RBF kernel, color coded. **B**. Visualization of an example set of decoding weights. Blue indicates negative values and yellow indicates positive values.

##### All HCP tasks

We’ve elaborated on the working memory and gambling tasks above. We also achieved the classification accuracy performance for all other tasks (presented in Appendix B.1). Here we summarize the averaged accuracy over all subjects for each task in Fig. 9. Consistent with the above results, we succeeded in achieving the best performance with the working memory and gambling tasks using the brain kernel. For other tasks, the brain kernel performed mildly better than the ridge prior and the smoothing prior estimates. The overall performance of these HCP task fMRI datasets indicates that the brain kernel is a better choice than the smoothing and the ridge priors.

**Fig 9.**
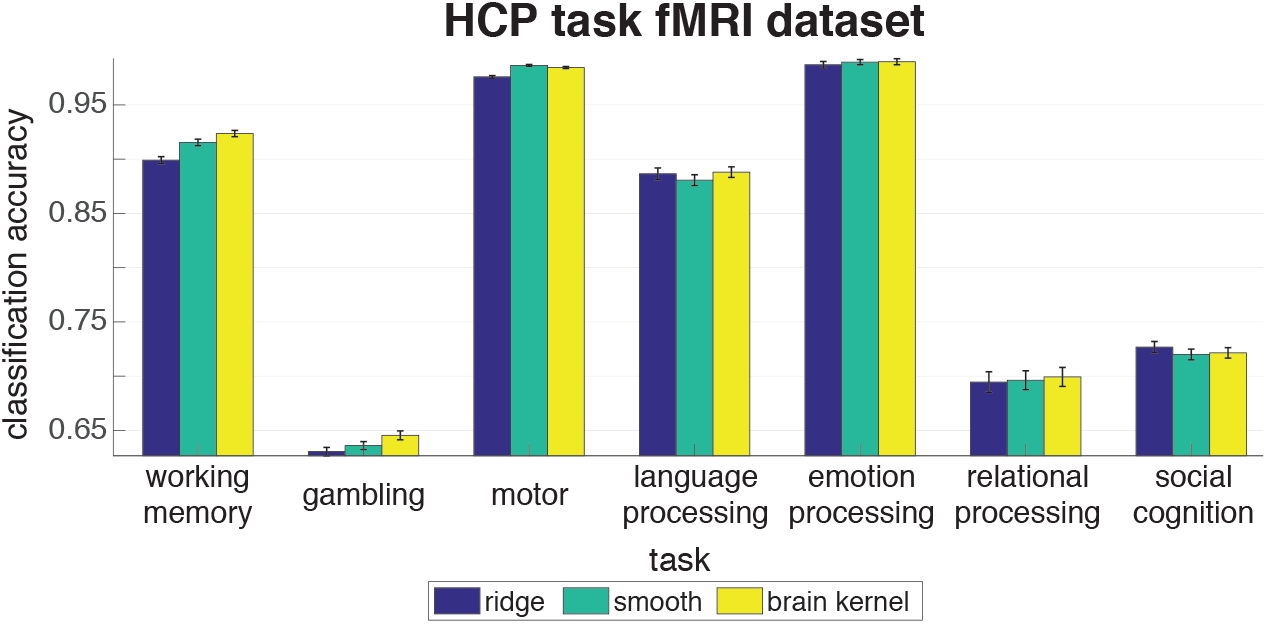
Accuracy performance averaged over all subjects for all task fMRI datasets in the HCP database. The x-axis indicates the task. The y-axis is accuracy performance.

#### 6.1.2 Visual recognition task

Next, we examined the problem of decoding faces and objects from fMRI measurements during a visual recognition task. Just to remind, the brain kernel was estimated using the resting-state fMRI from HCP, and we applied it to a popular fMRI dataset from a study of human ventral temporal cortex [31] for the decoding task. We extended the application beyond HCP, i.e., constructing the brain kernel on HCP and then trying on a completely different dataset. Therefore, we were looking at cross-dataset generalization, not just cross-task generalization within HCP. In this visual recognition experiment, six subjects were asked to recognize eight different types of objects (bottles, houses, cats, scissors, chairs, faces, shoes, and scrambled control images, examples in Fig. 10). Each subject participated 12 scanning runs. In each run, the subjects viewed images of eight object categories, with 11 whole-brain measurements per category. Each subject’s fMRI data was preprocessed using the fMRIprep package^2^ [32] and aligned to the MNI template. Both voxels in this dataset and voxels in the HCP database were all aligned in the same MNI space, allowing us to use eq. 19 to get the brain kernel covariance for the present dataset. Instead of analyzing the whole-brain data, we extracted ROIs with 1645 voxels in the ventral temporal cortex, which is thought to be involved in object recognition. The ROI mask was obtained from Nilearn [33].

**Fig 10.**
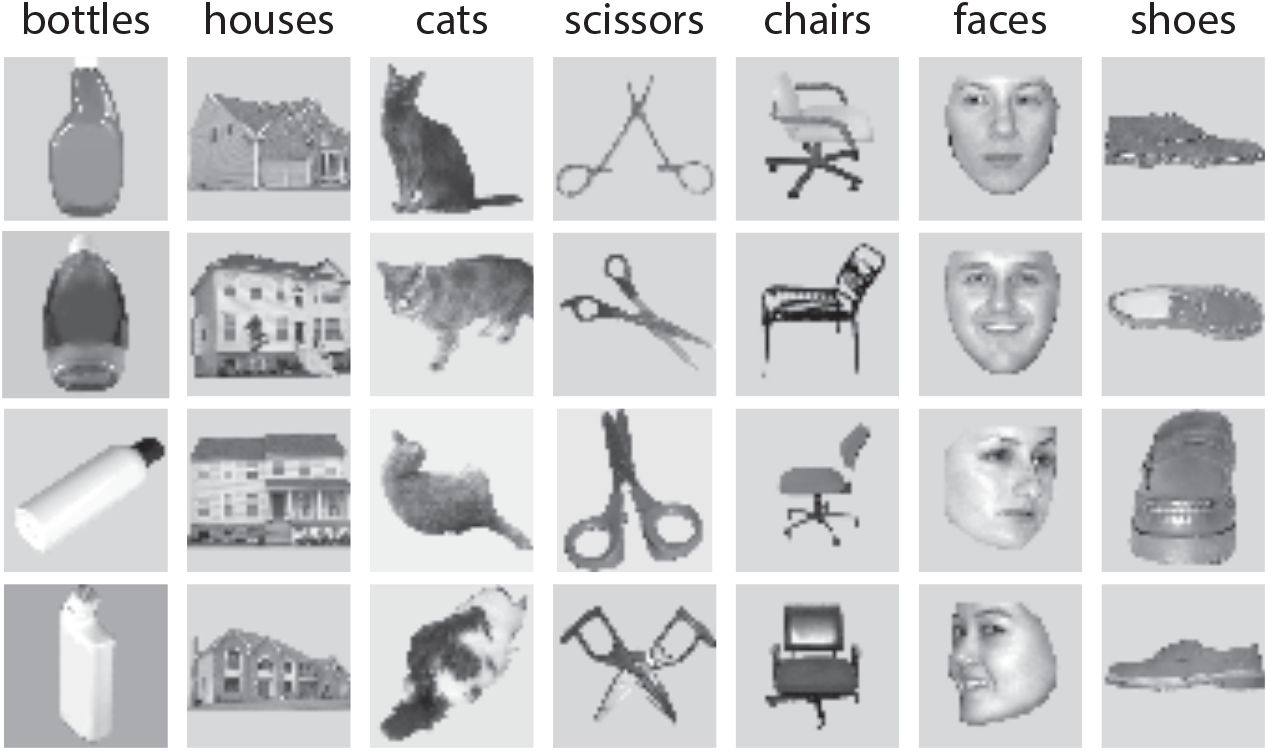
Examples of the stimuli for 7 categories (except for scrambled control images) [31].

We assessed performance by training Bayesian linear regression classifiers to discriminate between pairs of objects, e.g., face vs. bottle, for each of the 28 possible binary classifications among the eight objects (Fig. 10). We trained the weights w for each model using linear regression from fMRI measurements x to binary labels *y* ∈ {−1, +1}, and assessed accuracy on the test set using predicted labels 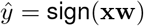. Here, we split the training and test sets by subjects. In this visual recognition dataset, we had six subjects but only 11 × 2 × 12 = 264 measurements in a 1645-voxel space for each subject in a binary decoding task. The number of training measurements was not sufficiently large to train a good classifier with a reasonable generalization performance. Therefore, different from the HCP tasks, we chose to do inter-subject analyses by using 5 subjects for training and one subject for test. We repeated this leave-one-subject-out manner for six times with each subject being used as the test set once and obtained the result in Fig. 11.

**Fig 11.**
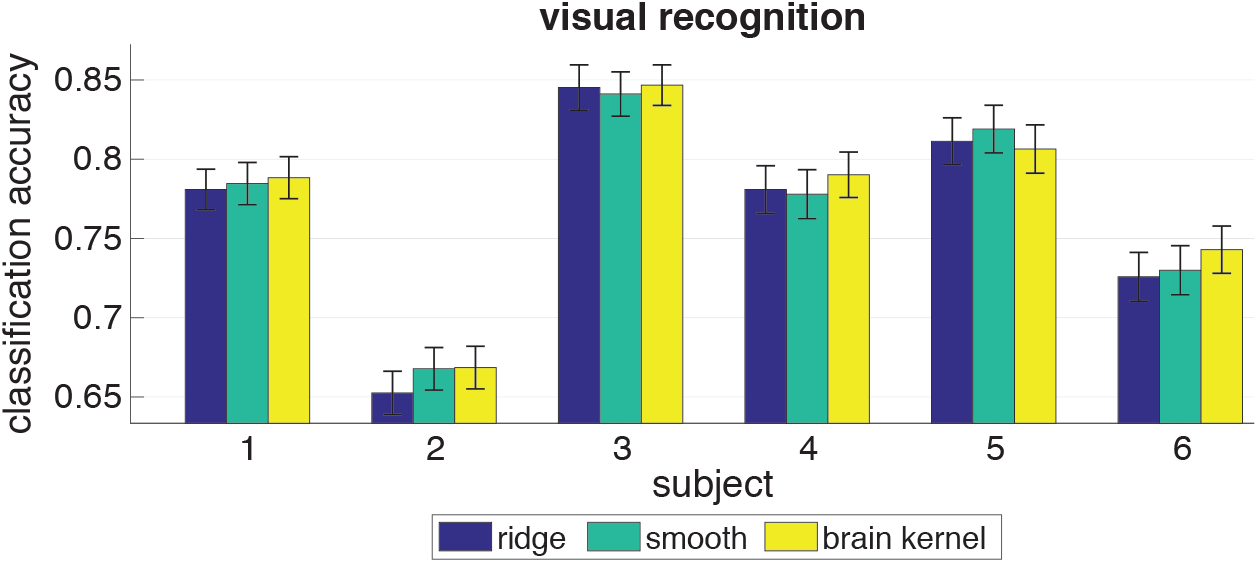
Accuracy performance on the visual recognition task. The x-axis indicates subject IDs. The y-axis is accuracy performance. We compared our brain kernel with a ridge prior and a smooth RBF kernel, color coded.

The averaged accuracy performance across four repeated runs for six subjects shows that the brain kernel performed comparably to the ridge and smoothing priors with better accuracy performance for 5 out of 6 subjects (Fig. 11). This indicates that the brain kernel can provide functional and structural support for most subjects and visual recognition tasks in this dataset. The improvement was statistically modest overall based on the standard errors, which could be a result of several factors: misalignment of the coordinate space to the HCP coordinate space used to estimate the brain kernel; mismatch between the resting-state covariance used to construct the brain kernel and covariance present during the visual recognition task; or the object recognition tasks may rely on fine-grained spatial response topographies that are poorly aligned across individuals.

### 6.2 Factor modeling

In this section, we illustrate a second type of application of the brain kernel. Instead of using it to regularize decoding weights, as in the previous section, we used it as a spatial prior for Bayesian factor analysis.

#### 6.2.1 Sherlock movie watching task

We examined the Sherlock fMRI dataset, in which participants were scanned while they watched the British television program “Sherlock” for 50 min [34]. The fMRI data comprised 1,973 TRs (Repetition Time), where each TR was 1.5 s of the movie. Before performing any analysis, the fMRI data were preprocessed and aligned to MNI space using the techniques described in the prior work [34]. We examined the brain data averaged across all subjects to smooth out individual variability. We identified 11 ROIs previously implicated in processing naturalistic stimuli, comprising the default mode network (DMN-A, DMN-B), the ventral and dorsal language areas, and the primary auditory and visual cortices [35].

For each ROI, we performed a standard factor analysis (FA) to factorize the voxel-by-time fMRI data into a latent source matrix and a factor matrix. Thus fMRI images can be considered as being generated by a covariate-dependent superposition of latent sources. Some following analyses, such as decoding and encoding tasks, can be performed with the factor matrix. The number of latent sources is much fewer than the number of time points, thus providing a parsimonious description of neural activity patterns that avoids many of the pitfalls of traditional voxel-based approaches. Similar FA based models have been proposed in [36, 37]. Here, we performed a Bayesian factor analysis of the form

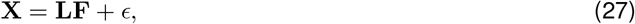

where **X** ∈ ℝ^*N*×*T*^ is the fMRI image with *N* voxels and *T* time points, **L** ∈ ℝ^*N*×*K*^ is the latent source matrix encoding the canonical spatial pattern (over voxels) associated with each latent source. **F** ∈ ℝ^*K*×*T*^ is the factor matrix containing the timeseries for *K* latent sources. *ϵ* is a Gaussian noise with 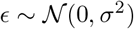. We assumed that the *i*th factor (column) is from a standard normal distribution, i.e., 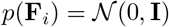. We also assumed that the *j*th latent source (column) is from a Gaussian prior 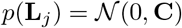 where **C** is the covariance matrix. Like the decoding tasks, we used a ridge prior, a smooth RBF kernel, and the brain kernel in the place of the covariance matrix **C**. With different priors, the latent sources **L** showed different characteristics imposed by these distinct regularizations.

Our goal was to infer the latent variables **L** and **F**, and the hyperparameters for **C** and σ^2^, denoted as *θ*. To quantify performance, we compared data explanation under the three prior with the same number of latent sources which was set to 10. In practice, we first standardized the fMRI image **X**, then tuned the hyperparameters using the marginal distribution of **X**, and finally inferred the factor matrix and the latent sources via maximum likelihood parameter estimation (details in Methods). To evaluate the performance of the three priors, we used a method called “co-smoothing” [38], depicted in Fig. 12. We first split the time points into two equal sets by taking the first half as one set and the latter half as the other set. We took the first set for training (the blue region) and the second set for inference (the pink region) and test (the yellow region). We trained the model with the neural activity in the training set to obtain the estimated latent sources **L**^∗^. We then kept the latent sources fixed for the inference and test purpose. Next we split the second set into five folds along the voxel axis, four for inference and one for test. We used the neural activity in the inference set to infer the factor matrix (mapping the latent sources to the time series) during the inference period given the latent sources **L**^∗^. Finally we predicted the neural activity for the left-out voxels in the test set given the latent sources and factor matrix. We repeated the inference and test step five times in a cross-validation fashion and obtained an averaged *R*^2^ value representing the performance of the prior. After the first run, we achieved three *R*^2^ values for the three priors (ridge, smooth and brain kernel). To better visualize the difference, we normalized the three *R*^2^ values so that the maximum was 1 and the minimum was 0. We then launched a second run by using the second set as the training set and the first set as the inference and test set. The final normalized test *R*^2^ value was an average of the two runs.

**Fig 12.**
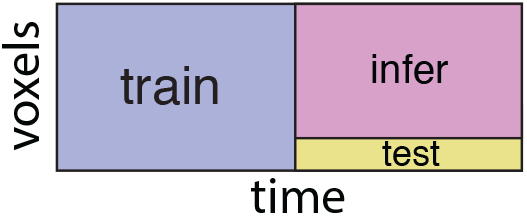
Co-smoothing evaluation. The blue region is the data used for training; the pink region contains voxels that are used to infer the factor matrix during the inference period; and the yellow region is used for test.

We compared the normalized test *R*^2^ value for the three priors for each ROI (Fig. 13). The error bars indicate standard errors. In most regions, the brain kernel outperformed the ridge prior and the smooth RBF kernel. This implies that when performing Bayesian FA, the brain kernel provided a superior prior covariance for the latent source matrix and may enhance performance in terms of data explanation.

**Fig 13.**
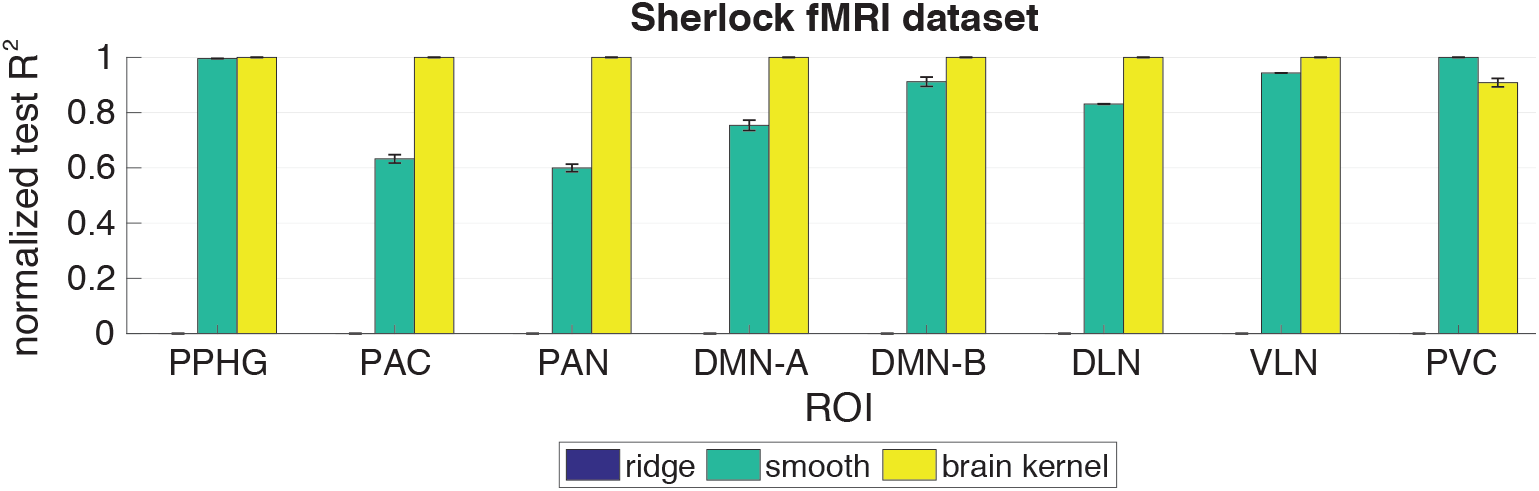
Normalized test *R*^2^ performance on the Sherlock fMRI dataset. The x-axis indicates ROIs (PPHG – Posterior Parahippocampal Gyrus; PAC – Primary Auditory Cortex; PAN – Primary Auditory Network; DMN-A – Default Mode Network-A; DMN-B – Default Mode Network-B; DLN – Dorsal Language Network; VLN – Ventral Language Network; PVC – Primary Visual Cortex). The y-axis is the normalized *R*^2^ performance on the test set (higher values indicate better performance). The error bars indicate standard errors. We compared our brain kernel with a ridge prior and a smooth RBF kernel, color coded.

#### 6.2.2 HCP tasks

We examined the task fMRI datasets in the HCP database with the same Bayesian FA model. We collected all the task fMRIs for the same 10 subjects and performed Bayesian FA for each subject in each task. We implemented the same “co-smoothing” evaluation and compared the normalized test *R*^2^ for the three priors averaged over all subjects for each task (Fig. 14). In most tasks, the brain kernel outperformed the ridge prior and the smooth RBF kernel. This implies that the brain kernel provided a superior prior covariance for the latent source matrix and may enhance performance in terms of data explanation for the HCP database.

**Fig 14.**
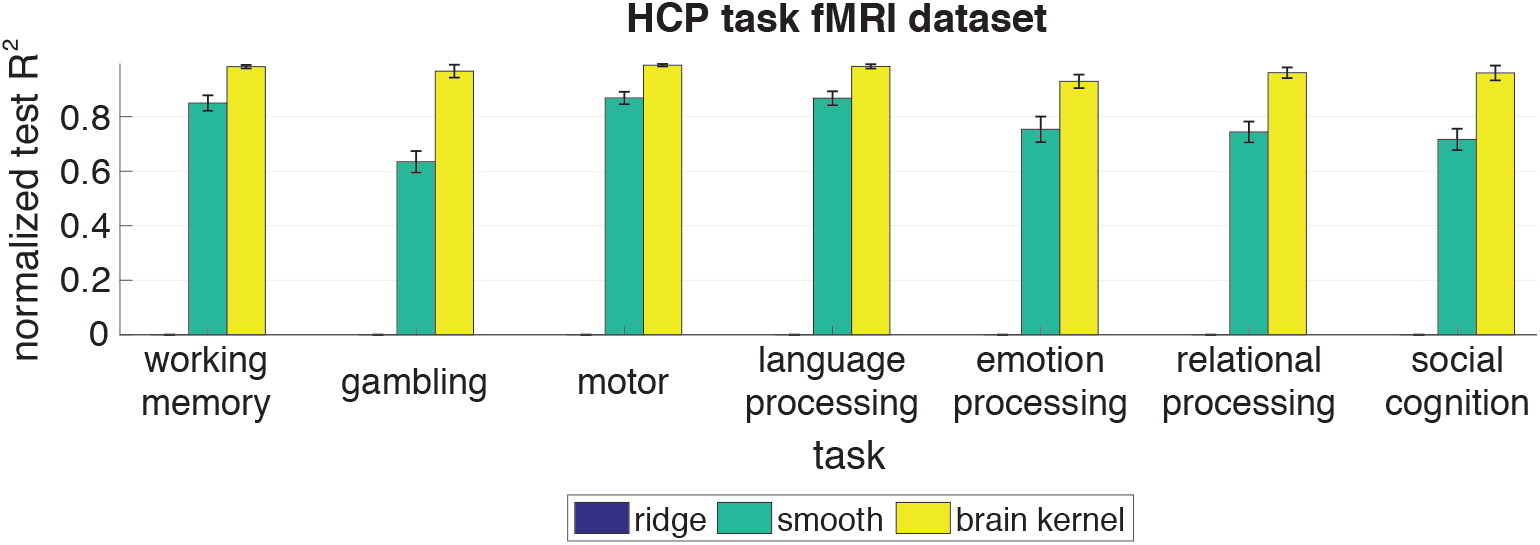
Normalized test *R*^2^ performance on the HCP task fMRI datasets. The x-axis indicates the task. The y-axis is the normalized test *R*^2^ performance averaged over 10 subjects for each task (higher values indicate better performance).

## 7 Discussion

We introduced the brain kernel model, a new model for the spatial covariance of fMRI data. This model takes the form of a covariance function, meaning that it can take any two continuous voxel locations in the brain as inputs and return the prior covariance of their activities. Unlike standard smoothness-inducing covariance functions used in the Gaussian process literature, which assume that correlation falls off monotonically with distance, the brain kernel allows widely-separated voxel locations (e.g., on opposite sides of the brain) to have high correlation, while pairs of nearby voxels can be nearly uncorrelated. This property arises from the fact that the nonlinear embedding function, parametrized with Gaussian processes, warps and stretches the brain in a higher-dimensional space so that widely-separated pairs of voxels can be mapped to nearby embedding locations, while pairs of nearby voxels can be moved far apart in embedding space.

We fit the brain kernel model using a nonlinear embedding of the 3D brain in a 20D space. We introduced an exact inference method for fitting the brain kernel model using block coordinate descent (BCD), and estimated the brain kernel using a large publicly-available dataset of resting-state fMRI measurements from many subjects. We found that the resulting brain kernel accurately captured the covariance of resting-state fMRI measurements.

We further demonstrated the applicability of the brain kernel using two different modeling tasks: brain decoding and factor analysis. In both cases, we used the brain kernel to define a prior distribution in place of a more conventional prior based on *ℓ*_2_ shrinkage or local smoothness. We showed that the brain kernel achieved gains in performance, illustrating that the correlations in resting-state fMRI may be usefully mined to aid analyses of task fMRI. A comprehensive summary of application results can be found in Appendix B.

Moreover, to examine other choices of covariance function for parametrizing the brain kernel, we re-fit the brain kernel to resting-state fMRI data using a Matern-3/2 covariance function. We also compared the RBF and Matern-3/2 brain kernels using the working-memory decoding task fMRI data shown in the main paper. We found that the RBF covariance function outperformed the Matern-3/2 covariance function in both modeling the covariance of resting-state data and decoding of working memory task data. More details can be found in Appendix C. We have therefore decided to keep our focus on the brain kernel parametrized with RBF covariance in the main paper. But exploring a wider family of possible covariance functions is indeed a worthwhile direction for further research.

The estimation of a good quality of the brain kernel requires as much fMRI data as possible. However, we want to emphasize is that our intention was not to suggest that experimentalists should collect more data than needed for their study. We merely meant to suggest that—if there are pre-existing publicly or privately available datasets that exhibit similar functional connectivity / covariance structure to patterns of activity observed during a particular study, then that additional data might be leveraged *using the brain kernel* to improve decoding performance or latent variable modeling of the data collected for the study of interest. This would indeed incur a computational burden (researchers would need to fit the brain kernel using the pre-existing dataset). However, no additional data would be needed; on the contrary, the brain kernel would make it possible to exploit the structure of the pre-existing data so that the significance level / effect sizes are larger in the study of interest, thus effectively increasing statistical power or reducing data requirements. For the purpose of using the resting-state brain kernel as an alternative prior apart from the RBF covariance, people can just download it from [link](https://github.com/waq1129/brainkernel/blob/main/brainkernel/brainkernel_latent.matlink) and use it instead of the RBF covariance, since we have already fit it with a giant resting-state fMRI dataset from diverse subjects.

### Limitations

Despite the successes presented above, it is important to acknowledge that we applied the brain kernel to several other datasets for both the decoding and the factor modeling tasks beyond the examples shown in the main section, for which it did not yield improvements over standard priors.

For decoding, we applied the brain kernel to the Sherlock movie dataset [34] whose decoded vectors contained semantic descriptions for the movie scenes. We observed that the smoothing prior was better than the brain kernel prior, both better than the ridge prior. Then we investigated the reason of the performance difference and figured out that the brain kernel was able to impose a strong cross-covariance assumption over many voxels; but much of the cross-covariance didn’t exist in the Sherlock movie data or could hurt the decoding performance. The optimal brain kernel prior corresponded to a small length-scale that led to less smoothing assumptions compared with the smoothing kernel. So this Sherlock movie dataset is not an ideal dataset that could leverage most of the resting-state functional connectivity to do the decoding task. We also applied it to a public fMRI dataset in which subjects viewed 5000 visual images (BOLD5000) [39]. The binary labels were living objects versus nonliving objects. We observed that the brain kernel did not reliably outperform the ridge prior and the smoothing prior estimates across subjects. The optimal smoothing prior and the brain kernel prior both converged to the ridge prior after hyperparameter estimation, i.e., the estimated length-scale was very small and all three priors had the same performance. Therefore, smoothness did not help the classification task. More details can be found in Appendix B. For these cases, regularizing the decoding weights with the brain kernel did not improve performance, suggesting that the covariance of resting-state fMRI did not provide useful information for classifying fMRI measurements in these tasks, or at least that the brain kernel was not capable of capturing it. It remains possible, however, that a brain kernel fit from task fMRI datasets might offer benefits for decoding fMRI data from these tasks.

For factor modeling, we also applied the brain kernel to the visual recognition task and the BOLD5000 dataset except for the HCP dataset and the Sherlock movie dataset presented in the main paper. For most subjects in both datasets, the brain kernel didn’t show a dominating performance over both the ridge prior and the smooth RBF kernel (Appendix B).

Beyond those reasons, we hypothesize that the lack of benefit observed on non-HCP (outside the HCP) datasets may arise from a mismatch between the covariance of resting-state fMRI observations used to fit the brain kernel and the covariance of task fMRI datasets. However, it might also arise from differences in acquisition parameters, preprocessing steps, or alignment between HCP and non-HCP datasets. Although we aligned all voxels with the MNI template, misalignment might still result from differences between processing pipelines.

We include more analyses about the reasons behind these limitations and under which circumstances the brain kernel could be a better prior option than the ridge prior and the smoothing prior in Appendix B. We can show only that if there is similar structure between the covariance of resting-state data used to estimate the brain kernel and the discriminative ROIs in the task data, the brain kernel will function as a better prior than a standard smoothing kernel. For factor modeling applications, the brain kernel is often more reliably effective because we typically use a large number of voxels to infer latents, rather than a few responsive voxels that may be selected in decoding tasks. Similarly, we would expect the brain kernel to give good performance when the task fMRI is composed of smooth latent sources that resemble the spatial correlation in the resting-state data. We can fit a best brain kernel or a best RBF kernel to the sample covariance and evaluate the similarity both qualitatively and quantitatively with the negative log-likelihood (NLL) value. If the best-fitting brain kernel resembles the sample covariance and the NLL value of the brain kernel is smaller, it’s promising to use the brain kernel as a prior for Bayesian factor modeling analysis.

Given that the computational burden is not higher than that of standard smoothing or shrinkage priors (and potentially smaller than shrinkage-inducing regularizers like LASSO), we hope that researchers will incorporate the brain kernel into standard analysis pipelines, and apply it in cases where it is observed to offer improved performance. We consider this to analogous to the ways in which existing regularizers such as smoothing priors or sparsity-inducing priors like LASSO are currently employed: researchers may conduct exploratory analyses to determine whether incorporating smoothness or sparsity improves performance, and then adopt these regularizers as warranted by the data.

### Outlook and future directions

Although the brain kernel did not achieve improved performance in all datasets and applications we considered, we feel it nevertheless holds great potential for fMRI data analysis. First, we cast the problem of estimating covariance of fMRI datasets as that of estimating a nonlinear mapping from 3D brain coordinates to a latent embedding space. This results in a compact representation of the full brain covariance matrix, requiring storage of only a *N_voxels_* × 20 matrix of embedding locations (when the embedding dimensionality is 20), as opposed to a full *N_voxels_* × *N_voxels_* covariance matrix. Moreover, because the brain kernel is a continuous covariance function, we can use it to model the covariance at arbitrary voxel locations, even those not contained in the original training dataset.

In addition to providing a compact parametrization of fMRI covariance, the brain kernel’s nonlinear embedding function may be useful for gaining insights into the geometry of correlations within and across brain regions. Examining the embedding of different brain regions (e.g., as shown in Fig. 4) allows researchers to directly visualize correlations in terms of distances between embedded voxels.

Although the brain kernel fit to the HCP resting-state fMRI data did not yield improvements on all task fMRI datasets we explored, it is possible that other methods for training or applying the brain kernel might produce bigger gains. For example, one might train the brain kernel on task fMRI datasets, or train a hierarchical version that gains statistical strength from combining datasets, while preserving flexibility to capture differences between the two kinds of data. A more ambitious possibility is to formulate a hierarchical version of the brain kernel that allows for per-subject variability. This would produce a hierarchical brain kernel in which each brain has own specific brain kernel, allowing for detailed differences in correlation maps across brains. All such applications will benefit from more robust methods for alignment and preprocessing, and it may be that these alone will be enough to improve the performance of the brain kernel. Although these directions are beyond the scope of the current paper, we feel that the idea of the brain kernel is one that researchers might extend and apply to novel settings and datasets.

Finally, an exciting possibility for future work is to combine the brain kernel with other advanced statistical modeling techniques. Methods for modeling structured variability, such as topographic ICA [37], and methods for structured sparsity, such as GraphNet [40], sparse overlapping sets lasso [41], and dependent relevance determination [42,43], rely on capturing dependencies between nearby voxels. All such methods might thus be improved by using nearness in functional embedding space provided by the brain kernel, instead of nearness in 3D Euclidean space inside the brain. Likewise, methods for linear alignment of functional data from multiple subjects such as hyperalignment [44, 45] might be extended using nonlinear warping of brain coordinates under the brain kernel. Therefore, we feel that the brain kernel holds promise for inspiring new data-driven prior distributions and new modeling approaches for capturing structure in fMRI data.

## Methods

### Block coordinate descent for the brain kernel model

The penalized least squares (eq. 18, PLS) has a computational complexity of *O*(*n*^2^) and memory storage of *O*(*n*^2^); however, the maximum a posteriori (eq. 15, MAP) has a computational complexity of *O*(*n*^3^) – to invert the covariance matrix – and memory storage of *O*(*n*^2^). *n* is the number of voxels, which often exceeds 10K. Gradient descent or Newton’s method is computationally impractical as the optimizer. Thus, we need to use a scalable inference method. Existing inference methods for large datasets [15–17] exploit low-rank approximations to the full Gaussian process, which, however, suffer from a loss of accuracy in covariance estimation. Thus, in this section, we develop a block coordinate descent algorithm as an exact inference method for the brain kernel model. Coordinate descent has been successfully applied to solve penalized regression models [18], to estimate covariance graphical lasso models [19], and to compute large-scale sparse inverse covariance matrices [4].

Our PLS and MAP estimators are non-convex smooth functions. We apply an iterative block coordinate descent method solved by the proximal Newton approach [20]. We first divide the voxel set {1,…, *n*} into blocks. Next, we iterate over all blocks, minimizing the functions with respect to the voxels within each block. Without loss of generality, we split the voxel set into two blocks, {1,…, *m*} (block 1) and {*m* + 1,…, *n*} (block 2), where *m* ≪ *n*, and focus on the first *m* columns of **Z** for the update. We partition **Z, C, S**, and **K**^−1^ as follows:

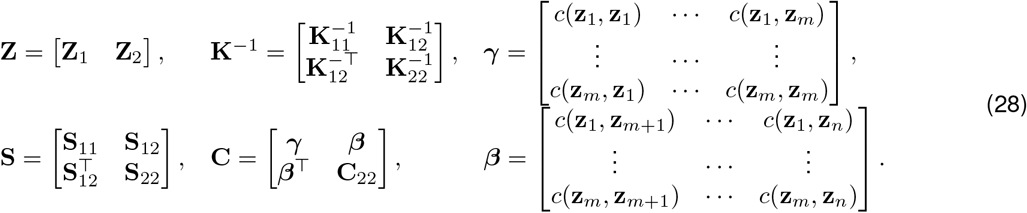

Here, subscripts represent the block indices, so **Z**_1_ and **Z**_2_ are the first *m* columns and the last *n* − *m* columns of **Z** and subscript 12 indicates the block matrix across the first *m* variables and the last *n* − *m* variables. Only *β* and *γ* contain the active variables in **Z**_1_ to optimize. **Z**_2_ is fixed.

Applying the block representation to eq. 18, we get the block PLS objective function for solving **Z**_1_:

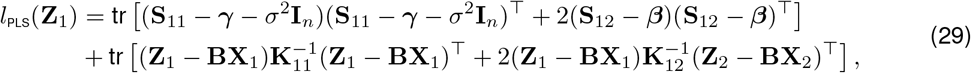

where **X**_1_ and **X**_2_ are the first *m* columns and the last *n* − *m* columns of **X**.

To formulate the block MAP estimator for **Z**_1_, we first apply the block matrix inversion to **C**,

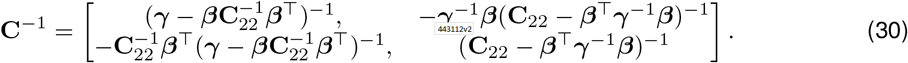

Incorporating this matrix inversion into the MAP estimator, we get the objective function

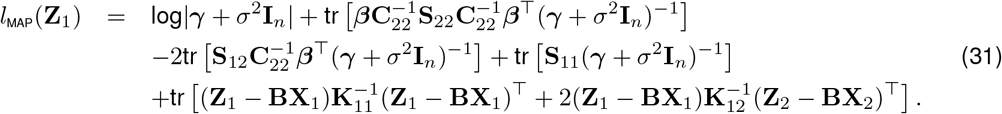

The time complexity of the block PLS estimator is *O*(*nm*^2^) per iteration, where *m* is the size of the block, and the time complexity of the MAP estimator is *O*(*n*^2^*m*) per iteration. For comparison, greedy gradient descent has complexity *O*(*n*^3^) per iteration. In the experiments, the block MAP estimator for Z with the block PLS estimator as a warm start is a practical optimization approach.

We now describe the block coordinate descent (BCD) update. We assume that the voxel indices {1,…, *n*} are divided into *k* blocks 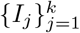, where *I_j_* is the set of indices corresponding to the columns of **Z** in the *j*’th block. Denote *I_j_* columns of **Z** to be 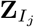. We cluster indices into blocks based on the spatial locations of the voxels and assume smooth measurements for nearby voxels. Thus, the size of a block should be at least one length-scale of the region defined by the kernel in x space to encourage dependencies among neighboring voxels. This smoothness assumption leads to a block-wise but not an element-wise update, which separates our BCD method from Informative Vector Machine (IVM) [46]. At each iteration *t*, we choose a nonempty index subset 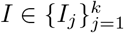. Then the objective functions *l*(**Z**_*I*_) in eq. 29 and 31 are optimized w.r.t. **Z**_*I*_ via L-BFGS [47]. After the t’th iteration, **Z**_*I*_ is updated, holding all other blocks fixed.

For high-dimensional non-convex problems, a good initialization is essential to finding a good optimum in practice. Two steps we exploited during implementation to mitigate the optimization issue with multi-modal, high-dimensional non-convexity.

First, we assumed the nonlinear latent embedding **z** to be a local warping of the linear embedding which is the mean of the posterior distribution for **z** (eq. 8). We first found a good estimate of **b** at the beginning of the optimization. This is equal to fitting a linear brain kernel (LBK) model. We estimated **B** in eq. 22 which is a matrix form of **b**. This involves an easier optimization since the parameter space of **b** is much smaller than **z**. Given the estimated **b**, 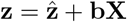 is optimized via estimating 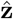 and fixing **b**. This eases the high-dimensional non-convex issue by learning a small parameter **b** and a local warping 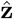 separately.

Secondly, we introduced the Laplacian eigenmap algorithm, an effective and tractable initialization for a single block of **Z** inspired by Laplacian eigenmaps.

The Laplacian eigenmap (LE) algorithm is a popular dimensionality reduction method that solves a generalized eigendecomposition [48]. LE defines a neighborhood graph on the data 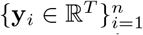, such as *k* nearest neighbors or an *ϵ*-ball graph, and weighs each graph edge *y_i_* ~ *y_j_* by a symmetric affinity function *V* (*y_i_*, *y_j_*) = *v_ij_*, typically Gaussian: 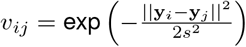 with *s* the length-scale. Given this weighted graph, LE seeks latent points 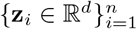 that are solutions to the optimization problem

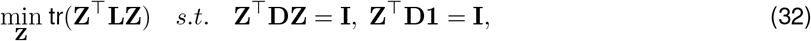

with the symmetric affinity matrix **V** ∈ ℝ^*n*×*n*^, the degree matrix 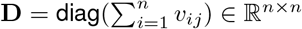, the graph Laplacian matrix **L** = **D** − **V** ∈ ℝ^*n*×*n*^, and **1** = [1,…, 1]^⊤^. Constraints eliminate the two trivial solutions **Z** = 0 by setting an arbitrary scale and **z**_1_ = … = **z**_*n*_ by removing 1, which is an eigenvector of **L** associated with a zero eigenvalue.

Following the previous two-block example, let **Z** be partitioned into **Z**_1_ and **Z**_2_. To update **Z**_1_ given **Z**_2_, the objective function is:

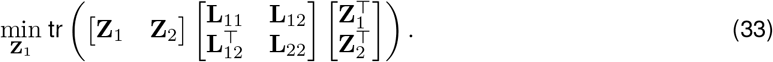

We don’t need to use the constraints from eq. 32 because the trivial solutions are removed given **Z**_2_. The solution is

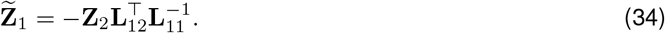

Given the current latent embeddings for all other coordinates, the algorithm seeks the best latent embedding of the unknown dimensions. Because of the computational efficiency of this approach, we use 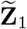 to initialize the BCD algorithm for each iteration and **Z**_2_ collapses all the fixed blocks.

In addition, if **V** has a Gaussian affinity function, the latent embedding **Z** is mapped from the observation **Y** with a radial basis function nonlinearity. For covariance estimation, we enforce the resemblance between another radial basis function (RBF) kernel on **Z** and the sample covariance of **Y**. We are essentially trying to map the observation space to itself with double layers of exponential transformations, which would result in a bad latent embedding for initialization. Therefore, instead of using a Gaussian function for the weights **V**, we use a function that inverts the RBF nonlinearity on a covariance matrix, defined as 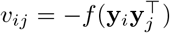. Here, *f*(*x*) = sign(*x*)log(|*x*| + 1) is the log-modulus transformation [49], which distributes the magnitude of the data while preserving the sign of the data in order to control against negative covariance values when taking the logarithm.

We chose the Laplacian eigenmap algorithm because it has the nice closed-form expression for the conditional expression **Z**_1_ given **Z**_2_. We were able to use **Z**_2_ to efficiently find a better initial position for **Z**_1_ that reduced the search over the entire parameter space of **Z**_1_. We tried a random start for **Z**_1_ which led to a very bad local optimum. We also tried to use the previous estimate to initialize **Z**_1_. That resulted in a stuck in a bad local optimum that was very close to the previous solution. So Laplacian eigenmap allowed us to move away from the previous estimate but also leverage **Z**_2_ effectively.

### Hyperparameter estimation for the brain kernel model

Our model includes five hyperparameters: {*δ, r*, **B**, *ρ*, σ^2^}. We set these parameters as follows.

#### Estimate *δ*

*δ* is the length-scale of the GP kernel mapping from coordinate space to latent space. We can optimize this parameter by taking the derivative of the GP log-likelihood w.r.t. *δ*; this involves inverting the kernel matrix with an *O*(*n*^3^) cost, which is computationally infeasible here. An inducing-point method [16] introduced extra inducing variables to optimize, which further increased the computational burden. Instead, we use a scalable spectral formulation for learning the length-scale *δ* of the GP kernel.

For a stationary Gaussian process 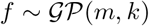, when *f* has a high degree of smoothness, the prior covariance **K** becomes approximately low rank, meaning that it has a small number of non-negligible eigenvalues. Because the kernel function for x space is shift invariant, the eigenspectrum of **K** has a diagonal representation in the Fourier domain, a consequence of Bochner’s theorem [50, 51],

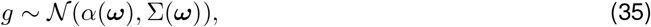

where *g* is the Fourier transform of *f*, *ω* is the frequency, *α*(*ω*) is the Fourier transform of the mean function *m*(x), and Σ = diag(*s*(*ω*)) is diagonal. This means that Fourier components are a priori independent, with prior variance

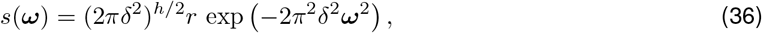

where *h* is the dimension of the input **x**. Without loss of generality, we assume the size of the spectral domain for each input dimension is *w*, and thus 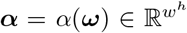 and 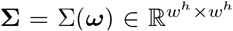 given a spectral representation *ω*. The mapping between *f*(x) and its Fourier transform *g*(*ω*) is then

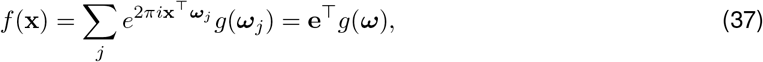

where e is a column vector with entries 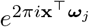 on the *j*’th position for the *j*’th frequency. Let 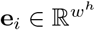 denote the Fourier vector for x_*i*_ (*i* is the index for voxels), then we can further define 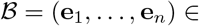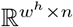 to be the Fourier matrix for an input coordinate matrix **X**. Let **z**_*j*_ ∈ ℝ^*n*^ denote the *j*’th latent embedding of **X** and *j* ∈ {1,…, *d*}. Note that **z**_*j*_ is the *j*’th row of the matrix **Z**. We can write 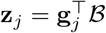, where 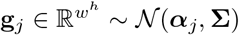. This implies that each latent embedding deterministically depends on a unique spectral function. These spectral functions are all sampled from multivariate Gaussian priors with different mean functions but the same covariance function in the spectral domain. Bringing the Fourier matrix 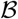 into the Gaussian prior (eq. 35), we derive the Gaussian prior over the latent **z**_*j*_ expressed with the spectral formulation as

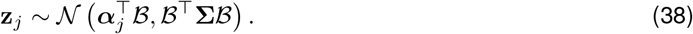

We can use this representation to optimize *δ*. If *δ* is in a large-value regime, we can ignore Fourier components above a certain high-frequency cutoff which leads to a lower-dimensional *ω* and a lower-dimensional optimization problem. Because *h* = 3 in fMRI analyses, we can control *ω* to be small enough so that *w^h^* is tractable relative to the large *n*, and 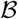 is a manageable high-dimensional matrix. Because we generally do not assume uniform gridding of coordinates **X**, 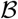 is a non-uniform DFT (not orthogonal). Although the Kronecker tricks cannot be used for scaling up with non-uniform DFT, we can employ block matrix inverse lemma to transform the spectral formulation into a lower-dimensional problem.

Then the standard negative GP log-likelihood to optimize for *δ* is

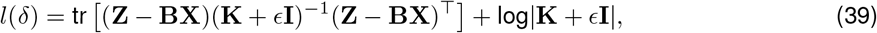

where *ϵ* is a small noise variance in the latent space to avoid ill-conditions for **K**. Let 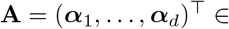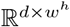. With the spectral representation, eq. 39 can be re-written as

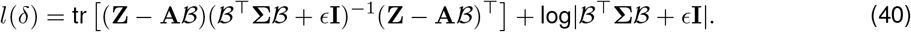

**Algorithm 1.**
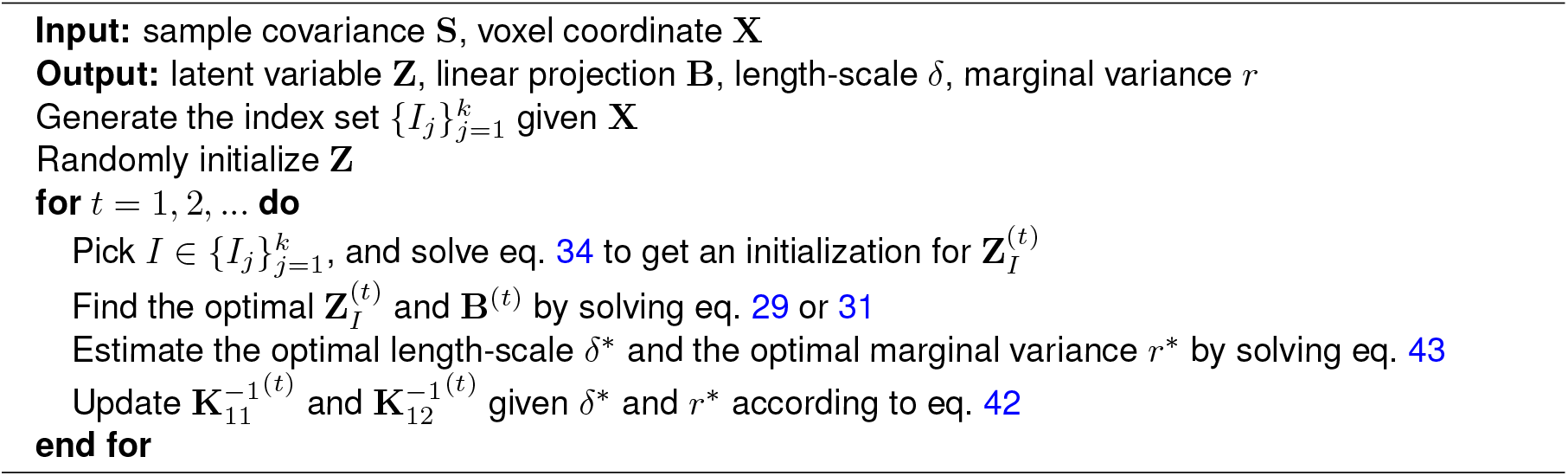
Block Coordinate Descent Algorithm for the Brain Kernel Model

Calculating 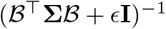 is still computationally expensive, but with the spectral factorization, we are able to use the block matrix inverse lemma as in eq. 30,

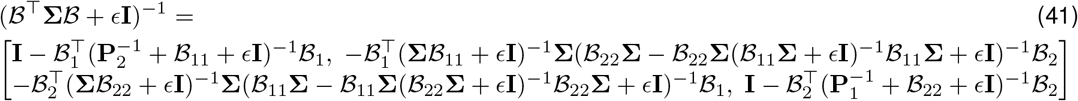

where 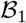 and 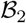 correspond to the Fourier bases for **X**_1_ and **X**_2_ respectively. Note that neither 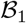 nor 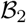 is invertible. Moreover, 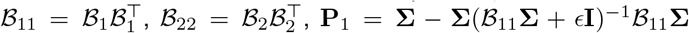 and 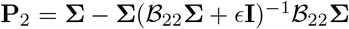. All matrices have size *w^h^* × *w^h^*, which is tractable to invert. We know that the spectral expression of *K*^−1^ is 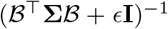. We can also express 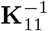 and 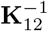 with the spectral formulation as

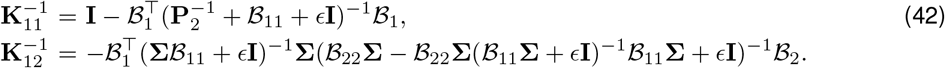

Consequently, the negative GP log-likelihood w.r.t. *δ* in the block form is represented as

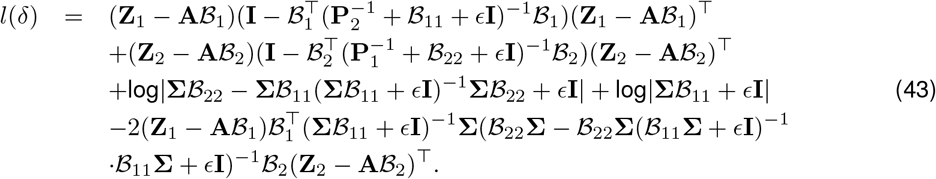

With eq. 43, the computational cost is reduced to max(*O*(*nw^h^*), *O*(*w^3h^*)), where *w* is the dimension of the spectral form per input dimension and *h* is the number of input dimensions. This complexity is much smaller than *O*(*n*^3^) when *n* ≫ *w* and *h* ≤ 3. Another benefit of this formulation is that 5 only exists in the diagonal of Σ (eq. 36), which makes the estimation straightforward.

#### Estimate *r*

*r* determines the scale of the latent embedding **Z**. Since *r* also exists on the diagonal of Σ (eq. 36), the optimization for *r* can be combined with learning *δ* using the same spectral representation in eq. 43.

#### Estimate *ρ*

*ρ* is the marginal variance of the covariance function. We assume that the data has been normalized to have zero mean and variance one. Thus we set *ρ* = 1.

#### Estimate B

**B** is the linear projection of the mean function. We can estimate **B** jointly with **Z** in each BCD iteration by optimizing the same objective function (eq. 29 and 31).

#### Estimate σ^2^

σ^2^ is the observation noise variance. We estimate σ^2^ using the eigenvalues of the sample covariance **S** [52].

Algorithm 1 describes the complete BCD algorithm for the brain kernel model. The convergence of the BCD algorithm (without parameter estimation) to a stationary point is addressed in the theoretical results in previous work [20]. There, a general block-coordinate-descent approach is analyzed to solve minimization problems of the form *F*(*x*) = *f*(*x*) + *λh*(*x*), which is composed of the sum of a smooth function *f*(·) and a separable convex function *h*(·), in our case *h*(*x*) = 0. According to Part (e) of Theorem 4.1 in [20], if *I_t_* at the *t*’th iteration is chosen by the generalized Gauss-Seidel rule,

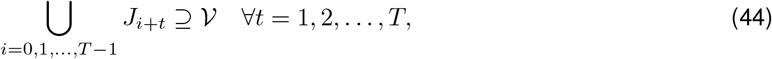

which is necessary to ensure that all variables are updated every *T* steps and 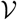 is the set of all variables, then each coordinate-wise minimum point of 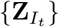 is a stationary point of *f*(**Z**).

### Hyperparameter optimization and latent variable inference for Bayesian factor analysis

Our goal was to infer the latent variables **L** and **F** and the hyperparameters for **C** and σ^2^, denoted as *θ*. We assumed that

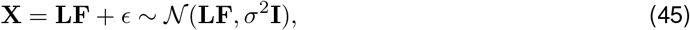

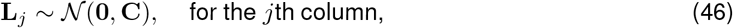

and

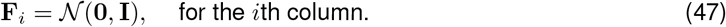

We first aimed at estimating the hyperparameters *θ* by marginalizing over **L** and **F**. The marginal distribution for **X** is

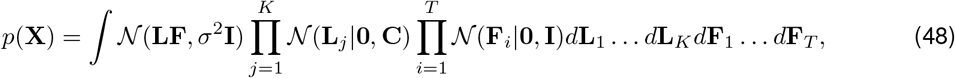

which is intractable. However, we could match the second order moment of **X** to the sample covariance 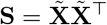 where 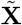 is the standardized data sample. The second order moment is derived as follows

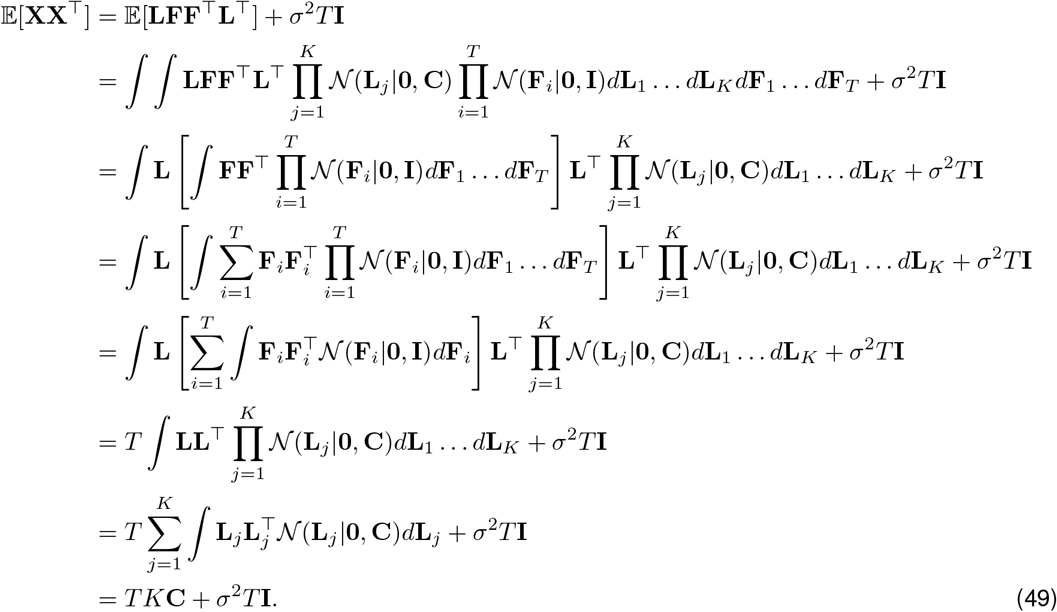

By matching eq. 49 to the sample covariance **S**, we were able to estimate the hyperparameters in **C** and σ^2^.

Next, we fixed the estimated hyperparameters and inferred **L**. Marginalizing over **F** we got

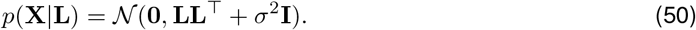

Then, we obtained the joint distribution between **X** and **L** as

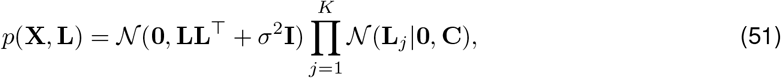

whose log likelihood is written as

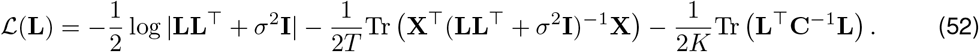

We maximized eq. 52 w.r.t. **L**. To speed up the optimization process, we initialized **L** via finding a closed-form solution approximately maximizing eq. 52. Denoting **LL**^⊤^ + σ^2^**I** as **Q**, we rewrote the log likelihood as

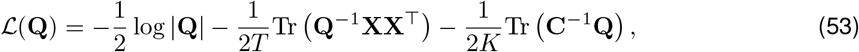

whose derivative is

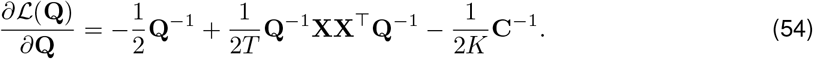

Setting the derivative to be 0, we arrived at

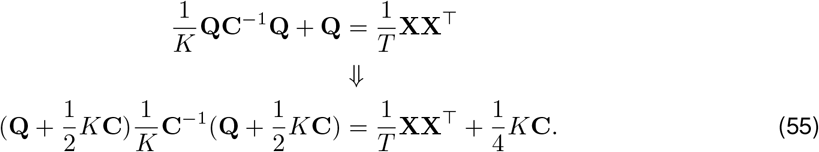

We needed to decompose the right-hand side into the multiplication of a symmetric matrix, **C**^−1^ and the same symmetric matrix. Here is our solution:

- Let **P** denote the Cholesky decomposition of 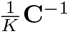, i.e., 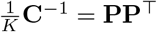.
- Denote 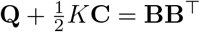 and let **B** = **P**^−⊤^ **A** where **A** is an unknown square matrix.
- Then we can rewrite eq. 55 as

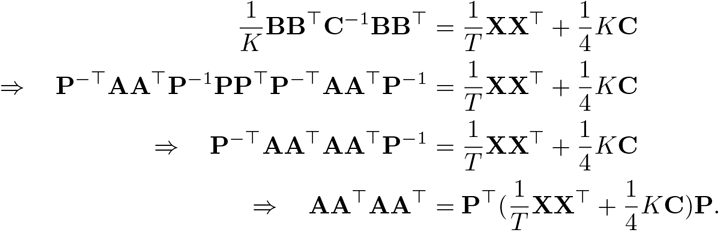
- Next, we represent **A** with its singular value decomposition (SVD), i.e., **A** = **USV**^⊤^.
- Finally we can rewrite eq. 55 as

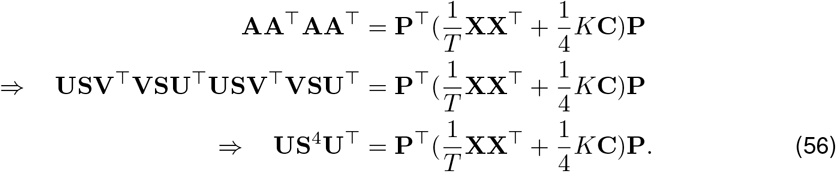
- Therefore, to solve eq. 55, we first factorize 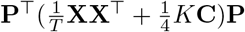 using SVD to obtain **U** and **S** according to eq. 56.
- Then 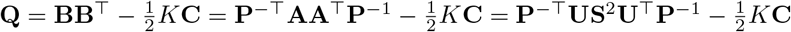.
- Ultimately, **L** can be obtained via 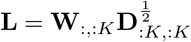 where **WDW**^⊤^ is the eigen-decomposition of **Q** − σ^2^**I**.

Given the above procedure, we were able to find an ideal initialization for **L** which made the optimization of eq. 52 much easier. Conditioned on the optimal **L**^∗^, we turned to inferring the optimal **F**^∗^ via maximizing

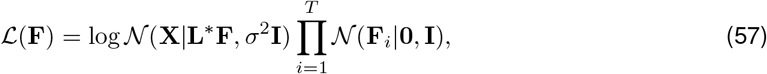

which has a closed-form solution, i.e., **F**^∗^ = (**L**^∗⊤^**L**^∗^ + σ^2^**I**)^−1^**L**^∗⊤^**X**.

Note that in order to obtain 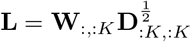, we needed to guarantee that we were able to find a positive semi-definite (p.s.d.) matrix **Q** − σ^2^**I**. Here are the lemma and the theorem:

#### Lemma

*If* **A** *is p.s.d., and* **B** *is symmetric p.s.d., then* **AB** *is also p.s.d.*.

*Proof.* If **A** and **B** are both p.s.d. and B is also symmetric, then suppose λ is an eigenvalue of **AB** with corresponding eigenvector x ≠ 0, i.e., **ABx** = λx. Then **BABx** = λBx and so x^⊤^**BABx** = λx^⊤^**Bx**. It is not hard to check that **BAB** will also be p.s.d.. For x s.t. x^⊤^**Bx** ≠ 0, we have 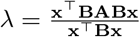. Both the numerator and the denominator are non-negative values, therefore λ ≥ 0. For x s.t. x^⊤^**Bx** = 0, we assume x = **Ve** where **B** = **VDV**^⊤^ is the eigen-decomposition and e is the linear weight vector. Then x^⊤^**Bx** = e^⊤^**V**^⊤^**BVe** = e^⊤^V^⊤^**VDe** = e^⊤^**De** = 0. Since **D** is a diagonal matrix with non-negative values, e should have zero elements corresponding to the non-zero eigenvalues. Therefore x is a linear combination of eigenvectors of **B** whose eigenvalues are zero, i.e., **x** = **V**_0_e where **V**_0_ contains all zero eigenvectors with **D**_0_ = 0 and e has no zero elements. Following that, we have **Bx** = **BV**_0_e = **V**_0_**D**_0_e = 0 ⇒ **ABx** = λ_**B**_**Ax** = 0 ⇒ λ = 0. Based on the derivation, we arrive at the conclusion that **AB** has non-negative eigenvalues thus **AB** is p.s.d..

#### Theorem

*If KC* − σ^2^**I** *is p.s.d., then* **Q** − σ^2^**I** *is p.s.d. based on the **Lemma***.

*Proof.*

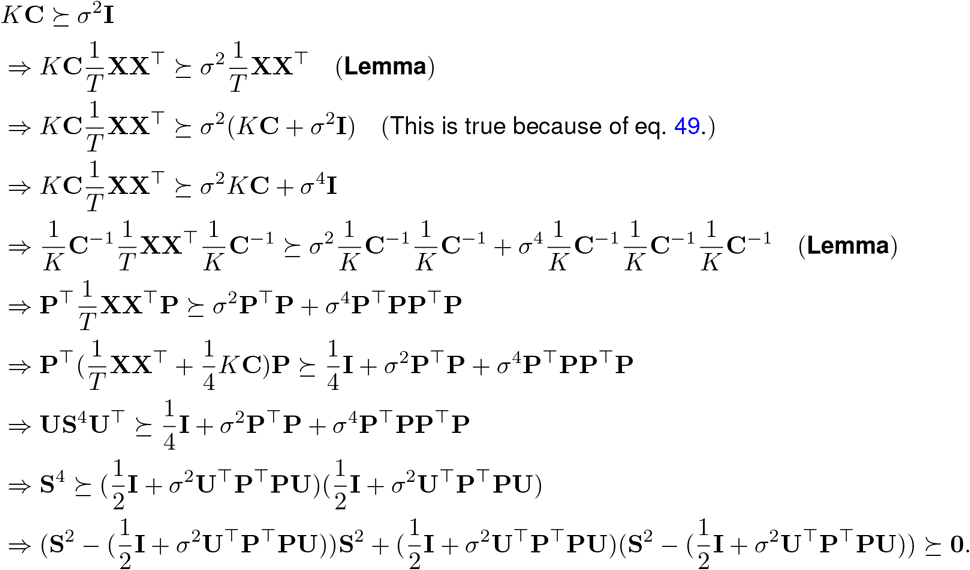

Denoting 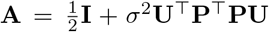 and 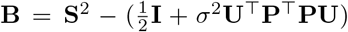, we can simplify the above inequality as

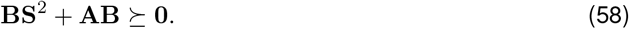

We now prove that if inequality 58 is true, then **B** ≽ **0**, using proof by contradiction:

If **B** ⋡ **0**, then there exists an eigenvector x s.t. **Bx** = λx, λ < 0. Then **x**^⊤^**BS**^2^**x** = λ**x**^⊤^**S**^2^**x** ≤ 0 due to the p.s.d. of **S**^2^. Similarly, for the same eigenvector x, we could arrive at the same conclusion for **AB**, i.e., x^⊤^**ABx** = λ**x**^⊤^**Ax** ≤ 0. This implies that x^⊤^(**BS**^2^ + **AB**)**x** ≤ 0 which is contradict to inequality 58. Therefore **B** ≽ 0. Now we could continue the deduction as follows,

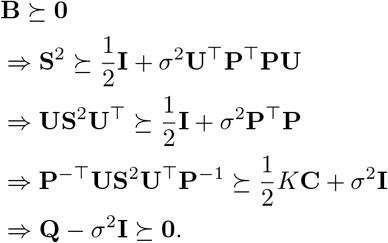

We can easily ensure that *KC* − σ^2^**I** is p.s.d., which guarantees that **Q** − σ^2^**I** is p.s.d. according to the **Theorem**. Therefore 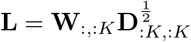 is valid.

## Code Availability

The code is available at https://github.com/waq1129/brainkernel.

## Data Availability

The HCP datasets are publicly available at https://www.humanconnectome.org/. The visual cortex dataset is publicly available at https://nilearn.github.io/modules/generated/nilearn.datasets.fetch_haxby.html. The BOLD5000 dataset is publicly available at https://bold5000.github.io/. The Sherlock movie dataset is publicly available at https://dataspace.princeton.edu/handle/88435/dsp01nz8062179.

## Acknowledgments

This work was supported by the McKnight Foundation (JP), NSF CAREER Award IIS-1150186 (JP), the Simons Collaboration on the Global Brain (SCGB AWD1004351) (JP) and a J. Insley Blair Pyne Fund Award (to JP, BE, KN).

## Appendices

## A Relationship between brain kernel, Gaussian process latent variable model (GPLVM) and deep GPs

The brain kernel model can be perceived from two perspectives.

- It is a Gaussian process latent variable model (GPLVM) with another Gaussian process prior over the latent. GPLVM [53, 54] extends the GP regression model by treating the inputs **X** = (**x**_1_,…, **x**_*n*_) as latent variables, distributed independently with standard normal distributions: 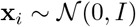, for *i* = 1,…, *n*. Unlike the regression setting, the goal of the GPLVM is to account for structure of an observed dataset in terms of a set of locations in a latent space, which are transformed by a smooth function to generate the observed function values. The primary quantity of interest in the GPLVM is therefore *P*(**X**|**f**), the conditional distribution over the latent variables given the observations. The smoothness induced by the GP prior distribution ensures a smooth mapping from latent variables to observed data. This means that when observed values **f**_*i*_ and **f**_*j*_ are close together, they are likely to arise from from points x_*i*_ and x_*j*_ that are close together in the latent space. Conversely, if observed values **f**_*i*_ and **f**_*j*_ are far apart, smoothness of *f* implies that the corresponding input values lie far apart in the latent space. In a nutshell, the brain kernel model extends the GPLVM to allow latent locations arise from a smooth nonlinear transformation of the 3D voxel locations in the brain.
- The brain kernel model can also be seen as a “deep Gaussian process” model [55] with two layers. It is a sequence of two nonlinear functions, each with a Gaussian process prior. In the main text, we explained that the latent embedding **z** is obtained by applying a function *f* over the input 3D coordinate x, where 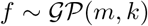. This is the first layer GP. The second component of the brain kernel model is a probability distribution over neural activity as a function of location in embedding space. Our modeling assumption is that neural activity changes smoothly as a function of location in embedding space, or equivalently, that correlations in neural activity fall off smoothly with distance in latent space. This motivates the use of a second GP to model each vector of neural activity. Thus:

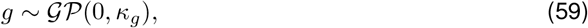

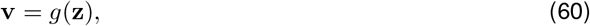

where *k_g_*(**z**_*i*_, **z**_*j*_) = exp (−||**z**_*i*_ − **z**_*j*_||^2^/2) is a standard RBF covariance function (marginal variance and length-scale = 1), and **v** ∈ ℝ^*n*^ denotes a vector of *n* voxels’ neural activity at latent locations {**z**_1_,…, **z**_*n*_}. Given that latent voxel locations are themselves a nonlinear function of the true voxel locations, this allows to write, equivalently: **v** = *g*(*f*(**x**_1_,…, **x**_*n*_)). Therefore, we could eventually obtain the neural activity **v** by applying a two-layer GPs to the input 3D coordinates **X**.

Another interesting connection between the brain kernel model and GPLVM is that the brain kernel model is a nonlinear extension of the dual probabilistic principal component analysis (dual PPCA) which is a special case of GPLVM, according to [54]. If the prior over **z** in eq. 8 is a normal prior and the kernel in eq. 11 is a linear kernel, we could write our brain kernel model as a dual PPCA formulation. Given this equivalence, we can tell that the major difference between the brain kernel model and PPCA is the use of a nonlinear kernel and a GP prior over the latent. The nonlinear kernel allows a nonlinear mapping between the latents and the measurements of neural activity. The GP prior allows us to impose a smoothness assumption over the latent space which regularizes the search of latent values and achieves a nonlinear warping from the voxel space to the latent space. PPCA cannot achieve either.

## B comprehensive summary of application results

In this section, we present a comprehensive summary of decoding and factor modeling results over four fMRI datasets. Table. 1 shows the datasets and the brain analyses we ran. Check marks indicate the brain kernel prior has the best performance; cross marks imply the brain kernel prior doesn’t have the best performance. We demonstrate more analyses by datasets in the following subsections.

**Table 1.** Datasets and brain analyses we ran

| dataset | decoding | factor modeling |
| --- | --- | --- |
| HCP | ✓ | ✓ |
| Sherlock movie | ✗ | ✓ |
| Visual recognition | ✓ | ✗ |
| BOLD5000 | ✗ | ✗ |

## B.1 HCP database

**Fig 15.**
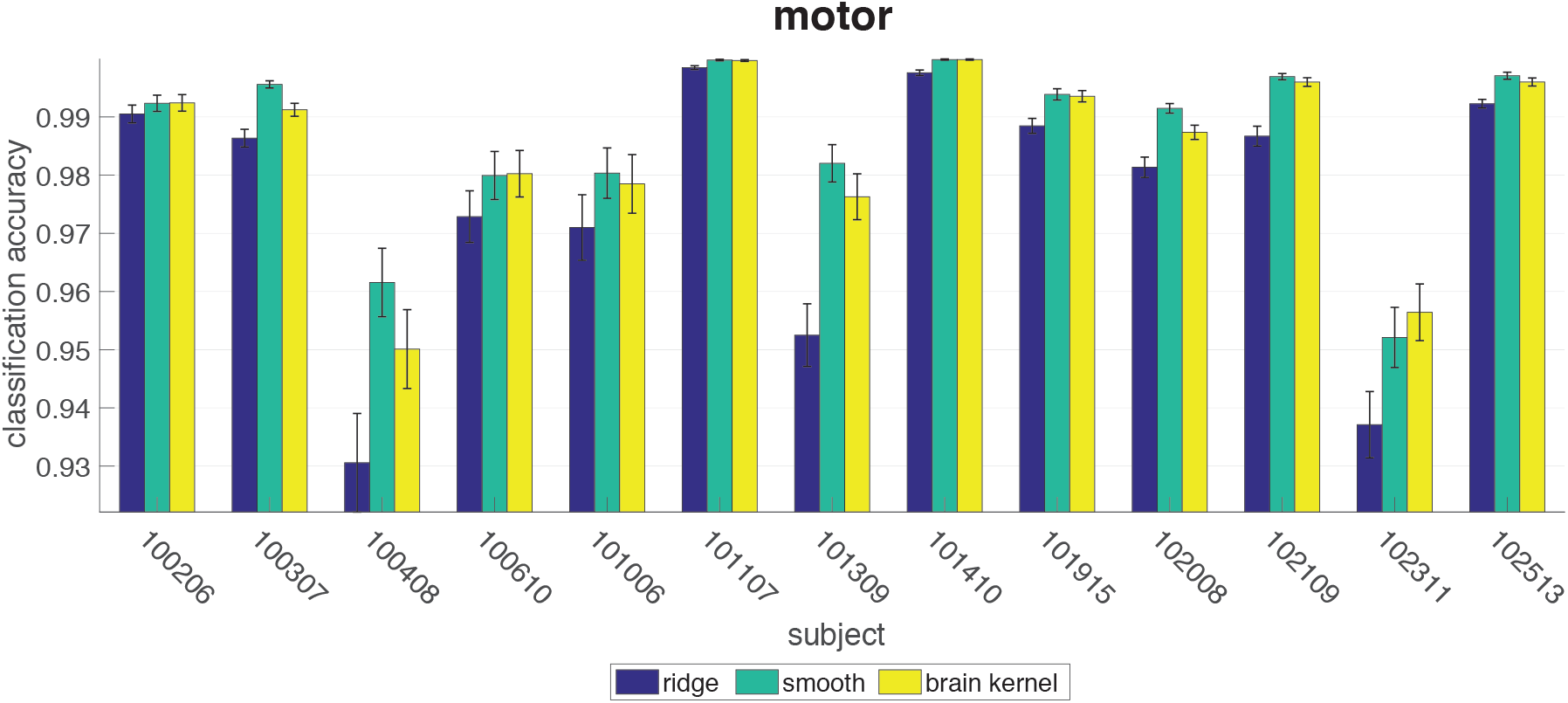
Accuracy performance on the motor task.

**Fig 16.**
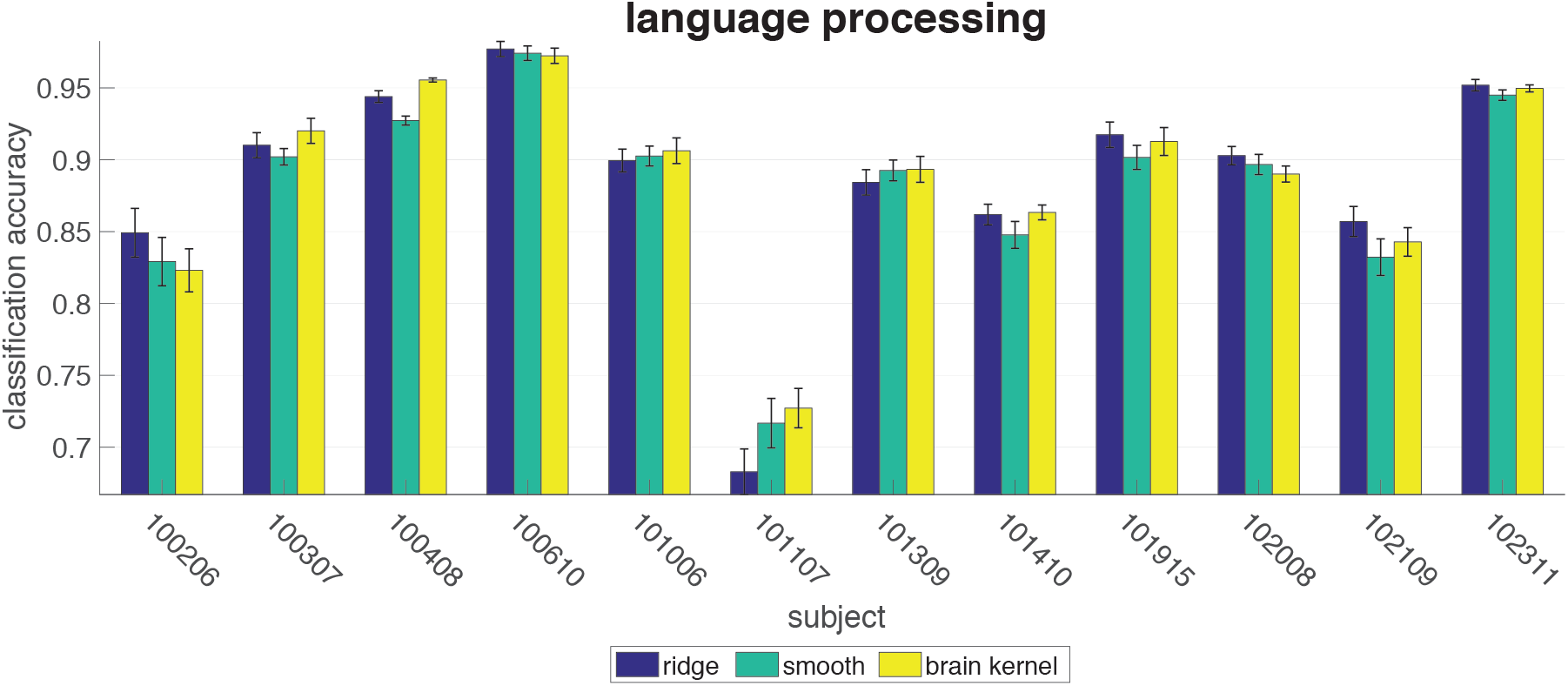
Accuracy performance on the language processing task.

**Fig 17.**
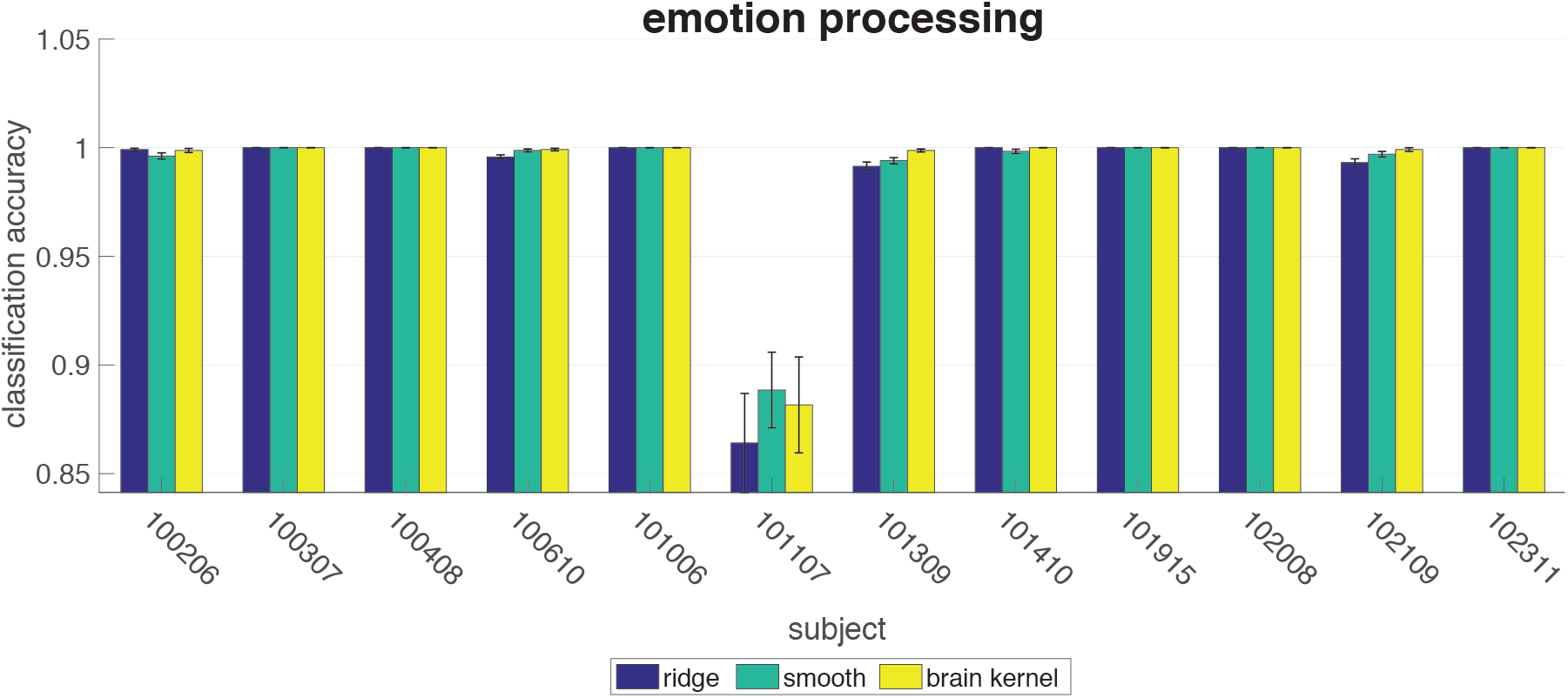
Accuracy performance on the emotion processing task.

**Fig 18.**
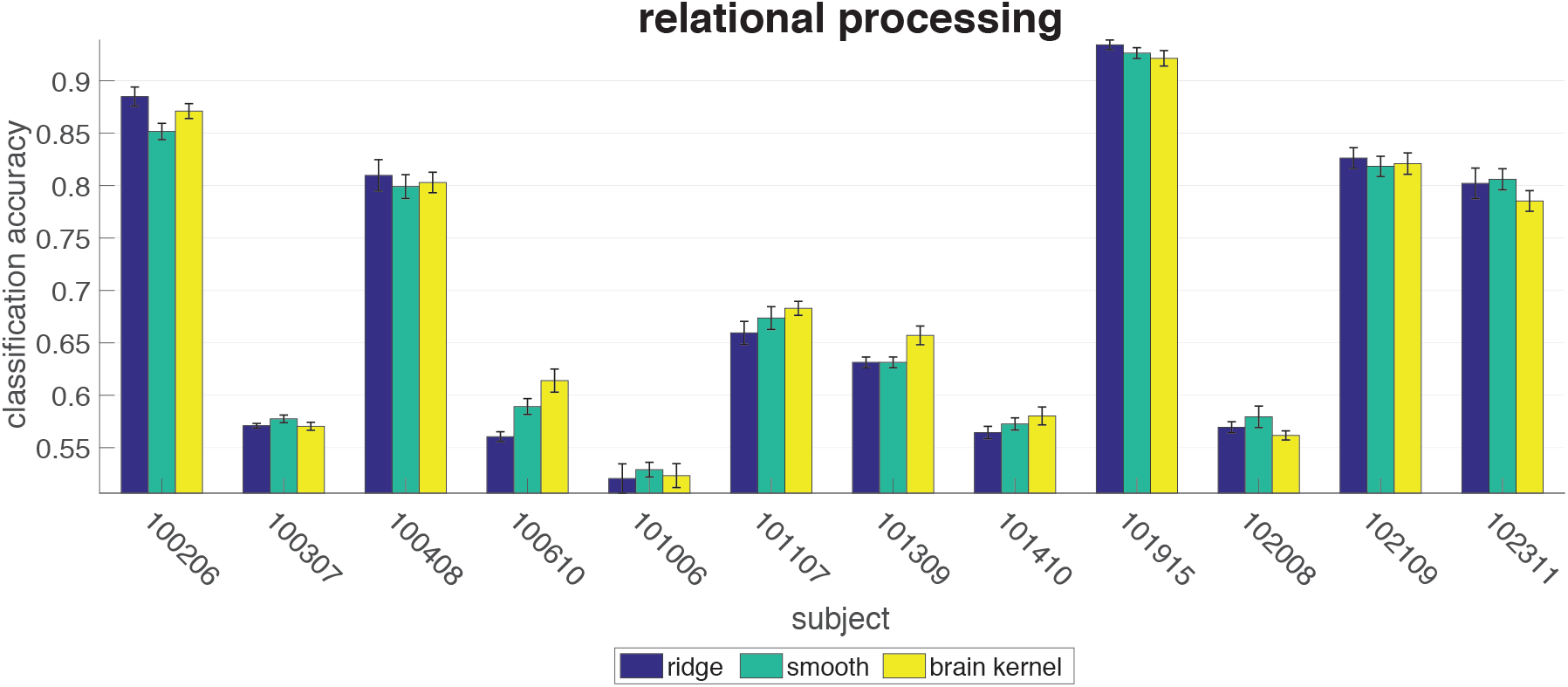
Accuracy performance on the relational processing task.

**Fig 19.**
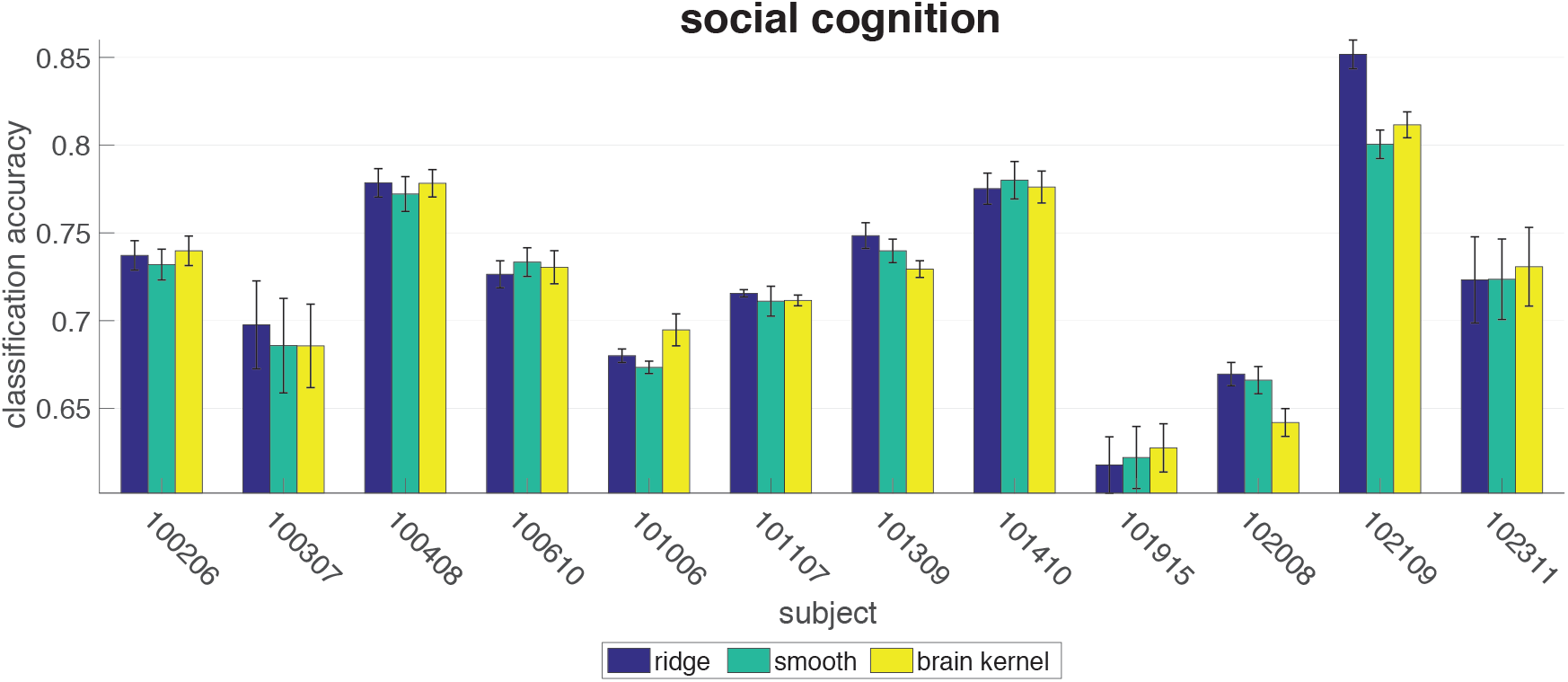
Accuracy performance on the social cognition task.

**Fig 20.**
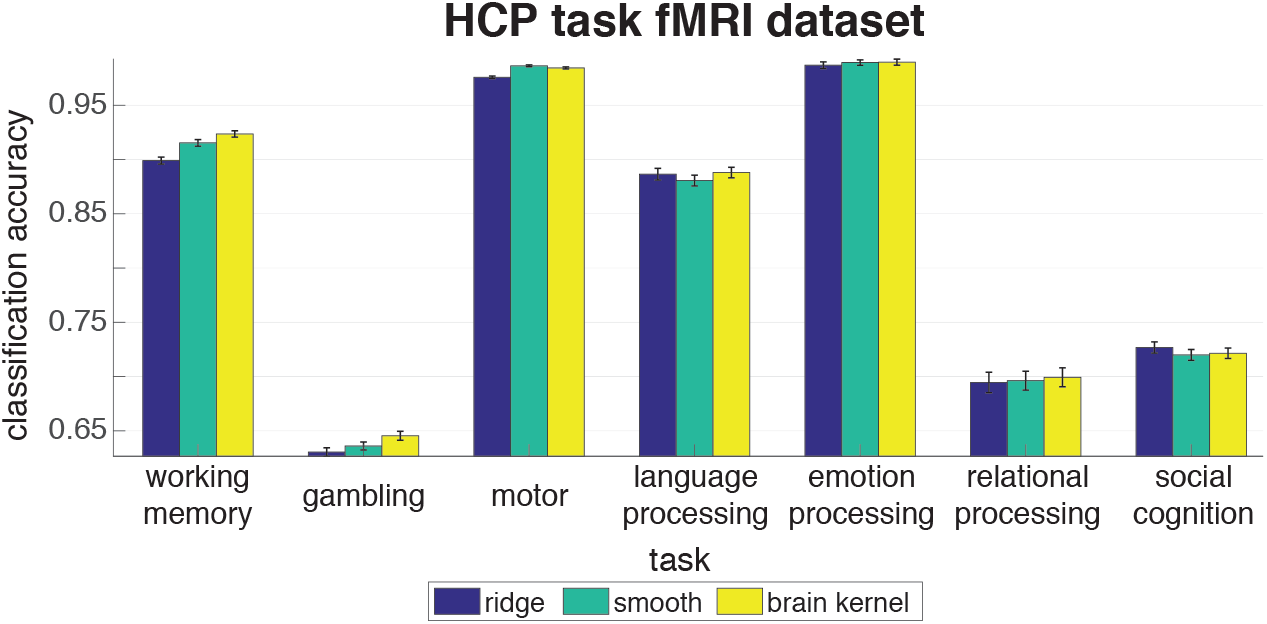
Accuracy performance for all task fMRI datasets in the HCP database averaged over subjects. The x-axis indicates the task. The y-axis is accuracy performance.

**Table 2.** The basic information of the fMRI scans [24]

| Task | Runs | Frames per run | Run Duration (min:sec) |
| --- | --- | --- | --- |
| Working Memory | 2 | 405 | 5:01 |
| Gambling | 2 | 253 | 3:12 |
| Motor | 2 | 284 | 3:34 |
| Language Processing | 2 | 316 | 3:57 |
| Social Cognition | 2 | 274 | 3:27 |
| Relational Processing | 2 | 232 | 2:56 |
| Emotion Processing | 2 | 176 | 2:16 |

## Decoding

We’ve elaborated on the working memory and gambling tasks in the main paper. In this section, we present the classification accuracy performance for the motor task, the language processing task, the emotion processing task, the relational processing task, and the social cognition task. Similar to the working memory and gambling tasks, each task fMRI was from the same HCP project as the resting-state fMRI. They shared the same coordinate system and preprocessing pipeline. The task fMRI was aligned with the MNI template with the same 59,412 voxels as the resting-state fMRI data. Instead of analyzing the whole-brain data for brain decoding, we worked with functional ROIs whose averaged activity levels were above a certain threshold for all tasks. We formulated each task as a binary classification problem. We used the same Bayesian linear regression classifiers as described in the working memory and gambling sections. We trained the classifier with three priors on one run and calculated the accuracy performance on the second run, then switched the training and test runs. This procedure was repeated ten times. The ±1 labels were treated as continuous target values during training, and the test accuracy was evaluated by taking the sign of the prediction. We computed the averaged accuracies across two runs and 10 repetitions for the three priors for all subjects in each dataset. Results are presented in Fig. 15 to Fig. 19. Other than the working memory and gambling tasks, the brain kernel performed mildly better than the ridge prior and the smoothing prior estimates. Here we also summarize the averaged accuracy over all subjects for each task in Fig. 20. The overall performance of these HCP task fMRI datasets indicates that the brain kernel is still a better choice than the smoothing and the ridge priors.

## Factor modeling

We examined the task fMRI datasets in the HCP database with the Bayesian FA model. We collected all the task fMRIs for the same 10 subjects and performed Bayesian FA for each subject in each task. We implemented the “co-smoothing” evaluation and compared the test *R*^2^ for the three priors averaged over all subjects for each task (the normalized test *R*^2^ in Fig. 21 and the standard test *R*^2^ in Fig. 22). In most tasks, the brain kernel outperformed the ridge prior and the smooth RBF kernel. This implies that the brain kernel provided a superior prior covariance for the latent source matrix and may enhance performance in terms of data explanation for the HCP database.

**Fig 21.**
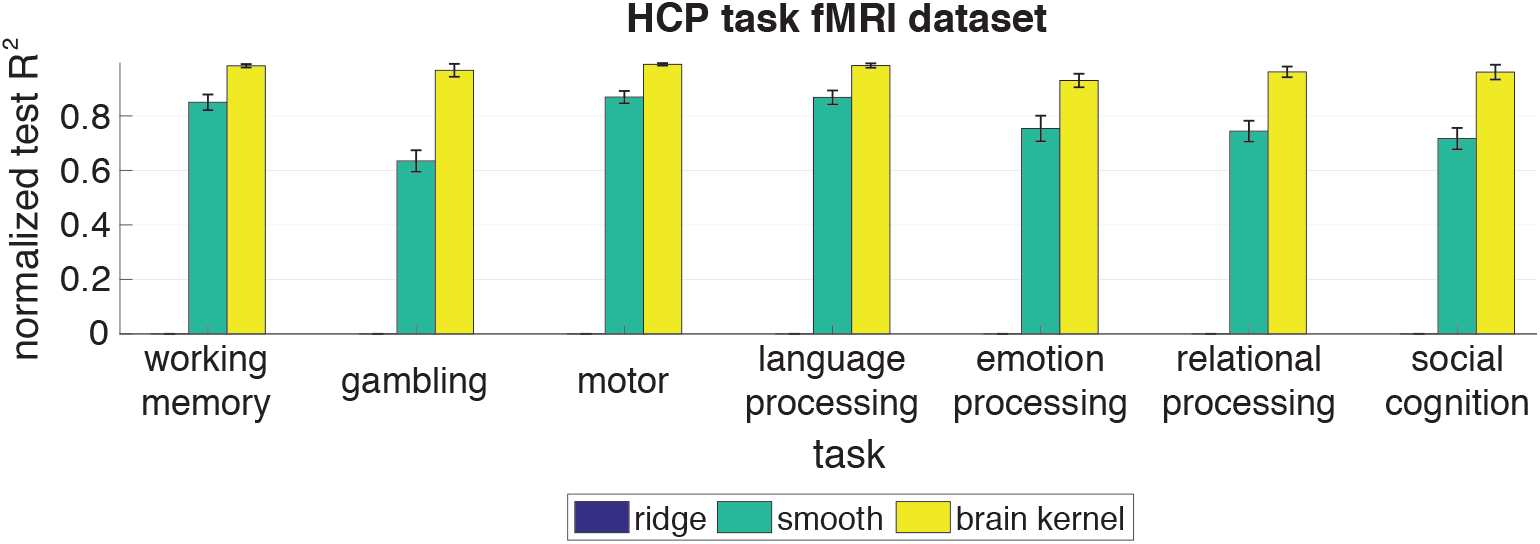
Normalized test *R*^2^ performance on the HCP task fMRI datasets. The x-axis indicates the task. The y-axis is the normalized test *R*^2^ performance averaged over 10 subjects for each task (higher values indicate better performance). The error bars indicate standard errors. We compared our brain kernel with a ridge prior and a smooth RBF kernel, color coded.

**Fig 22.**
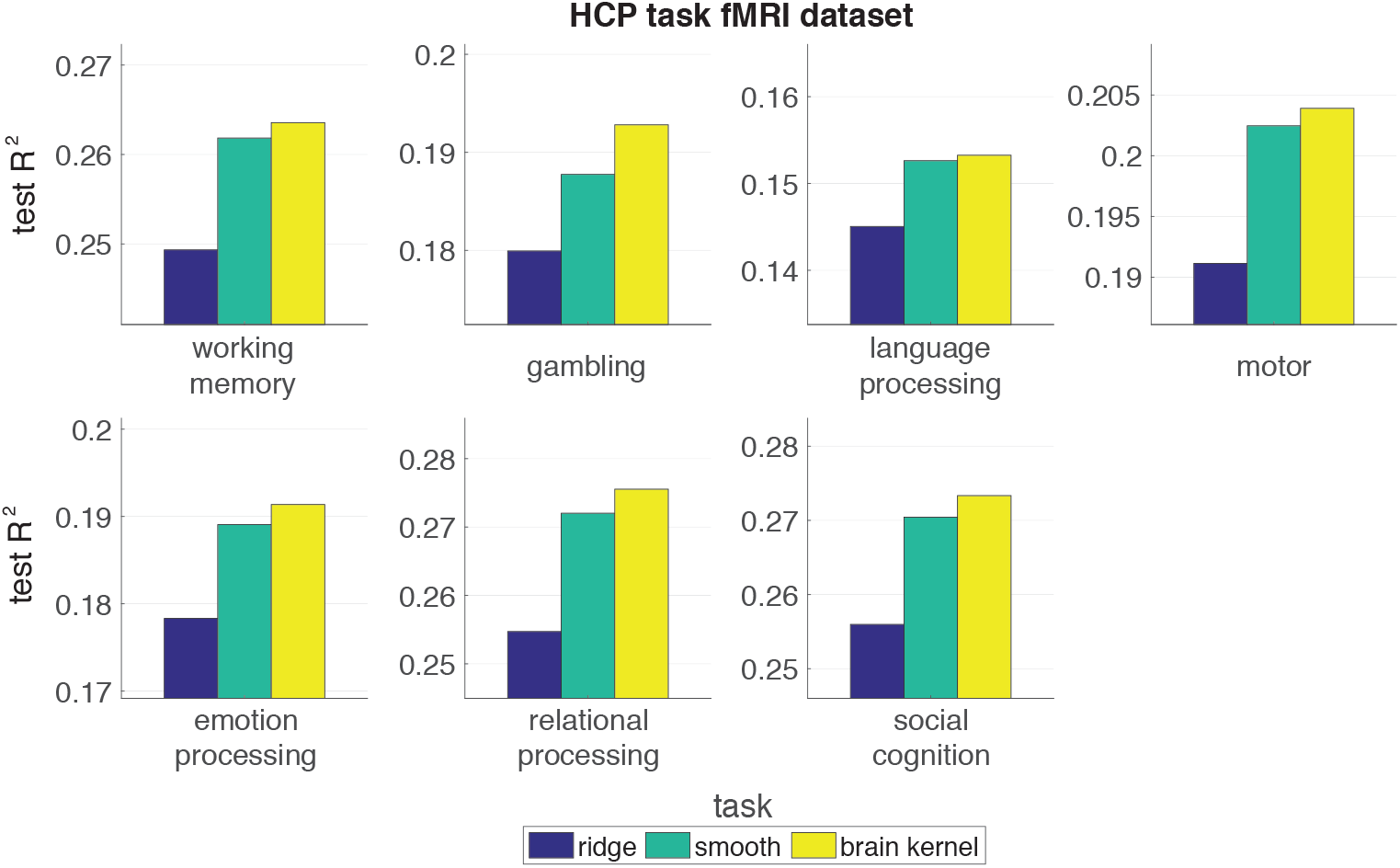
Test *R*^2^ performance on the HCP task fMRI datasets.

We did further inspection into the covariance matrices shown in Fig. 23 for the working memory task. The top-left sub-figure is the sample covariance of the selected working memory ROIs indicated in red in the bottom-left sub-figure. The right two columns show the optimal RBF and brain kernel during decoding (top row) and factor modeling (bottom row). To understand the estimated covariance, we first recall that the smoothing prior and the brain kernel prior are constructed as follows:

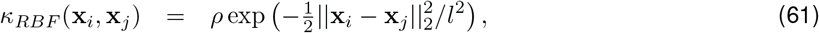

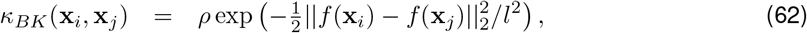

where *ρ* is the marginal variance and *l* is the length-scale. We know *f* for the brain kernel prior. We selected the optimal hyperparameters via a 5-fold cross validation. The optimal hyperparameters for decoding were selected that maximized the classification accuracy. The optimal hyperparameters for factor modeling were obtained when matching eq. 49 to the sample covariance. The matching was evaluated by calculating the negative log-likelihood (NLL) of the data coming from a Gaussian distribution with a zero mean and a covariance (eq. 61 or 62). The smaller the NLL is, the better. The optimal prior covariance matrices were evaluated with the optimal *ρ* and *l* for the smoothing prior and the brain kernel prior respectively. For decoding, the best smoothing RBF kernel only captured local smoothness while the brain kernel captured some off-diagonal covariance across hemispheres. The corresponding optimal weights are also visualized in Fig. 7B. For factor modeling, since our goal was to find the latent sources that best captured the statistical structures in the data, the optimal prior covariance should resemble the sample covariance. Given this intuition, we could visually identify the brain kernel as a better approximation than the RBF kernel. Moreover, the NLL value for the brain kernel is also smaller than the RBF kernel, indicating that the brain kernel approximated the sample covariance quantitatively better. The interpretation of these covariance matrices provide us some intuition of why the brain kernel prior worked with the HCP database.

**Fig 23.**
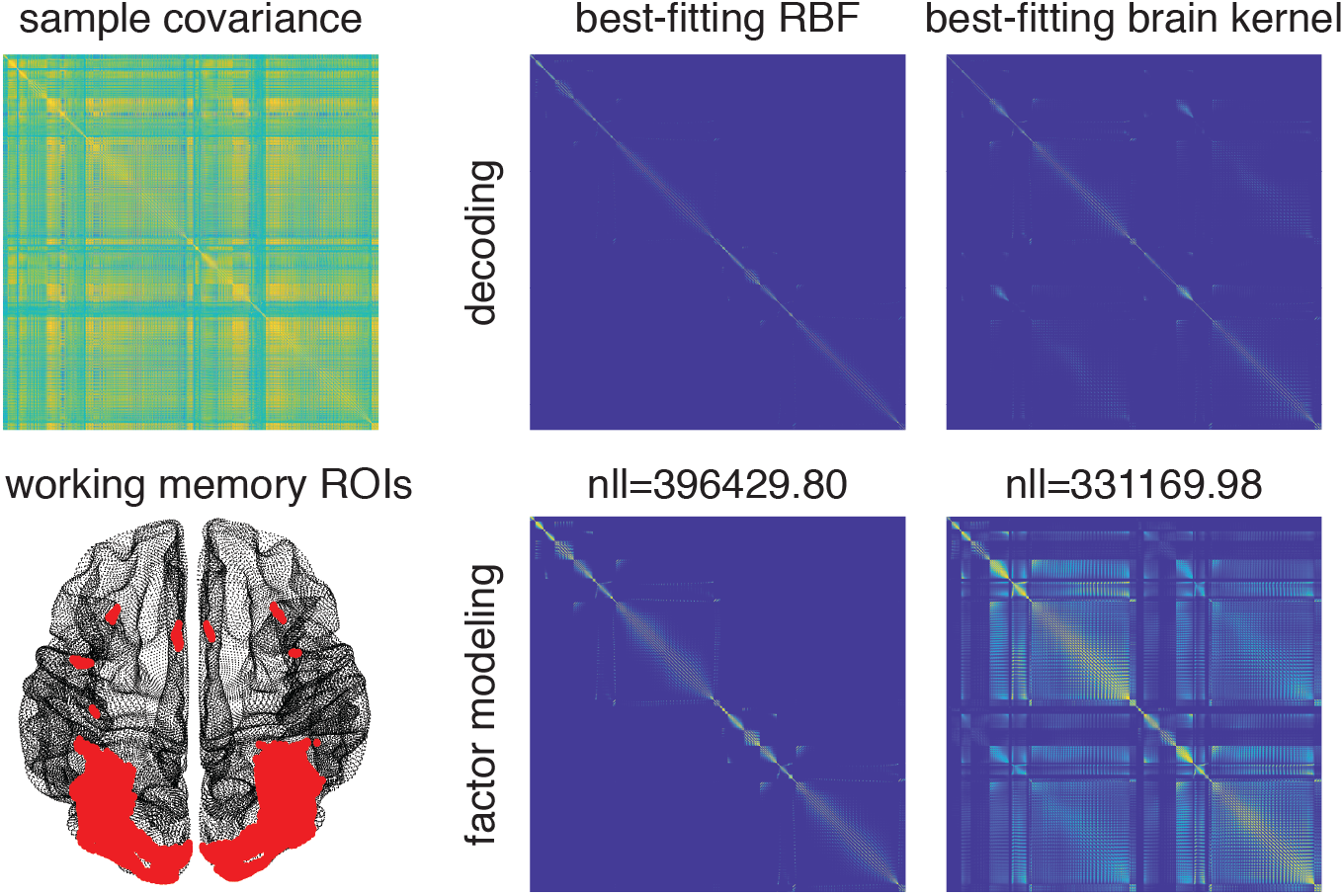
Best-fitting RBF and brain kernel for the working memory task.

Moreover, for factor modeling, we carried out the same analysis as we did in Fig. 23 to all other tasks in HCP as summarized in Table 3. The first column corresponds to the ROIs we used from each task. The second column indicates which dataset we used: resting-state or task. Although this is a factor modeling task, the essence of this analysis is to find the best kernel that fits the sample covariance by tuning the hyperparameters in eq. 61 and eq. 62. We could do this analysis for both resting-state and task fMRIs. The working memory ROI with resting-state means that we used the resting-state data with the working memory ROI. The third and fourth columns contain the optimal length-scale and optimal NLL when fitting the smoothing RBF kernel. The last two columns contain the optimal length-scale and optimal NLL when fitting the brain kernel. The two NLL values shown in Fig. 23 locate in the fourth and sixth cells on the fourth row. The creation of this table is mainly to understand how well the brain kernel captures spatial covariance patterns observed during task performance.

**Table 3.** Fitting the RBF kernel and the brain kernel to both the resting-state fMRI and the task fMRIs in HCP.

| task ROI | state | Smoothing RBF |  | Brain Kernel |  |
| --- | --- | --- | --- | --- | --- |
|  |  | length-scale | NLL | length-scale | NLL |
| working memory | resting | 12.51 | 10321.5 | 1.00 | 10302 |
|  | task | 3.95 | 396429.80 | 0.52 | 331169.98 |
| gambling | resting | 10.68 | 14204.5 | 1.00 | 14177.9 |
|  | task | 3.45 | 88200.56 | 0.52 | 116572.23 |
| motor | resting | 5.68 | 10765.9 | 1.00 | 10756.2 |
|  | task | 3.01 | 100834.85 | 0.52 | 86879.27 |
| language process | resting | 6.65 | 12687.4 | 1.00 | 12672.6 |
|  | task | 2.63 | 198685.16 | 0.39 | 120010.67 |
| emotion process | resting | 12.51 | 12345.4 | 1.00 | 12321.6 |
|  | task | 2.29 | 129664.89 | 0.17 | 32696.75 |
| relational process | resting | 10.68 | 11268.8 | 1.00 | 11246.5 |
|  | task | 3.95 | 215712.36 | 0.59 | 269039.82 |
| social cognition | resting | 7.79 | 13043.4 | 1.00 | 13019.3 |
|  | task | 2.63 | 162456.39 | 0.68 | 328688.55 |

## Comparison between resting and task

We summarize the best-fitting resting-data and task-specific length-scale and the optimal negative log-likelihood (NLL) in Table 3. Different tasks have different length-scales when fitting with both the smoothing kernel and the brain kernel. For the brain kernel, the optimal length-scales are 1 for all task ROIs using resting-state data, while the optimal length-scales are smaller than 1 for all task data, indicating the task covariance fell off faster than the resting state data in the nonlinear embedding space. This is also observed in the Euclidean space employed by the smoothing kernel.

## Comparison between smoothing and brain kernel

In addition, it is difficult to make claims about the absolute performance of the brain kernel model on the task fMRI datasets. However, we can show the relative performance between the brain kernel and the RBF smoothing kernel. In Table 3, some tasks (including working memory, motor, language process, and emotion process) have smaller NLLs with the brain kernel than with the RBF kernel. This implies that for these tasks, the brain kernel latent we estimated using the resting-state data captured more task covariance than a Euclidean-distance based RBF kernel. For the other tasks (including gambling, relational process, and social cognition), the brain kernel has larger NLL values than the RBF kernel. The covariance patterns for all these tasks are similar to Fig. 23 where the brain kernel covariance looks much more alike to the sample covariance with more off-diagonal structures while the RBF kernel is sparse.

When resting-state and task fMRI share the same voxel space, the brain kernel doesn’t have the voxel alignment issue. The good performance indicates the resting-state functional connectivity is able to provide the kernel most neutral with respect to task-induced states and capture the majority of task covariance. In many fMRI experiments, researchers collect task fMRI as well as massive resting-state fMRI while participants are resting. In this case, both resting-state and task fMRIs are collected with the same experimental setup and processed with the same pipeline. Then it’s likely to achieve improved brain analyses on the task data using the brain kernel prior which is estimated from the resting-state fMRI. With these analyses, we hope to convince the readers that the brain kernel is a reasonable and better prior option than the smoothing RBF.

## B.2 Sherlock movie fMRI dataset

We examined the Sherlock fMRI dataset, in which participants were scanned while they watched the British television program “Sherlock” for 50 min [34]. The fMRI data comprised 1,973 TRs (Repetition Time), where each TR was 1.5 s of the movie. Before performing any analysis, the fMRI data were preprocessed and aligned to MNI space using the techniques described in prior work [34]. We examined the brain data averaged across all subjects to smooth out individual variability. We identified 11 ROIs previously implicated in processing naturalistic stimuli, comprising the default mode network (DMN-A, DMN-B), the ventral and dorsal language areas, and the primary auditory and visual cortices [35].

## Decoding

For each ROI, we performed a standard decoding task to relate the fMRI signal to the representation of the semantic content of each movie frame [56]. We trained the decoding model using the first half of the movie and tested on the second half of the movie. The decoding task was called “scene classification”. We divided up the second half of the movie into 25 uniformly-sized chunks, and used the correlation to match predicted semantic content with the ground truth for each chunk, and reported the percentage of the chunks that the match was perfect. Since there are 25 chunks, random chance performance at this task is 4%. Fig. 24 presents the classification accuracy performance. We observe that the smoothing prior was better than the brain kernel prior, both better than the ridge prior. Then we investigated the reason of the performance difference. Fig. 25 presents the sample covariance for the DLN ROI and its corresponding optimal covariance matrices. Observing the top row, we can tell that the smoothing covariance exhibited much off-diagonal covariance across voxels, while the brain kernel only presented a small amount. If we increase the length-scale l for the brain kernel, we will see more off-diagonal covariance emerge; accordingly, extra cross-covariance will also emerge that imposes harmful correlations against the decoding task. Therefore, the conclusion is the brain kernel is able to impose a strong cross-covariance assumption over many voxels; but much of the cross-covaraince doesn’t exist in the Sherlock movie data or could hurt the decoding performance. The cross-validation procedure conservatively selected a small length-scale that led to less smoothing assumptions compared with the smoothing kernel. So this Sherlock movie dataset is not an ideal dataset that could leverage most of the resting-state functional connectivity to do the decoding task.

**Fig 24.**
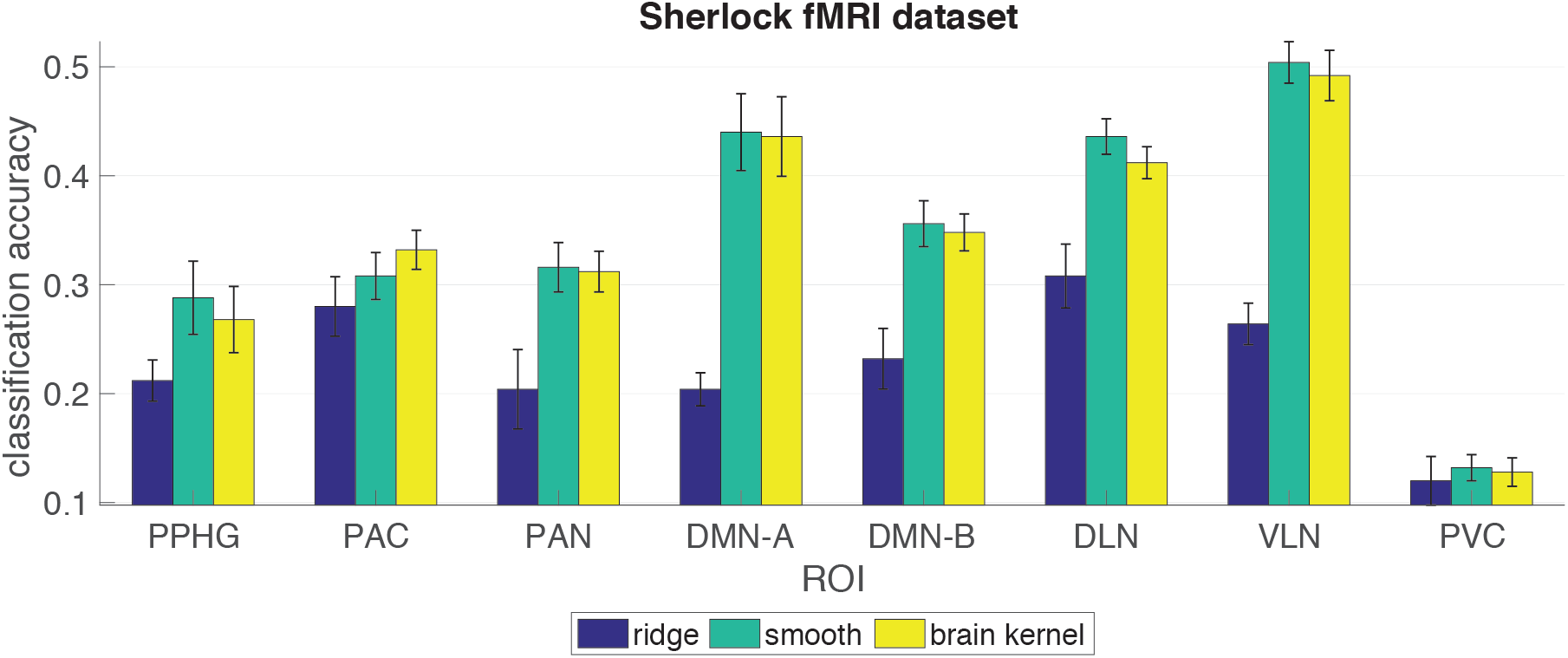
Accuracy performance on the Sherlock movie fMRI dataset. The ROI information is the same as Fig. 13

**Fig 25.**
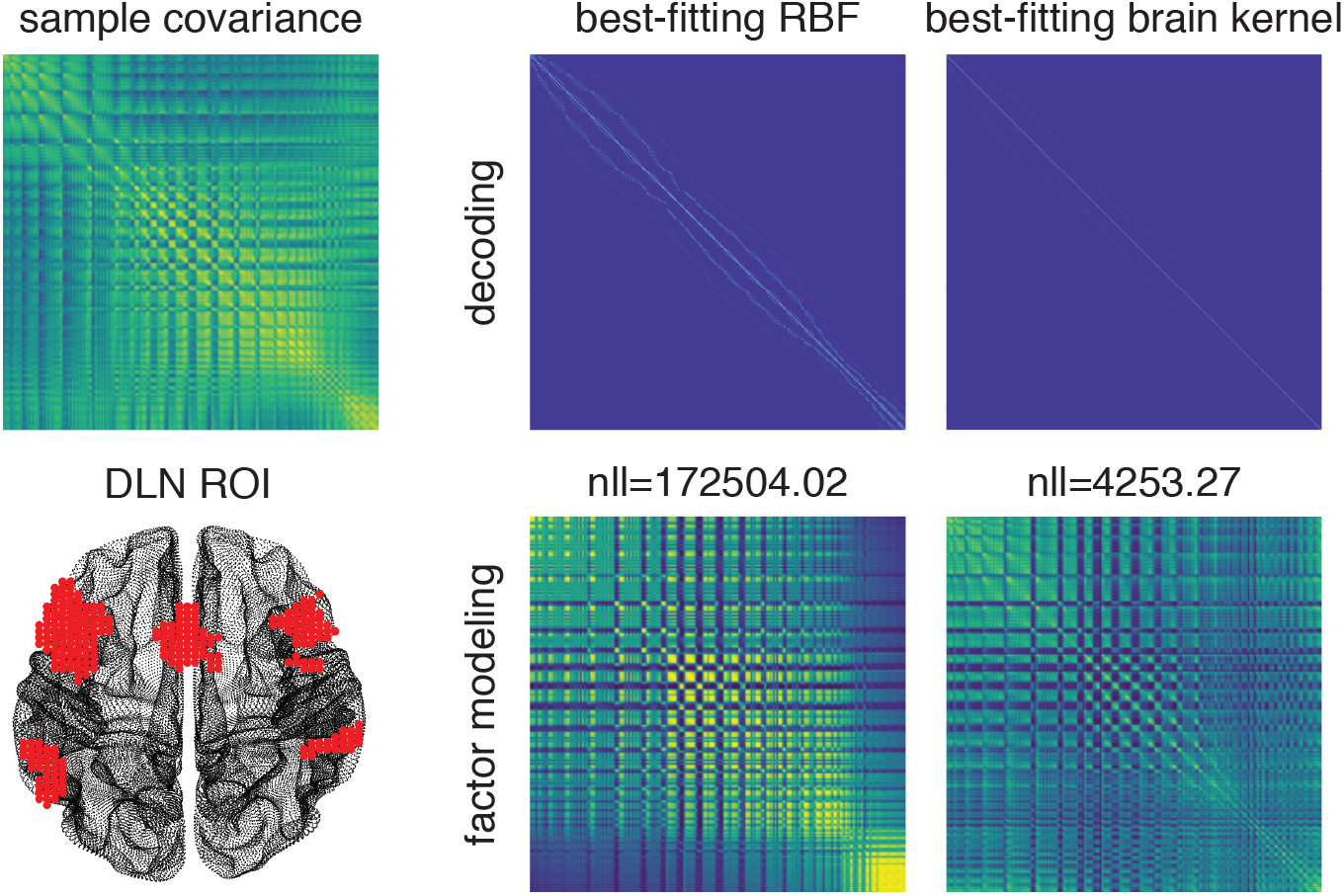
Best-fitting RBF and brain kernel for the DLN ROI.

## Factor modeling

For each ROI, we also performed a standard factor analysis (FA) to factorize the voxel-by-time fMRI data into a latent source matrix and a factor matrix. We achieved three *R*^2^ values for the three priors (the normalized test *R*^2^ in Fig. 26 and the standard test *R*^2^ in Fig. 27). In most regions, the brain kernel outperformed the ridge prior and the smooth RBF kernel. By observing Fig. 25 bottom row, we can claim that visually the best-fitting BK resembled the sample covariance more than the best-fitting RBF. The NLL value of the brain kernel is also smaller, implying that the brain kernel served as a better approximation to the sample covariance. This provides a nice intuition of why the BK prior performed well for factor modeling with the Sherlock movie dataset. This result implies that when performing Bayesian FA, the brain kernel provided a superior prior covariance for the latent source matrix and may enhance performance in terms of data explanation.

**Fig 26.**
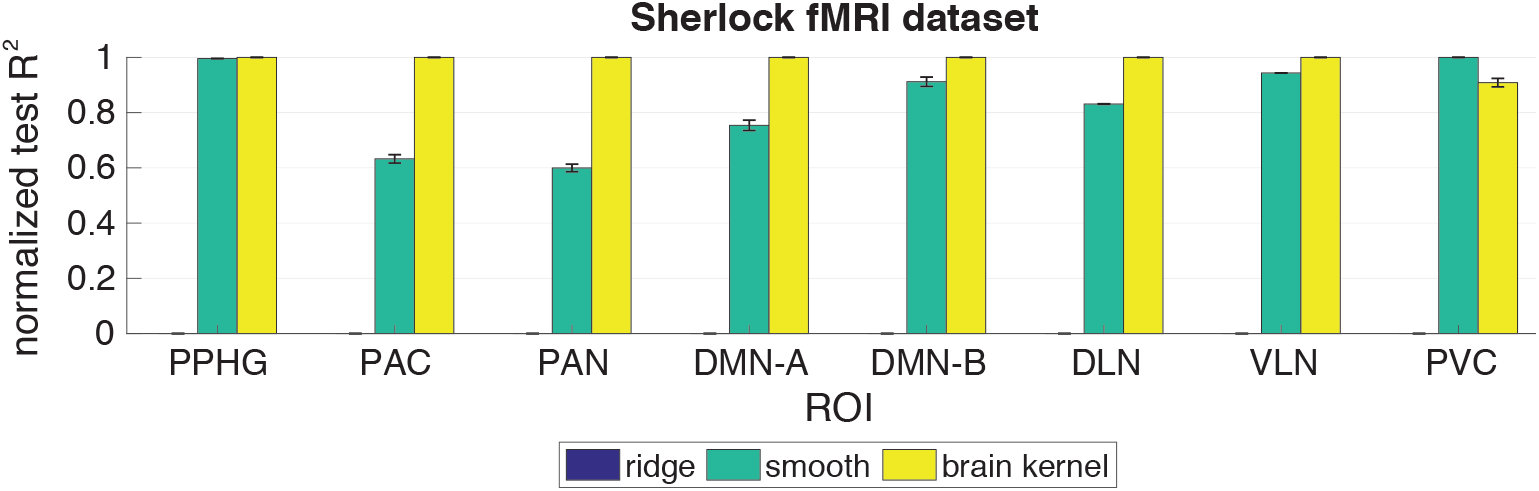
Normalized test *R*^2^ performance on the Sherlock fMRI dataset. The x-axis indicates ROIs (PPHG – Posterior Parahippocampal Gyrus; PAC – Primary Auditory Cortex; PAN – Primary Auditory Network; DMN-A – Default Mode Network-A; DMN-B – Default Mode Network-B; DLN – Dorsal Language Network; VLN – Ventral Language Network; PVC – Primary Visual Cortex). The y-axis is the normalized *R*^2^ performance on the test set (higher values indicate better performance). The error bars indicate standard errors. We compared our brain kernel with a ridge prior and a smooth RBF kernel, color coded.

**Fig 27.**
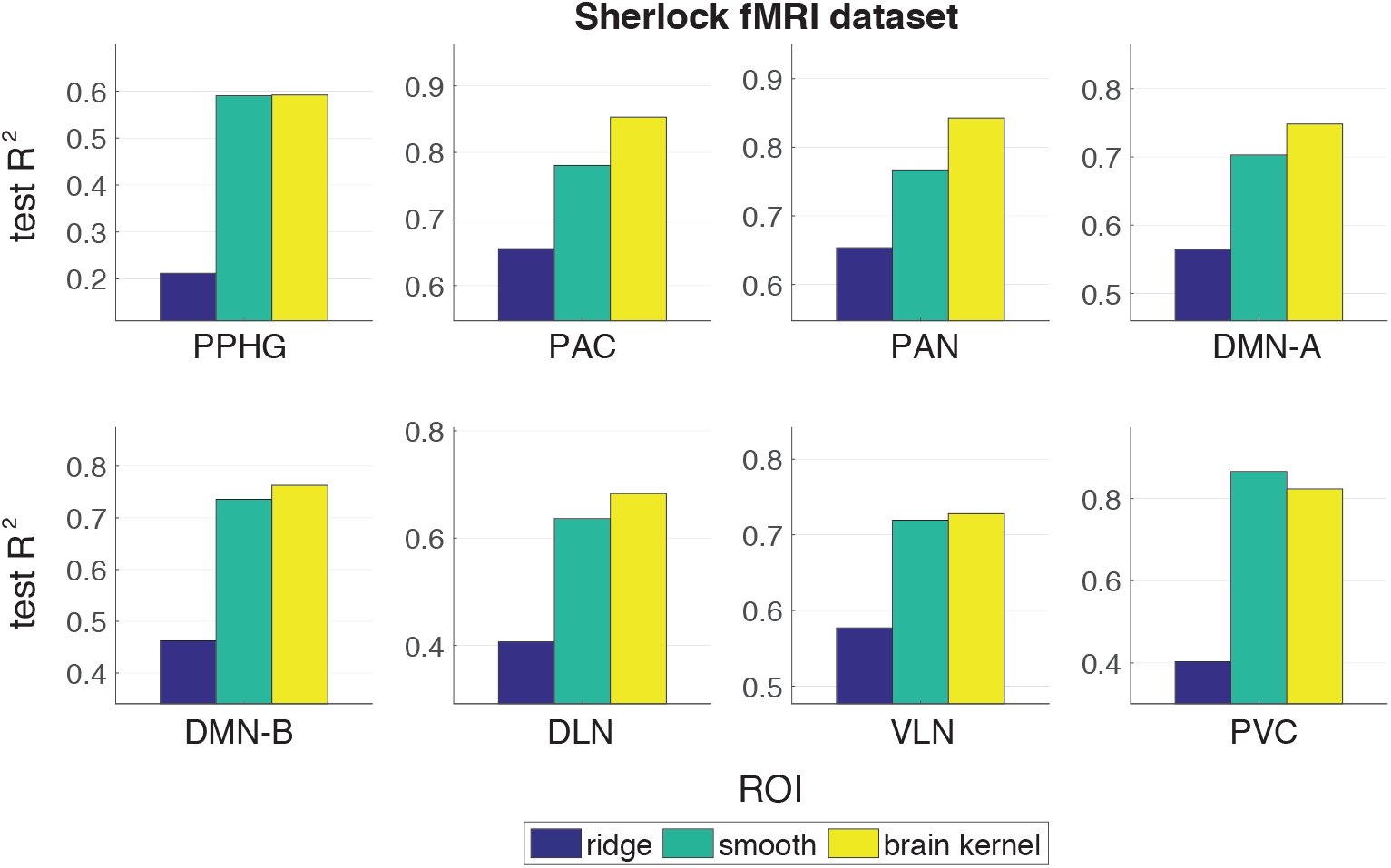
Test *R*^2^ performance on the Sherlock fMRI dataset.

## B.3 Visual recognition task

In the visual recognition experiment, six subjects were asked to recognize eight different types of objects. Each subject participated 12 scanning runs. In each run, the subjects viewed images of eight object categories, with 11 whole-brain measurements per category. Each subject’s fMRI data was preprocessed using the fMRIprep package^3^ [32] and aligned to the MNI template. We extracted ROIs with 1645 voxels in the ventral temporal cortex, which is thought to be involved in object recognition. The ROI mask was obtained from Nilearn [33].

## Decoding

We assessed performance by training Bayesian linear regression classifiers to discriminate between pairs of objects, e.g., face vs. bottle, for each of the 28 possible binary classifications among the eight objects. We trained the weights **w** for each model using linear regression from fMRI measurements x to binary labels *y* ∈ {−1, +1}, and assessed accuracy on the test set using predicted labels 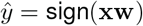. Here, we split the training and test sets by subjects. In this visual recognition dataset, we had six subjects but only 11 × 2 × 12 = 264 measurements in a 1645-voxel space for each subject in a binary decoding task. The number of training measurements was not sufficiently large to train a good classifier that led to a reasonable generalization performance. Therefore, we chose to do inter-subject analyses by using 5 subjects for training and one subject for test. We repeated this leave-one-subject-out manner for six times with each subject being used as the test set once and obtained the result in Fig. 28. The averaged accuracy performance across four repeated runs for six subjects shows that the brain kernel performed comparably to the ridge and smoothing priors with better accuracy performance for 5 out of 6 subjects. This indicates that the brain kernel can provide functional and structural supports for most subjects and tasks. Again, we present the optimal prior covariance for the smoothing prior and the brain kernel prior respectively in Fig. 29 top row. The brain kernel covariance exhibited some off-diagonal values implying that the cross-covariance over the two hemispheres enhanced the decoding performance. The improvement is statistically modest overall based on the standard errors, which could be a result of several factors: misalignment of the coordinate space to the HCP coordinate space used to estimate the brain kernel; mismatch between the resting-state covariance used to construct the brain kernel and covariance present during the visual recognition task; or the object recognition tasks may rely on fine-grained spatial response topographies that are poorly aligned across individuals.

**Fig 28.**
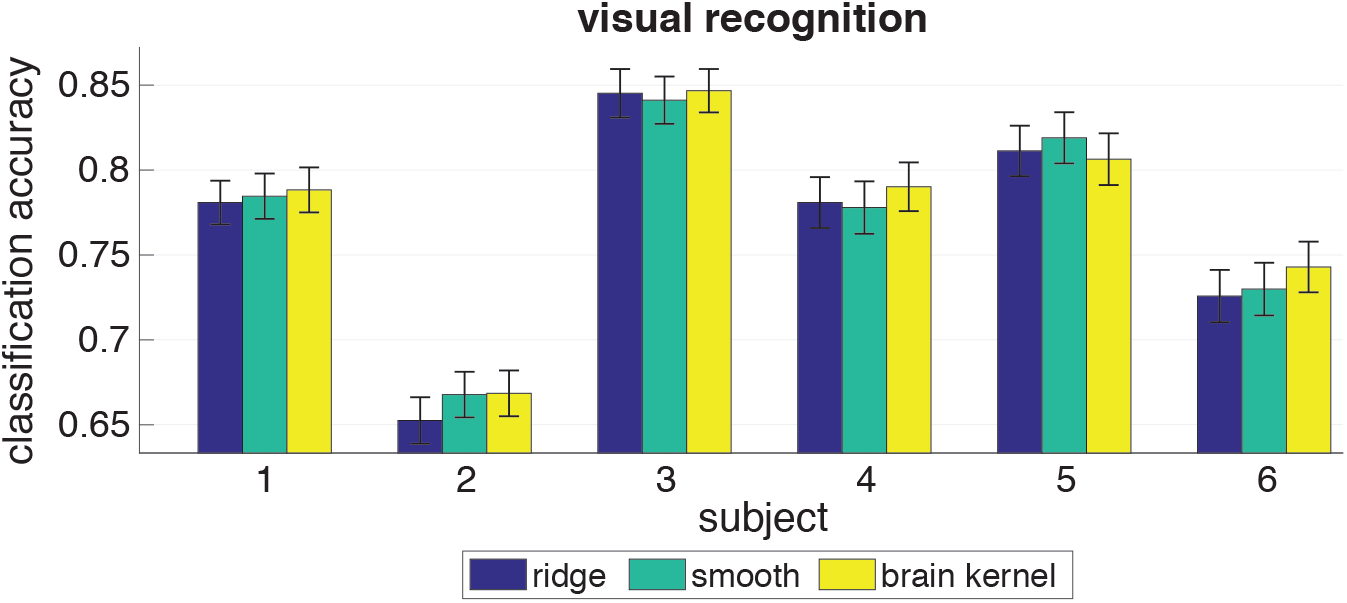
Accuracy performance on the visual recognition task. The x-axis indicates subject IDs. The y-axis is accuracy performance. We compared our brain kernel with a ridge prior and a smooth RBF kernel, color coded.

**Fig 29.**
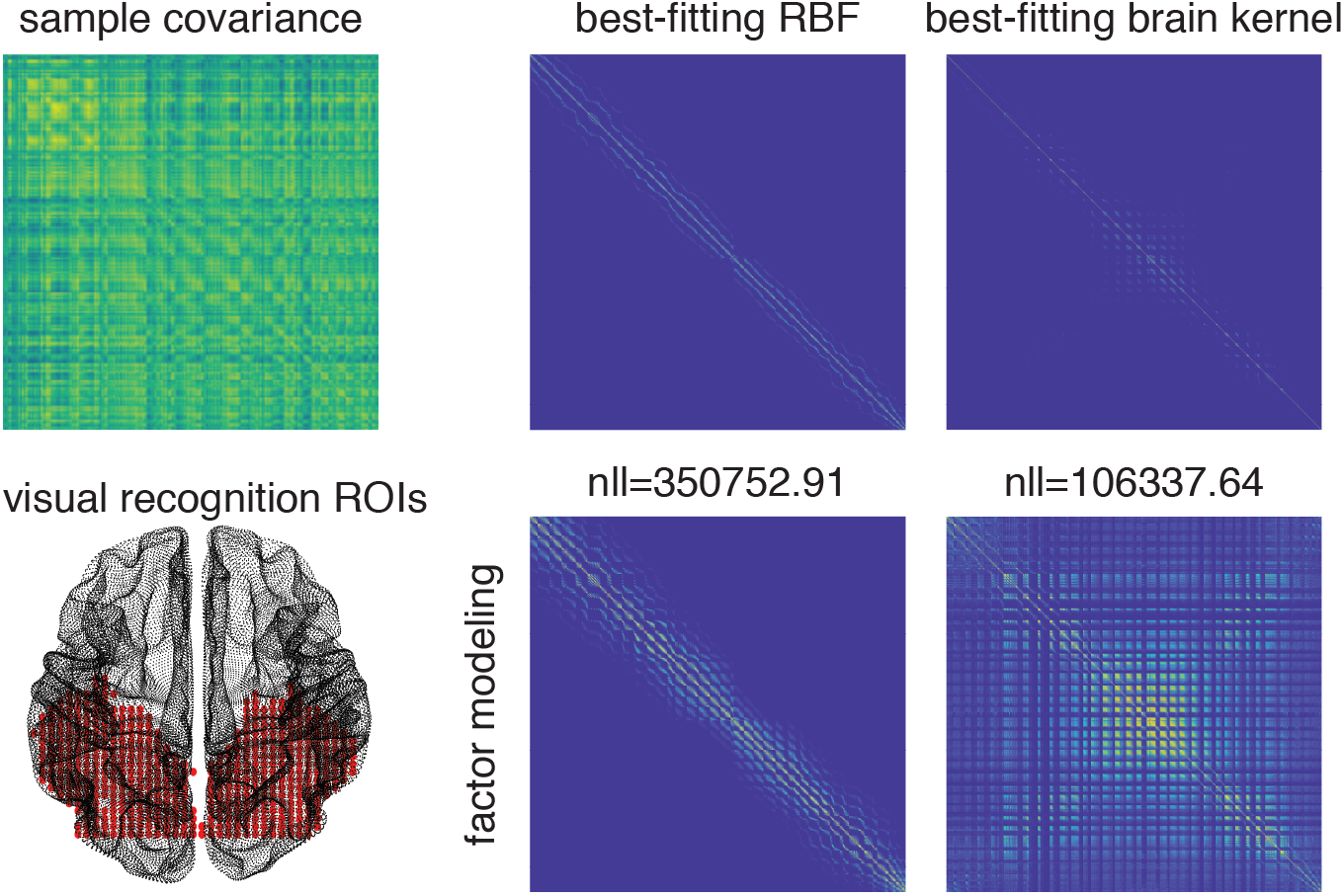
Best-fitting RBF and brain kernel for the visual recognition task.

## Factor modeling

We examined the visual recognition task fMRI dataset with the same Bayesian FA model. We implemented the same “co-smoothing” evaluation as described in the main paper. The only difference is, instead of splitting the runs into halves for training and test within each subject, we treated one subject as the test set and the other five subjects as the training set. This is consistent with the inter-subject classification set-up in the decoding task. We report the normalized test *R*^2^ (Fig. 30) and the standard test *R*^2^ (Fig. 31) for the three priors when treating each subject as the test set. The brain kernel didn’t show a dominating performance over the smooth RBF kernel for most subjects. One thing we need to emphasize here is that both decoding and factor modeling were performed in an inter-subject manner. This made the training data much noisier. The noisiness is not detrimental for decoding since we aimed at extracting a small number of ROI voxels to decode task variables. While for factor modeling, the goal was to extract latent sources that are shared among subjects. These latent sources consist of a large amount of voxels, in which case the noisiness and miss-alignment issues would be magnified. Thus if we look at the *R*^2^ values in Fig. 31, all are negative implying that none of these priors led to reasonable latent sources and signal reconstruction. Moreover, we can visualize the sample covariance and the best-fitting RBF and brain kernel in Fig. 29 bottom row. The RBF kernel was more conservative focusing on the diagonal, while the brain kernel presented a lot of off-diagonal covariance. We could qualitatively evaluate that the brain kernel didn’t resemble the sample covariance very much despite its smaller NLL value. Thereby the off-diagonal cross-covariance could potentially harm the performance. This provides an intuition of why the BK prior performed worse for factor modeling with the visual recognition dataset. But notably, both the smoothing prior and the brain kernel prior improved the *R*^2^ values upon the ridge prior.

**Fig 30.**
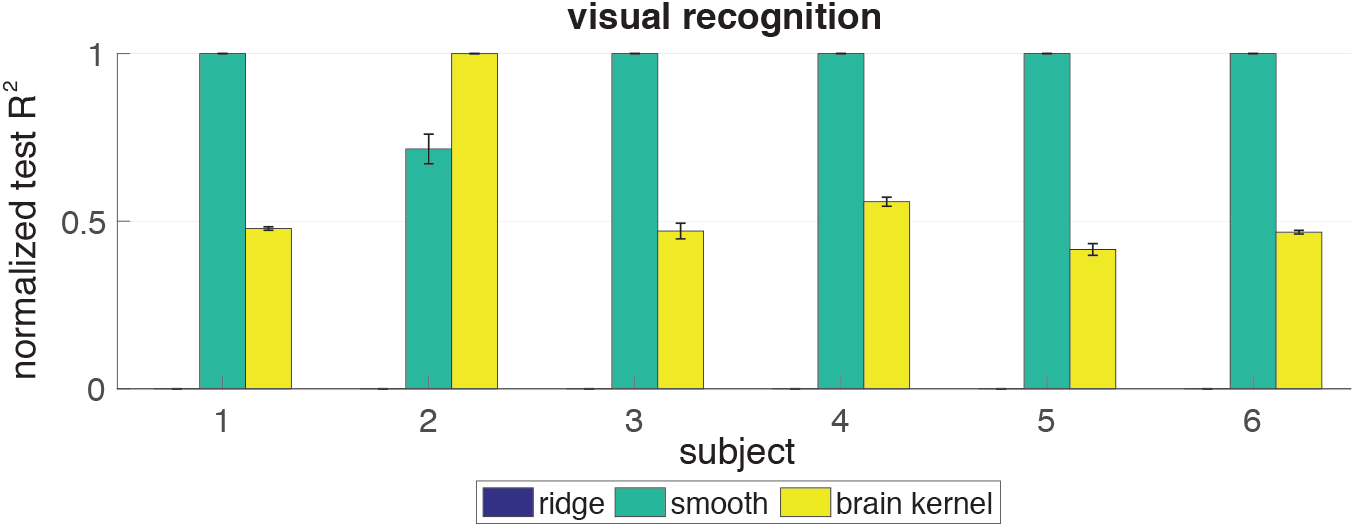
Normalized test *R*^2^ performance on the visual recognition task.

**Fig 31.**
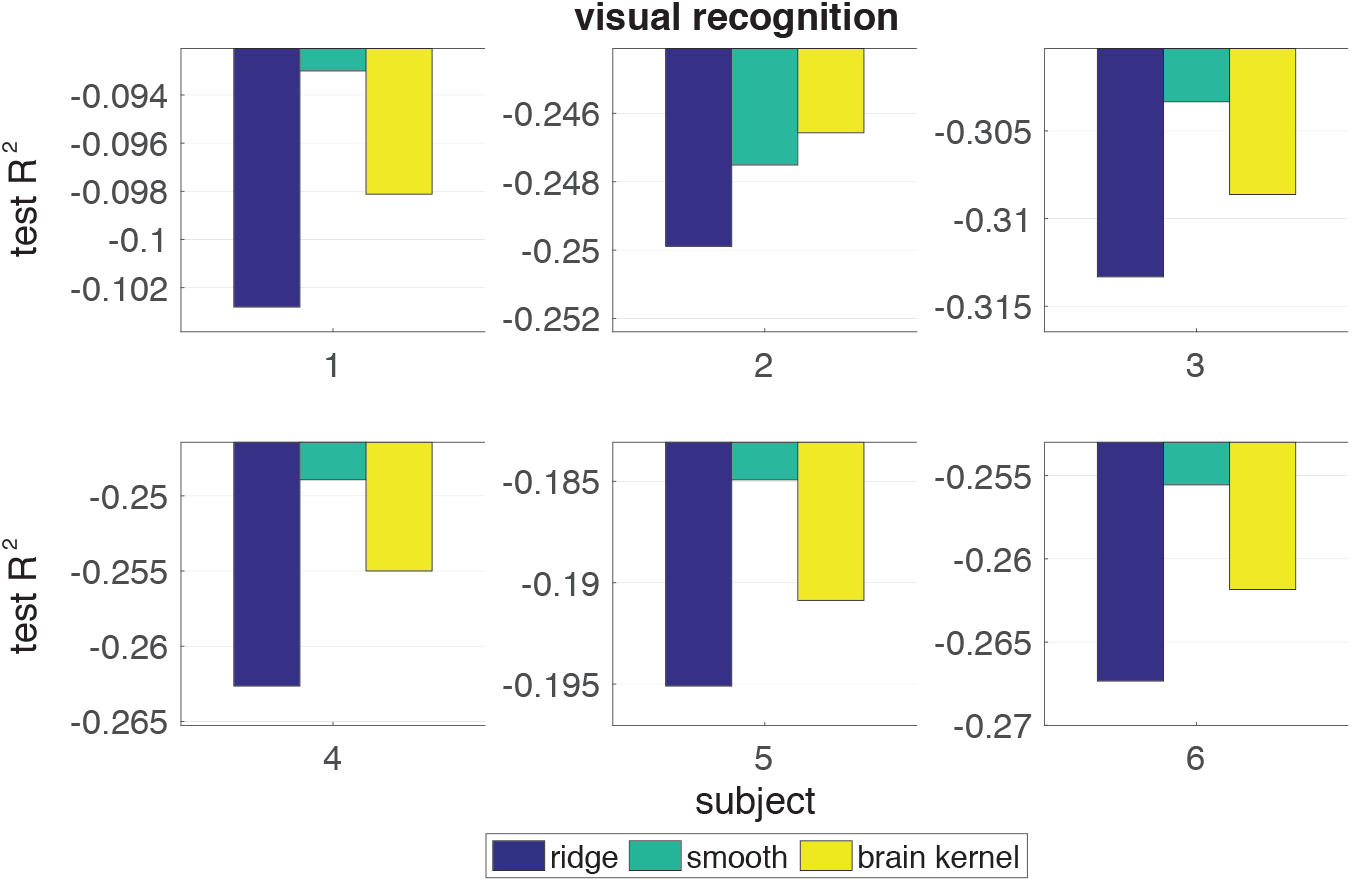
Test *R*^2^ performance on the visual recognition task.

## B.4 BOLD5000 dataset

BOLD5000 is a public fMRI dataset in which subjects viewed 5000 visual images [39]. We used the fMRI datasets collected for 4 subjects with 15 sessions per subject and 333 TRs per session. We extracted predefined functional ROIs [39] from the whole-brain data with 3804 voxels.

## Decoding

The decoding task was binary classification. For each subject, we used 14 sessions for training and 1 session for test. Thus the final accuracy was an average over 15 runs by treating each session as the test set once (Fig. 32). The binary labels for the decoding task were “living” vs “non-living”, which contained a lot of ambiguities. So the decoding task itself was quite difficult. We observe that the brain kernel did not reliably outperform the ridge prior and the smoothing prior estimates across subjects. Moreover, we observe from Fig. 33 top row that the smoothing prior and the brain kernel prior both converged to the ridge prior after hyperparameter estimation, i.e., the estimated length-scale was very small and all three priors had the same performance. Therefore, smoothness did not help the classification task.

**Fig 32.**
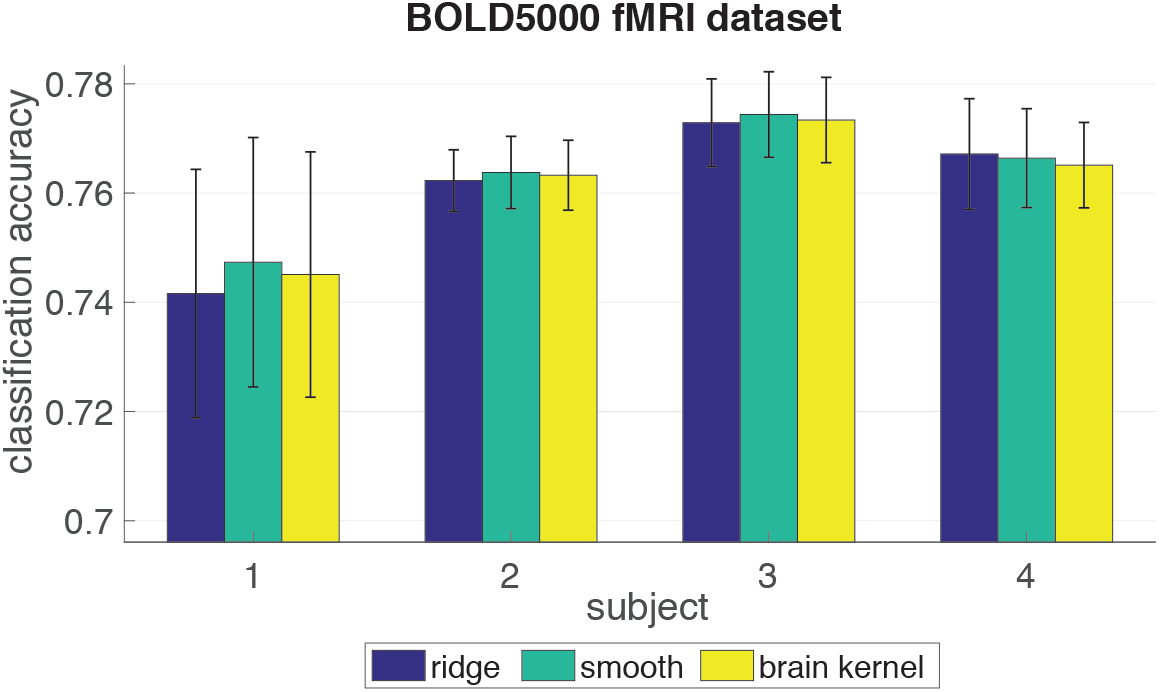
Accuracy performance on the BOLD5000 dataset.

**Fig 33.**
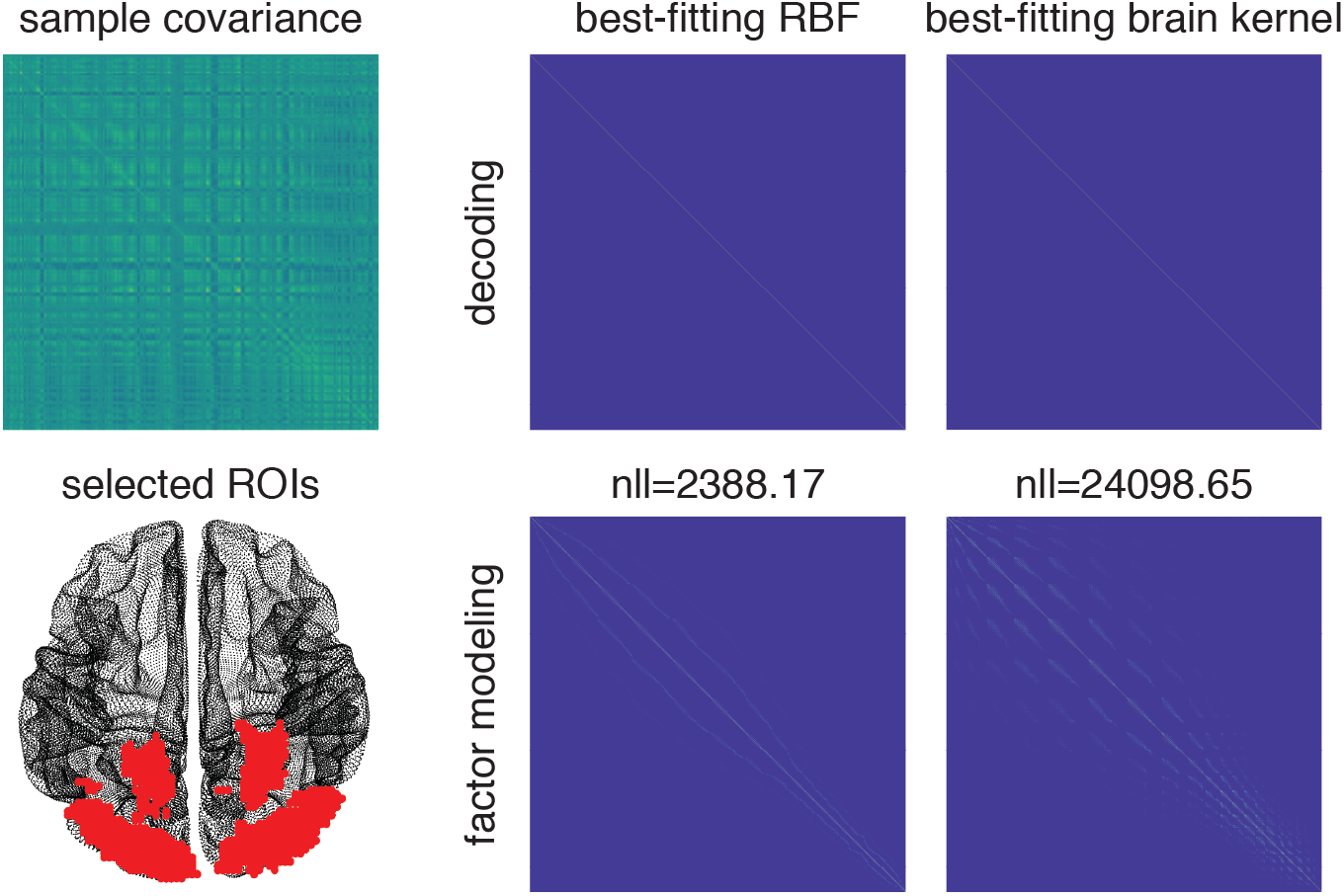
Best-fitting RBF and brain kernel for the BOLD5000 dataset.

## Factor modeling

When evaluating the performance using “co-smoothing”, we split the 15 sessions into halves within each subject, one for training and one for inference and test as described in the main paper. We report the normalized test *R*^2^ (Fig. 34) and the standard test *R*^2^ (Fig. 35) for the three priors for each subject. For all subjects, the brain kernel didn’t show a better performance over either the ridge prior or the smooth RBF kernel. All *R*^2^ values are very small, similar to the visual recognition dataset. We inspect the optimal prior covariance in Fig. 33 bottom row. Both have very sparse patterns, close to the ridge prior. Neither of them captured the statistical structures in the sample covariance. Quantitatively, the NLL value of the brain kernel is larger, implying that the brain kernel served as a worse approximation to the sample covariance compared with the smoothing prior.

**Fig 34.**
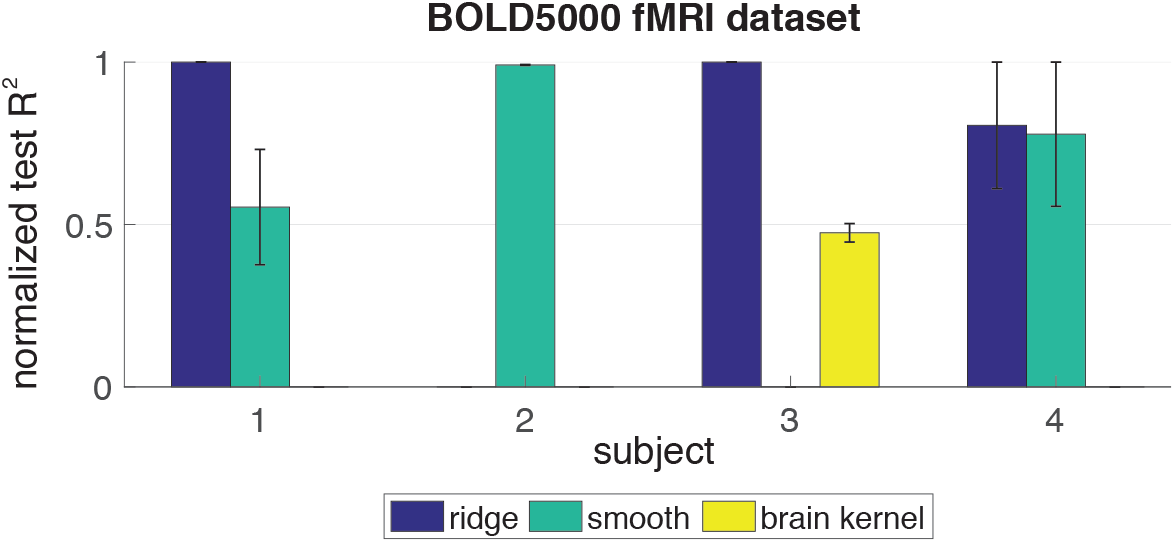
Normalized test *R*^2^ performance on the BOLD5000 dataset.

**Fig 35.**
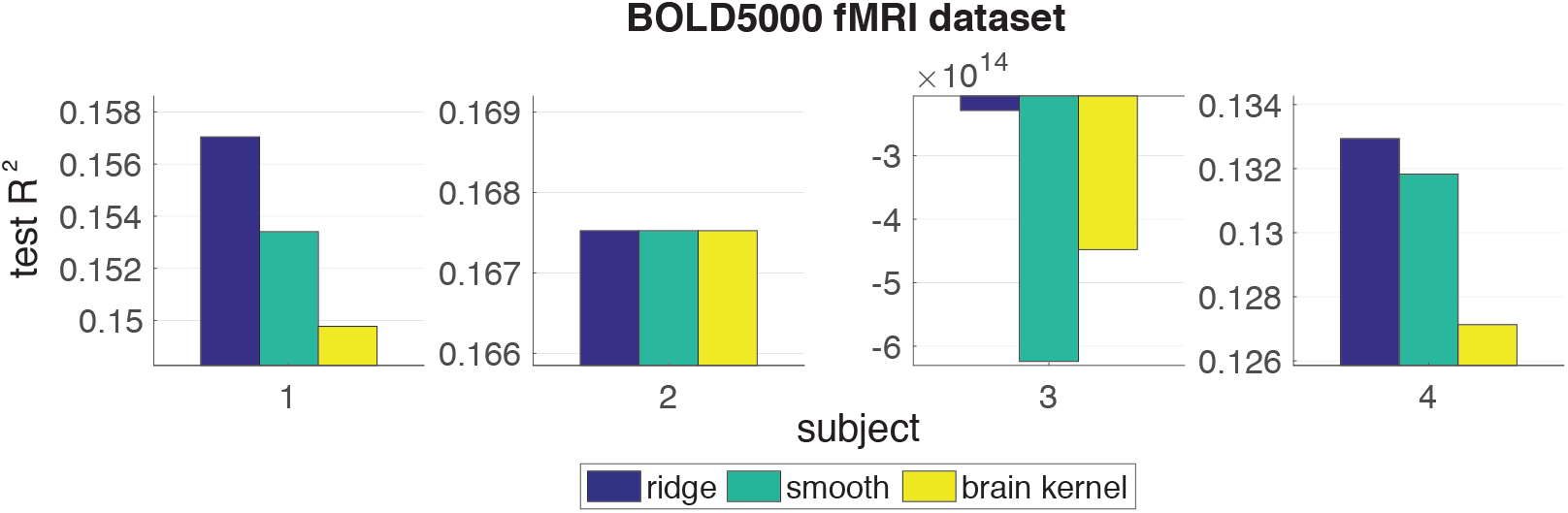
Test *R*^2^ performance on the BOLD5000 dataset.

## C Brain kernel with Matern-3/2

To examine other choices of covariance function for parametrizing the brain kernel, we re-fit the brain kernel to resting-state fMRI data using a Matern-3/2 covariance function. Fig. 36 below shows the covariance matrices obtained from the two resulting brain kernels (RBF and Matern-3/2), along with the sample covariance, for the ROIs used in Fig. 4 in the main paper. We found that the brain kernel with RBF covariance function achieved higher training log-likelihood (lower negative log-likelihood, “NLL”) than the Matern-3/2 covariance function, with negative log-likelihoods of 11250 and and 11252.6, respectively. Thus, changing to a Matern-3/2 parametrization did not improve performance.

**Fig 36.**
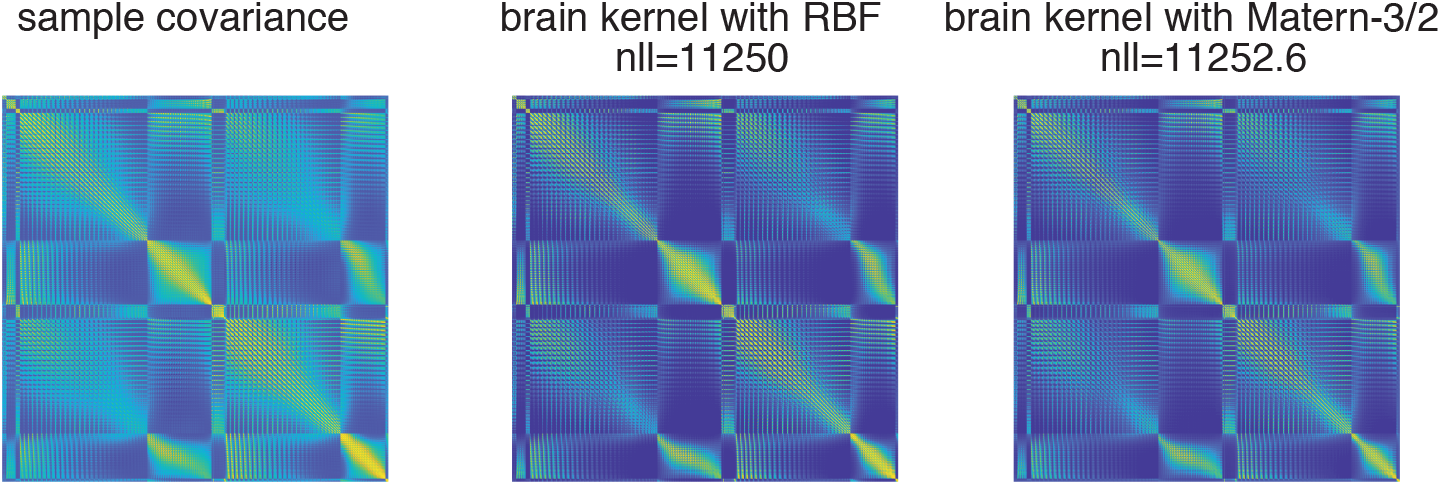
Comparison of the brain kernels fitted with RBF and Matern-3/2 against the sample covariance.

We also compared the RBF and Matern-3/2 brain kernels using the working-memory decoding task fMRI data shown in the main paper. Results are shown below in Fig. 37.

**Fig 37.**
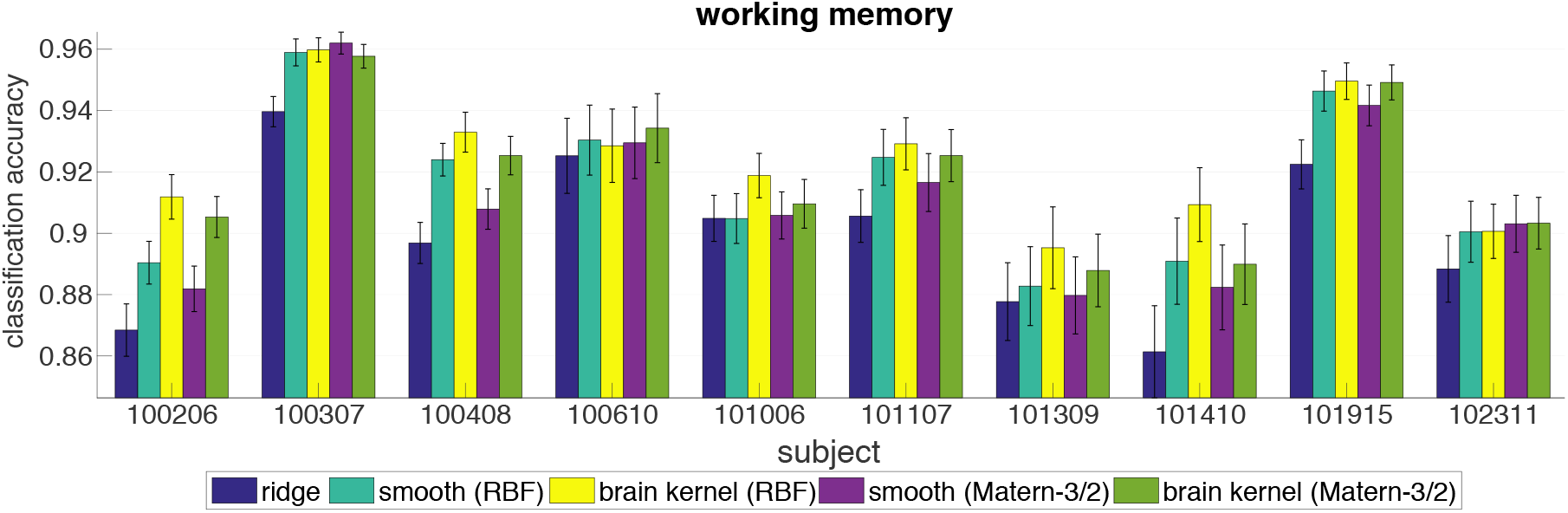
Classification accuracy in the working memory task using regularization under: 1) ridge prior; 2) smoothing (RBF) prior; 3) RBF brain kernel; 4) smoothing (Matern-3/2) prior; and 5) Matern-3/2 brain kernel. In general, the classifiers regularized with RBF covariance outperformed those with Matern-3/2 covariance.

Thus, we found that the RBF covariance function outperformed the Matern-3/2 covariance function in both modeling the covariance of resting-state data and decoding of working memory task data. We have therefore decided to keep our focus on the brain kernel parametrized with RBF covariance in the main paper.

## Footnotes

1 https://github.com/SheffieldML/GPmat

2 https://github.com/poldracklab/fmriprep

3 https://github.com/poldracklab/fmriprep

## References

1. Schäfer J, Strimmer K, et al. A shrinkage approach to large-scale covariance matrix estimation and implications for functional genomics. Statistical applications in genetics and molecular biology. 2005;4(1):32.

2. Bickel PJ, Levina E. Regularized estimation of large covariance matrices. The Annals of Statistics. 2008; p. 199–227.

3. Hsieh CJ, Sustik MA, Dhillon IS, Ravikumar PK, Poldrack R. BIG & QUIC: Sparse inverse covariance estimation for a million variables. In: NIPS; 2013. p. 3165–3173.

4. Treister E, Turek JS. A block-coordinate descent approach for large-scale sparse inverse covariance estimation. In: NIPS; 2014. p. 927–935.

5. Varoquaux G, Gramfort A, Poline JB, Thirion B. Brain covariance selection: better individual functional connectivity models using population prior. In: NIPS; 2010. p. 2334–2342.

6. Van Essen DC, Smith SM, Barch DM, Behrens TE, Yacoub E, Ugurbil K, et al. The WU-Minn human connectome project: an overview. Neuroimage. 2013;80:62–79.

7. Brett M, Johnsrude IS, Owen AM. The problem of functional localization in the human brain. Nature reviews neuroscience. 2002;3(3):243.

8. Stein ML. Interpolation of spatial data: some theory for kriging. Springer Science & Business Media; 2012.

9. Rasmussen CE. Gaussian processes in machine learning. In: Advanced lectures on machine learning. Springer; 2004. p. 63–71.

10. Kitterle FL, Kaye RS. Hemispheric symmetry in contrast and orientation sensitivity. Attention, Perception, & Psychophysics. 1985;37(5):391–396.

11. Di Lollo V. Hemispheric symmetry in duration of visible persistence. Perception & Psychophysics. 1981;29(1):21–25.

12. Westcott M. Hemispheric symmetry of the EEG during the Transcendental Meditation technique. Department of Psychology, University of Durham, Durham, England. 1973;.

13. Fan J, Fan Y, Lv J. High dimensional covariance matrix estimation using a factor model. Journal of Econometrics. 2008;147(1):186–197.

14. Fan J, Liao Y, Liu H. An overview of the estimation of large covariance and precision matrices. The Econometrics Journal. 2016;19(1):C1–C32.

15. Lawrence ND. Learning for Larger Datasets with the Gaussian Process Latent Variable Model. In: AISTATS. vol. 11; 2007. p. 243–250.

16. Damianou AC, Titsias MK, Lawrence ND. Variational inference for uncertainty on the inputs of gaussian process models. arXiv preprint arXiv:14092287. 2014;.

17. Hensman J, Fusi N, Lawrence ND. Gaussian processes for big data. arXiv preprint arXiv:13096835. 2013;.

18. Wu TT, Lange K. Coordinate descent algorithms for lasso penalized regression. The Annals of Applied Statistics. 2008; p. 224–244.

19. Wang H. Coordinate descent algorithm for covariance graphical lasso. Statistics and Computing. 2014;24(4):521–529.

20. Tseng P, Yun S. A coordinate gradient descent method for nonsmooth separable minimization. Mathematical Programming. 2009;117(1):387–423.

21. Fox MD, Snyder AZ, Zacks JM, Raichle ME. Coherent spontaneous activity accounts for trial-to-trial variability in human evoked brain responses. Nature neuroscience. 2006;9(1):23–25.

22. Smith SM, Fox PT, Miller KL, Glahn DC, Fox PM, Mackay CE, et al. Correspondence of the brain’s functional architecture during activation and rest. Proceedings of the National Academy of Sciences. 2009;106(31):13040–13045.

23. Cole MW, Ito T, Bassett DS, Schultz DH. Activity flow over resting-state networks shapes cognitive task activations. Nature Neuroscience. 2016;.

24. WU-Minn H. 1200 subjects data release reference manual; 2017.

25. Delgado MR, Nystrom LE, Fissell C, Noll D, Fiez JA. Tracking the hemodynamic responses to reward and punishment in the striatum. Journal of neurophysiology. 2000;84(6):3072–3077.

26. Binder JR, Gross WL, Allendorfer JB, Bonilha L, Chapin J, Edwards JC, et al. Mapping anterior temporal lobe language areas with fMRI: a multicenter normative study. Neuroimage. 2011;54(2):1465–1475.

27. Buckner RL, Krienen FM, Castellanos A, Diaz JC, Yeo BT. The organization of the human cerebellum estimated by intrinsic functional connectivity. Journal of neurophysiology. 2011;.

28. Hariri AR, Brown SM, Williamson DE, Flory JD, De Wit H, Manuck SB. Preference for immediate over delayed rewards is associated with magnitude of ventral striatal activity. Journal of Neuroscience. 2006;26(51):13213–13217.

29. Smith R, Keramatian K, Christoff K. Localizing the rostrolateral prefrontal cortex at the individual level. Neuroimage. 2007;36(4):1387–1396.

30. Barch DM, Burgess GC, Harms MP, Petersen SE, Schlaggar BL, Corbetta M, et al. Function in the human connectome: task-fMRI and individual differences in behavior. Neuroimage. 2013;80:169–189.

31. Haxby JV, Gobbini MI, Furey ML, Ishai A, Schouten JL, Pietrini P. Distributed and overlapping representations of faces and objects in ventral temporal cortex. Science. 2001;293(5539):2425–2430.

32. Esteban O, Blair R, Markiewicz CJ, Berleant SL, Moodie C, Ma F, et al.. poldracklab/fmriprep: 1.0.0-rc5; 2017. Available from: https://doi.org/10.5281/zenodo.996169.

33. Abraham A, Pedregosa F, Eickenberg M, Gervais P, Mueller A, Kossaifi J, et al. Machine learning for neuroimaging with scikit-learn. Frontiers in neuroinformatics. 2014;8:14.

34. Chen J, Leong YC, Honey CJ, Yong CH, Norman KA, Hasson U. Shared memories reveal shared structure in neural activity across individuals. Nature neuroscience. 2017;20(1):115.

35. Simony E, Honey CJ, Chen J, Lositsky O, Yeshurun Y, Wiesel A, et al. Dynamic reconfiguration of the default mode network during narrative comprehension. Nature communications. 2016;7:12141.

36. Gershman SJ, Blei DM, Pereira F, Norman KA. A topographic latent source model for fMRI data. NeuroImage. 2011;57(1):89–100.

37. Manning JR, Ranganath R, Norman KA, Blei DM. Topographic factor analysis: a bayesian model for inferring brain networks from neural data. PloS one. 2014;9(5):e94914.

38. Wu A, Pashkovski S, Datta SR, Pillow JW. Learning a latent manifold of odor representations from neural responses in piriform cortex. In: Advances in Neural Information Processing Systems; 2018. p. 5378–5388.

39. Chang N, Pyles JA, Marcus A, Gupta A, Tarr MJ, Aminoff EM. BOLD5000, a public fMRI dataset while viewing 5000 visual images. Scientific data. 2019;6(1):49.

40. Grosenick L, Klingenberg B, Katovich K, Knutson B, Taylor JE. Interpretable whole-brain prediction analysis with GraphNet. NeuroImage. 2013;72:304–321.

41. Rao N, Cox C, Nowak R, Rogers TT. Sparse overlapping sets lasso for multitask learning and its application to fmri analysis. Advances in neural information processing systems. 2013;26:2202–2210.

42. Wu A, Park M, Koyejo OO, Pillow JW. Sparse Bayesian structure learning with dependent relevance determination priors. In: Ghahramani Z, Welling M, Cortes C, Lawrence ND, Weinberger KQ, editors. Advances in Neural Information Processing Systems 27. Curran Associates, Inc.; 2014. p. 1628–1636. Available from: http://papers.nips.cc/paper/5233-sparse-bayesian-structure-learning-with-dependent-relevance-determination-priors.pdf.

43. Wu A, Koyejo O, Pillow J. Dependent relevance determination for smooth and structured sparse regression. Journal of Machine Learning Research. 2019;20(89):1–43.

44. Xu H, Lorbert A, Ramadge PJ, Guntupalli JS, Haxby JV. Regularized hyperalignment of multi-set fMRI data. In: 2012 IEEE Statistical Signal Processing Workshop (SSP). IEEE; 2012. p. 229–232.

45. Haxby JV, Guntupalli JS, Nastase SA, Feilong M. Hyperalignment: Modeling shared information encoded in idiosyncratic cortical topographies. ELife. 2020;9:e56601.

46. Lawrence ND. Gaussian Process Latent Variable Models for Visualisation of High Dimensional Data. In: Nips. vol. 2; 2003. p. 5.

47. Nocedal J. Updating quasi-Newton matrices with limited storage. Mathematics of computation. 1980;35(151):773–782.

48. Belkin M, Niyogi P. Laplacian eigenmaps for dimensionality reduction and data representation. Neural computation. 2003;15(6):1373–1396.

49. John J, Draper N. An alternative family of transformations. Applied Statistics. 1980; p. 190–197.

50. Lázaro-Gredilla M, Quiñonero-Candela J, Rasmussen CE, Figueiras-Vidal AR. Sparse spectrum Gaussian process regression. The Journal of Machine Learning Research. 2010;11:1865–1881.

51. Wu A, Aoi MC, Pillow JW. Exploiting gradients and Hessians in Bayesian optimization and Bayesian quadrature. arXiv preprint arXiv:170400060. 2017;.

52. Tipping ME, Bishop CM. Probabilistic principal component analysis. Journal of the Royal Statistical Society: Series B (Statistical Methodology). 1999;61(3):611–622.

53. Lawrence ND. Gaussian process latent variable models for visualisation of high dimensional data. In: Advances in neural information processing systems; 2004. p. 329–336.

54. Lawrence N, Hyvärinen A. Probabilistic non-linear principal component analysis with Gaussian process latent variable models. Journal of machine learning research. 2005;6(11).

55. Damianou A, Lawrence N. Deep gaussian processes. In: Artificial Intelligence and Statistics; 2013. p. 207–215.

56. Vodrahalli K, Chen PH, Liang Y, Baldassano C, Chen J, Yong E, et al. Mapping between fMRI responses to movies and their natural language annotations. NeuroImage. 2018;180:223–231.

